# Prognostically favorable immune responses to ovarian cancer are distinguished by self-reactive intra-epithelial plasma cells

**DOI:** 10.1101/2025.05.12.652297

**Authors:** Allyson C. Banville, Céline M. Laumont, Karanvir Singh, Breeze Gladwin, Jaden Dedora, Liam Mitchell, Jen Wang, Alex Miranda, Bridget Mateyko, Elizabeth A. Chavez, Gian Luca Negri, Se Wing Grace Cheng, Julian Smazynski, Katy Milne, Kylee Wright, Malia Lampard, Nicole Gierc, Phineas T. Hamilton, Sandra E. Spencer, Shreena Kalaria, Talen Oostenbroek, Victor Negrea Puskas, Christian Steidl, Gregg B. Morin, Farouk Nathoo, Brad H. Nelson

## Abstract

Tumor-infiltrating B cells (TIL-Bs) are strongly associated with patient survival; however, the underlying mechanisms are poorly understood. Using integrated single-cell and spatial biology approaches, we defined at clonal resolution the molecular phenotypes, tumor reactivity patterns, and microenvironmental locations of TIL-Bs in high-grade serous ovarian cancer (HGSC). Prognostic benefit was associated with a TIL-B-rich tumor microenvironment with marked infiltration of malignant epithelium by plasma cells (PCs). PCs spanned five molecular phenotypes; exhibited high rates of somatic hypermutation and clonal expansion; and expressed predominantly IgG1 antibodies recognizing broadly expressed nuclear, cytoplasmic, and cell surface self-antigens. Many PC-derived antibodies were polyreactive. Self- and poly-reactive TIL-Bs penetrated tumor epithelium and stroma and expressed interferon-stimulated genes, indicating strong *in situ* activation. The self- and poly-reactive nature of TIL-B responses, reminiscent of autoimmune disease, may provide a means for the immune system to combat tumor heterogeneity and could potentially be harnessed for more effective immunotherapy.

## INTRODUCTION

High-grade serous tubulo-ovarian cancer (HGSC) is one of several cancer types in which tumor-infiltrating lymphocytes (TILs) are strongly associated with patient survival^1–7^ yet today’s immunotherapies have limited efficacy^8–12^. This could partly reflect the fact that HGSC has a relatively low neoantigen load^13–15^, extensive copy number variation^15–17^, and pervasive intra-tumoral heterogeneity that persists from diagnosis to end-stage disease^18–20^.

To develop more effective immunotherapies for HGSC and related malignancies, it is essential to understand the mechanisms by which TILs promote patient survival despite these unfavorable tumor features. We^21–23^ and others^24,25^ have shown that the most immunologically “hot” HGSC tumors contain not only tumor-infiltrating T cells (TIL-Ts) but also B cells (BCs) and their differentiated counterparts plasma cells (PCs), collectively referred to as tumor-infiltrating B cells (TIL-Bs). The combined presence of TIL-Bs and TIL-Ts at diagnosis is among the strongest predictors of long-term survival in HGSC.^26,27^ Furthermore, TIL-Bs show prognostic benefit in a wide range of other cancers and are associated with response to immune checkpoint inhibition (ICI).^28–30^

TIL-Bs are widely thought to promote anti-tumor immunity through the production of antibodies to tumor-associated antigens. Such antibodies could potentially disrupt the expression or function of important tumor proteins, as demonstrated in anecdotal reports in HGSC^31–33^ and lung cancer^34^. Theoretically, antibodies could also flag tumor cells for destruction through complement-dependent cytotoxicity (CDC), antibody-dependent cellular cytotoxicity (ADCC) and antibody-dependent cellular phagocytosis (ADCP). Apart from antibodies, TIL-Bs could potentially deploy cell-based mechanisms, including serving as antigen-presenting cells (APCs) to CD4 and CD8 T cells^35^ and producing immunostimulatory cytokines^36,37^. Recent large-scale single-cell RNA sequencing (scRNA-seq) studies of human cancers have identified several molecular subsets of TIL-Bs as well as their clonal architecture as revealed by B Cell Receptor (BCR) sequences.^34,38–41^ In general, these studies have identified TIL-B phenotypes that could potentially mediate any of the above functions. However, there is very limited data linking TIL-B phenotypes to tumor reactivity, which is critical to determining which of these possible anti-tumor functions is active in human tumors and should be prioritized for immunotherapeutic development.

We report here the first systematic and linked analysis of TIL-B phenotypes, clonal architecture, tumor reactivity, and spatial locations in human cancer. Applying single-cell, spatial, and recombinant antibody-based approaches to tumors with prognostically favorable TIL-B patterns, we found substantial infiltration by clonally expanded, self-reactive PCs with highly activated phenotypes. This suggests that TIL-Bs use autoimmune-like mechanisms to respond to tumors, which could provide an effective means to combat the intra-tumoral heterogeneity that characterizes advanced disease. Our findings highlight new opportunities and challenges for safe and effective B-cell-directed immunotherapies for HGSC and other cancers.

## RESULTS

### HGSC patient survival is associated with infiltration of tumor epithelium by BCs and PCs

We applied multi-color immunofluorescence (mcIF) to a large HGSC cohort (cohort A, N = 981, **Supplemental Table S1**) to measure the co-occurrence patterns and prognostic significance of TIL-Bs, TIL-Ts, and tumor-associated macrophages (TAMs), as well as the programmed death 1 (PD-1) and PD-1 ligand 1(PD-L1) pathway. BCs and PCs were detected at average densities of 34.8 and 78.9 cells/mm^2^ respectively, compared to average densities of 69.3 and 88.5 cells/mm^2^ for CD8 and CD4 T cells, respectively. The majority (75%) of TIL-B-positive tumors contained more PCs than BCs (referred to as PC-rich tumors), whereas BCs outnumbered PCs in 23% of tumors (BC-rich tumors) (**Figure 1A** and **Supplemental Table S2**). Unsupervised clustering based on BC and PC densities revealed four groups of tumors, which we designated BME1 to BME4 to signify “B cell-defined microenvironment” (**Figure 1A,B**). BME1 (31.7% of tumors) had negligible BCs or PCs. BME2 (29.0%) had stromal PCs with negligible BCs, whereas BME3 (15.6%) had both stromal PCs and BCs. BME4 (23.8%) was distinguished by the infiltration of both stroma and epithelium by PCs and BCs. In particular, the density of epithelial PCs increased 11.6-fold between BME3 and BME4, exceeding the increases seen for epithelial BCs (6.2-fold), CD8+ T cells (3.2-fold) and PD-L1+ macrophages (2.0-fold) (**Figure 1C**, **Supplemental Table S2** and **Supplemental Figure S1A**). Accordingly, the majority (80%) of BME4 tumors were PC-rich. The increased densities of epithelial PCs and BCs in BME4 tumors reflected an increase in their total cell numbers rather than a preferential shift from stroma to epithelium, as the proportion of epithelial BCs and PCs (relative to total BCs and PCs) was similar in BME4 and BME3 tumors (**Figure 1C**, **Supplemental Figure S1A** and **Supplemental Table S3**).

**Figure 1.**
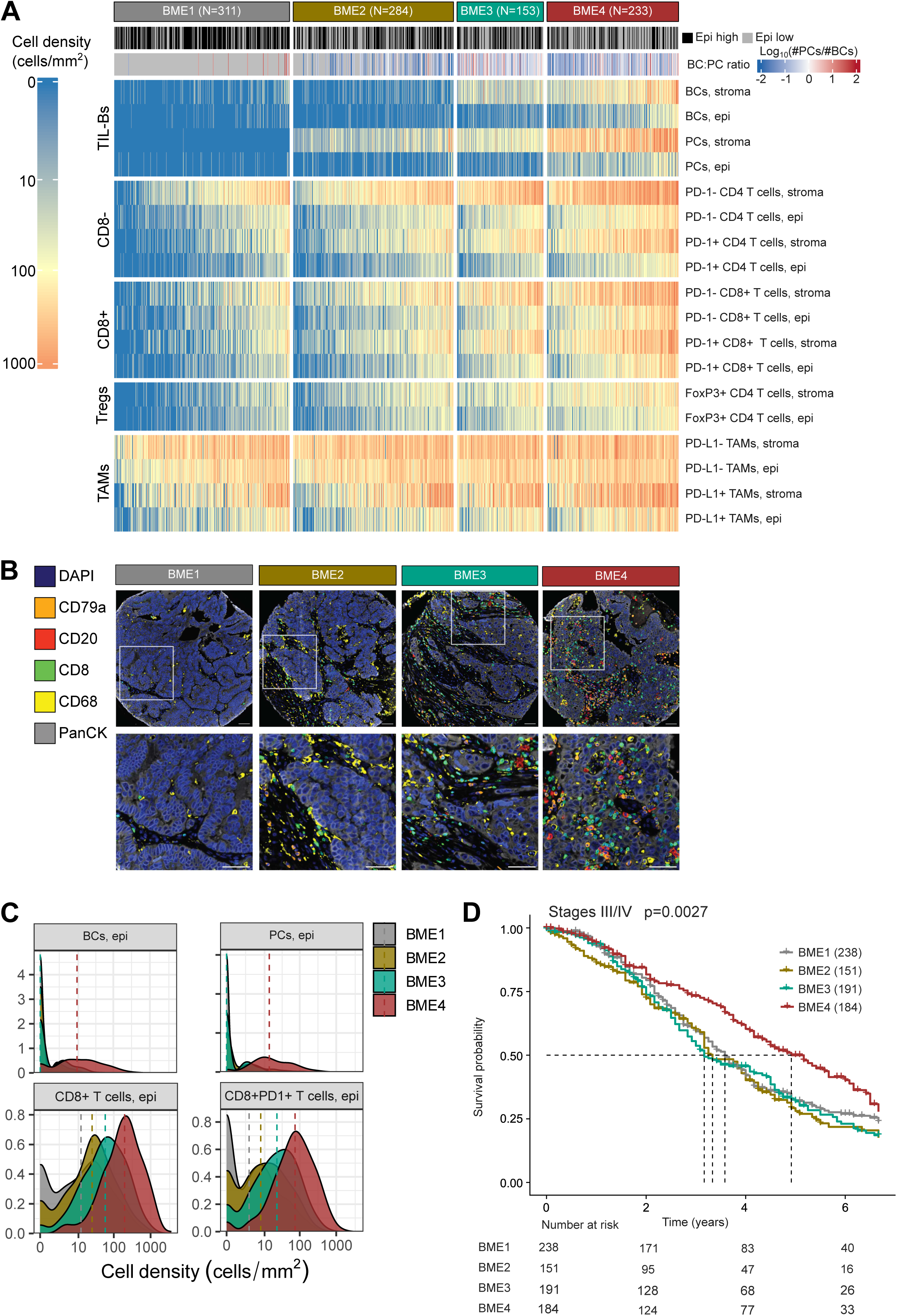
Multicolor IF (mcIF) analysis of HGSC patient tumors reveals four distinct B-cell-defined environments (BMEs) corresponding to distinct survival profiles. **(A)** Heatmap depicting cell densities (cells/mm^2^) of the indicated tumor-infiltrating immune cells across the four BMEs defined by unsupervised clustering based on TIL-B content. Epi-low = <50% epithelium. Epi-high = ≥50% epithelium. **(B)** Example mcIF images. Scale bar: 50 µm. **(C)** Epithelial B cells (BCs), plasma cells (PCs), and CD8+ T cell densities by BME type. Y-axes represent probability densities. The vertical dashed lines indicate the median. **(D)** Kaplan-Meier survival plot by BME type. Cohort A (N=981). Epi: Epithelium. TAM: Tumor-associated macrophage. Treg: T regulatory cell.

Cox proportional hazards modelling of BME2 to BME4 revealed that BME4 alone was associated with a significantly lower hazard of death compared to BME1 (**Supplemental Table S4**). Likewise, by Kaplan-Meier analysis, BME4 was associated with significantly higher overall survival compared to all other BMEs (**Supplemental Figure S1B**), an effect that was most pronounced in stage III/IV cases (**Figure 1D**) and tumors with high epithelial content, in line with our prior report^27^ (**Supplemental Tables S4** and **S5** and **Supplemental Figure S1C**). Thus, the prognostic benefit of TILs was exclusive to BME4 tumors, which were distinguished by a high density of epithelial TIL-Bs, in particular PCs.

### TIL-B phenotypes, class switching, somatic hypermutation, and clonal expansion patterns

We used scRNA-seq with matched single-cell gene expression (scGEX) and B cell receptor (scBCR) sequencing to investigate the transcriptional profiles and clonal composition of TIL-Bs from 14 primary (untreated) HGSC cases (cohort B, **Figure 2A** and **Supplemental Tables S6**), which by mcIF showed a representative range of TIL-B and TIL-T densities (data not shown). In 5/14 (36%) cases, the immune infiltrate was dominated by myeloid cells (75-89% of CD45+ cell infiltrate), whereas the remaining 9/14 tumors were dominated by lymphocytes (48-95% of CD45+ cell infiltrate) (**Figure 2A, Supplemental Figure S2** and **Supplemental Table S7**), as seen in other cancers^42^. Of the nine lymphocyte-dominated tumors, seven had substantial TIL-Bs (five PC-rich and two BC-rich) (**Figure 2B** and **Supplemental Table S7**).

**Figure 2.**
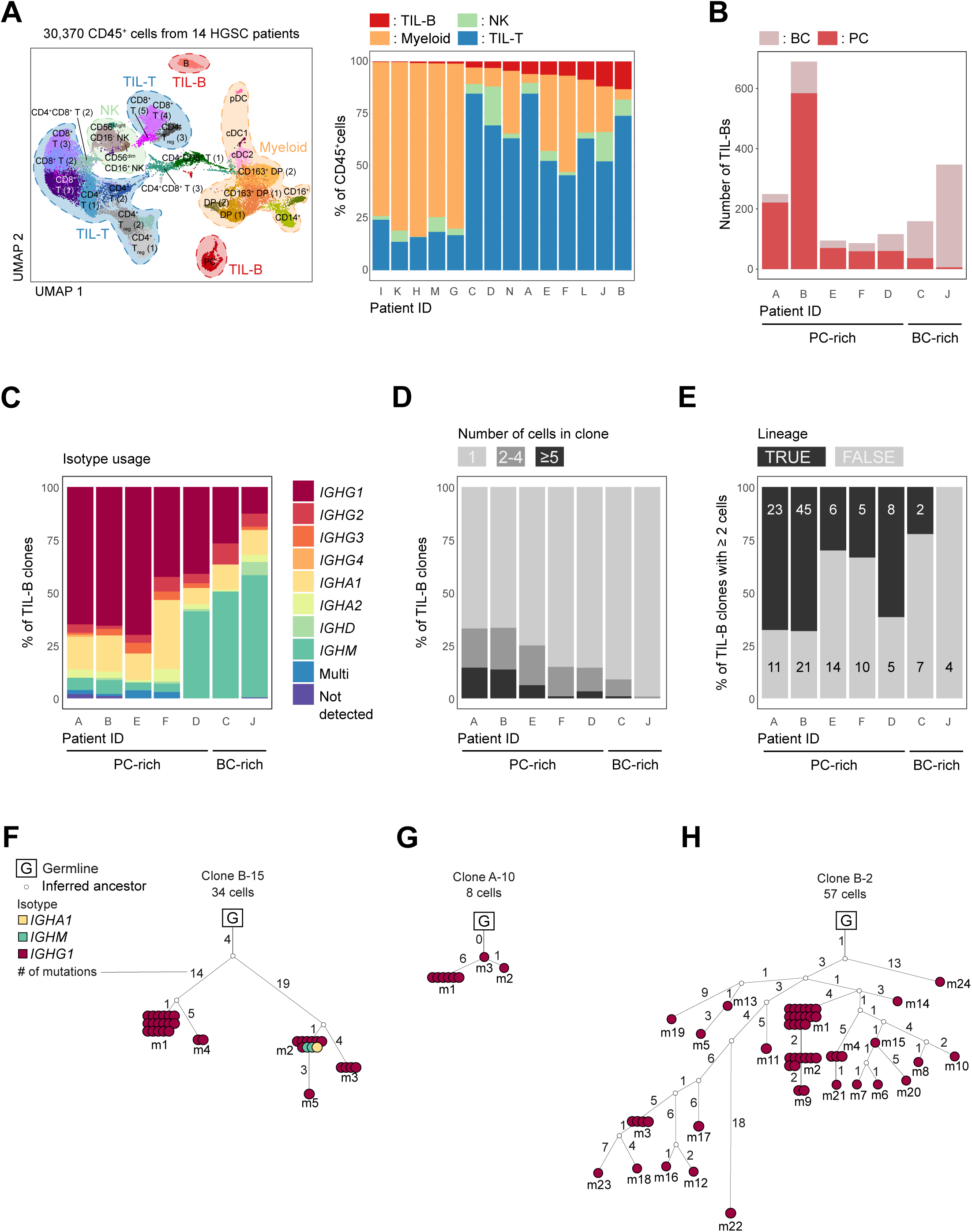
Single-cell gene expression (scGEX) and B cell receptor (scBCR) sequencing reveals TIL-B phenotypes, class switching, somatic hypermutation, and clonal expansion patterns. **(A)** Left: UMAP clustering of all cells in the scGEX dataset. Overlaid cell phenotypes are based on canonical marker genes (**Supplemental Figure S2**). Right: Proportions of main immune cell phenotypes in scGEX dataset by patient. **(B-E)** TIL-B characteristics from tumors with 20 or more TIL-Bs. **(B)** Number of B cells (BCs) and plasma cells (PCs) per patient. **(C)** Isotype usage by patient. “Multi” indicates clones comprising two or more isotypes. **(D)** Clonal expansion by patient. **(E)** Proportion of expanded TIL-B clones with (TRUE) or without (FALSE) lineages by patient. Overlays indicate the number of clones. **(F-H**) Example TIL-B clonal lineages. Mutant (m) number identifies individual clonotypes. Number of circles = number of cells. Numbers on branches indicate the total number of nucleotide mutations between relevant clonotypes. Cohort B (N=14). HGSC: High-grade serous ovarian cancer. NK: Natural killer. TIL-B: Tumor-infiltrating B lymphocyte. TIL-T: Tumor-infiltrating T lymphocyte.

The scGEX data revealed three BC and five PC phenotypic clusters (**Supplemental Figure S3A**). A sixth PC cluster appeared to comprise “doublets” involving T cells and NK cells (as seen in other scRNA-seq studies^43^), despite the measures we took to remove such artefacts. The BC clusters comprised naïve-like, memory-like, and activated BCs (**Supplemental Figure S3B**). Similar to other cancer scRNA-seq datasets^34,38–41^, naïve-like BCs (27.8% of BCs) were characterized by the expression of *IGHM*, *IGHD*, *TCL1A, CD72*, and *FCER2*; memory-like BCs (34.5% of BCs) by expression of *TNFRSF13B*, *AIM2* and *CD27*; and activated BCs (34.3% of BCs) by expression of *CD69*, *CD83* and *NR4A2*. Most naïve-like (100%) and activated (71.5%) BCs were found in BC-rich tumors, while most memory-like BCs (70.4%) were in PC-rich tumors. BCs demonstrated relatively high scores for gene modules associated with antigen presentation (**Supplementary Figure S4A**), and memory-like BCs in particular expressed cytokines such as *TGFB1*, *LTA*, and *LTB* (**Supplemental Figure S4B**).

The five PC clusters comprised **(*i*)** plasmablasts (PBs), **(*ii*)** IgG+ PCs, **(*iii*)** interferon-stimulated gene (ISG)-negative PCs and **(*iv*)** ISG-positive PCs, and **(*v*)** MCL1+ PCs (**Supplemental Figure S3C,D**). The PB cluster expressed HLA and proliferative genes characteristic of conventional PBs (e.g., *STMN1*, *TUBA1B*, and *HMGN2*).^44^ The IgG1+ PC cluster showed striking expression of *IGHGP, IGHG2, IGHG1*, and *IGKC*, similar to other cancer scRNAseq datasets^40^. ISG- and ISG+ PCs expressed genes related to protein folding, degradation, and secretion. ISG+ PCs additionally expressed higher levels of *IFI6, ISG15,* and *IFITM1*; genes related to signaling and cell cycle; and the pro-apoptotic cytokine TRAIL (*TNFSF10*). The ISG+/- clusters resembled those previously described in cancer as “conventional” and “stressed” PCs^40^; ISG+ PCs also resembled a previously described phenotype over-represented in tumors versus other tissues^45^. The MCL1+ PC cluster expressed the apoptosis-regulating gene *MCL1*^46^ and three genes (*FOSB, JUN,* and *PRDM1*) that are highly expressed by long-lived PCs^44^. Compared to BC clusters, all five PC clusters demonstrated lower scores for gene modules associated with antigen presentation (**Supplemental Figure S4A**) and elevated scores for gene modules associated with antibody secretion^47^ (**Supplemental Figure S3E**), although the IgG+ PC cluster had among the lowest scores for PCs, suggesting possible uncoupling of transcription and translation. In contrast to BCs, PCs did not express appreciable levels of *TGFB1*, *LTA* or *LTB* (**Supplemental Figure 4B**).

Cross-referencing the scGEX and scBCR data revealed that, in both BC- and PC-rich tumors, the predominant isotypes expressed by PCs were IgG1 (100% in BC-rich and 77.0% in PC-rich tumors, respectively) and to a lesser extent IgA1 (0% and 8.9%, respectively); in contrast, the majority of BCs expressed IgM (59.8% and 32.4%, respectively) (**Supplemental Table S8**). Accordingly, the dominant isotypes were IgG1 and IgA1 in PC-rich tumors and IgM in BC-rich tumors (**Figure 2C**). The highest degree of somatic hypermutation (SHM) was seen in PC-rich tumors (mean = 15.9 and 16.4 mutations among BCs and PCs, respectively) compared to BC-rich tumors (mean = 10.2 and 7.8 mutations, respectively) (**Supplemental Figure S5A**).

Evaluating the scBCR data, we defined a clonotype as one or more cells expressing identical BCR sequences and a clone as one or more clonotypes expressing related BCRs (see Methods for relational thresholds). Three types of clones were identified: non-expanded clones (one unique BCR observed in one cell, i.e., “singletons”), expanded clones without lineages (one unique BCR observed in multiple cells), and expanded clones with lineages (multiple related BCRs observed in multiple cells). Across all evaluated tumors, 94.9% of BCs comprised non-expanded clones compared to only 47.8% of PCs. All three BC clusters showed similar levels of expansion (mean = 1.0-1.3 cells per clone), yet memory-like BCs had higher levels of mutations compared to naïve-like and activated BCs (mean = 21.1 versus 7.8 and 8.8 mutations per clonotype, respectively, **Supplemental Figure S4B**). PC-rich tumors showed an almost five-fold higher proportion of expanded clones compared to BC-rich tumors (mean = 24.1% versus 4.9%, respectively, **Figure 2D** and **Supplemental Table S7**) and a higher average clone size (mean = 5.9 cells versus 1.1 cells, respectively). PBs, IgG+ PCs, and ISG-/+ PCs showed similar levels of expansion (mean = 3.9-4.3 cells per clone) whereas MCL1+ PCs showed a relatively higher level of expansion (mean = 6.9 cells per clone). IgG+ PCs and ISG-/+ PCs had higher mutation counts (17.1-17.9) than PBs and MCL1+ PCs (12.2 and 11.7 mutations per clonotype, respectively; **Supplemental Figure S5B**). Thus, BCs were predominantly IgM+ non-expanded clones, whereas PCs were mostly IgG1+ with substantially higher degrees of SHM and clonal expansion.

Virtually all (97.8%) expanded clones with lineages were from PC-rich tumors (**Figure 2E** and **Supplemental Table S7**). Of the expanded clones with lineages, 98.1% exhibited SHM compared to 87.5% of expanded clones without lineages and 66.4% of non-expanded clones (**Supplemental FigureS5C**). Moreover, expanded clones with lineages had increased mutation counts (mean = 16.8 mutations) compared to those without lineages (mean = 12.1 mutations) or non-expanded clones (12.8 mutations) (**Figure 2F-H**, **Supplemental Figure S5C**). The vast majority of clonotypes within lineages were class-switched to IgG1 (87.8%) compared to IgA1 (4.8%) or other isotypes (0.6%), and a minor proportion of lineages (6.8%) contained multiple isotypes (typically IgM in combination with IgG1 or IgA1) (**Figure 2F**); a similar distribution was seen for expanded clones without lineages (**Supplemental Table S8**). Thus, PC-rich tumors were distinguished by relatively high levels of SHM, clonal expansion and class switching to IgG1.

### Clonally expanded TIL-Bs recognize intracellular and cell surface antigens expressed by normal and malignant cell types

To investigate the antigen reactivity of TIL-Bs, we made recombinant antibodies (rAbs) corresponding to the 51 most expanded clones (using a threshold of 5 or more observed cells) across the seven TIL-B-positive tumors (**Supplemental Table S9**). With this threshold, 50/51 clones (98%) came from PCs from PC-rich patients (**Figure 2D**) and most (78.4%) were members of lineages. The vast majority (92.2%) were IgG1 (either exclusively or in combination with other isotypes) (**Supplemental Table S9**); to enable direct comparisons of binding properties, all rAbs were made with an IgG1 backbone.

As an initial assessment of antigen reactivity, rAbs were screened by flow cytometry for binding to a panel of four HGSC, three “non-HGSC malignant” and three non-malignant cell lines. By intracellular staining, a large majority (82%) of rAbs recognized at least one HGSC cell line (**Figure 3A,B** and **Supplemental Figure S6A,B**); of these, all but one also recognized non-HGSC and/or non-malignant cell lines. By cell surface staining, 15.7% of rAbs recognized one or more HGSC cell lines, and 19.6% recognized any malignant or non-malignant cell line. The vast majority (90%) of cell surface-reactive rAbs also displayed intracellular reactivity, although not necessarily to the same cell lines. Enzymatic dissociation of OVCAR3 cells prior to staining eliminated the binding of all cell-surface-reactive rAbs, confirming the involvement of cell surface antigens (**Supplemental Figure S6C**). Only 6/51 (11.8%) of rAbs showed cell surface binding to healthy donor-derived peripheral blood mononuclear cells (PBMCs, primarily NK cells and macrophages) (**Figure 3A** and **Supplemental Figure S7**), suggesting that TIL-Bs may preferentially recognize antigens on epithelial versus lymphoid cells.

**Figure 3.**
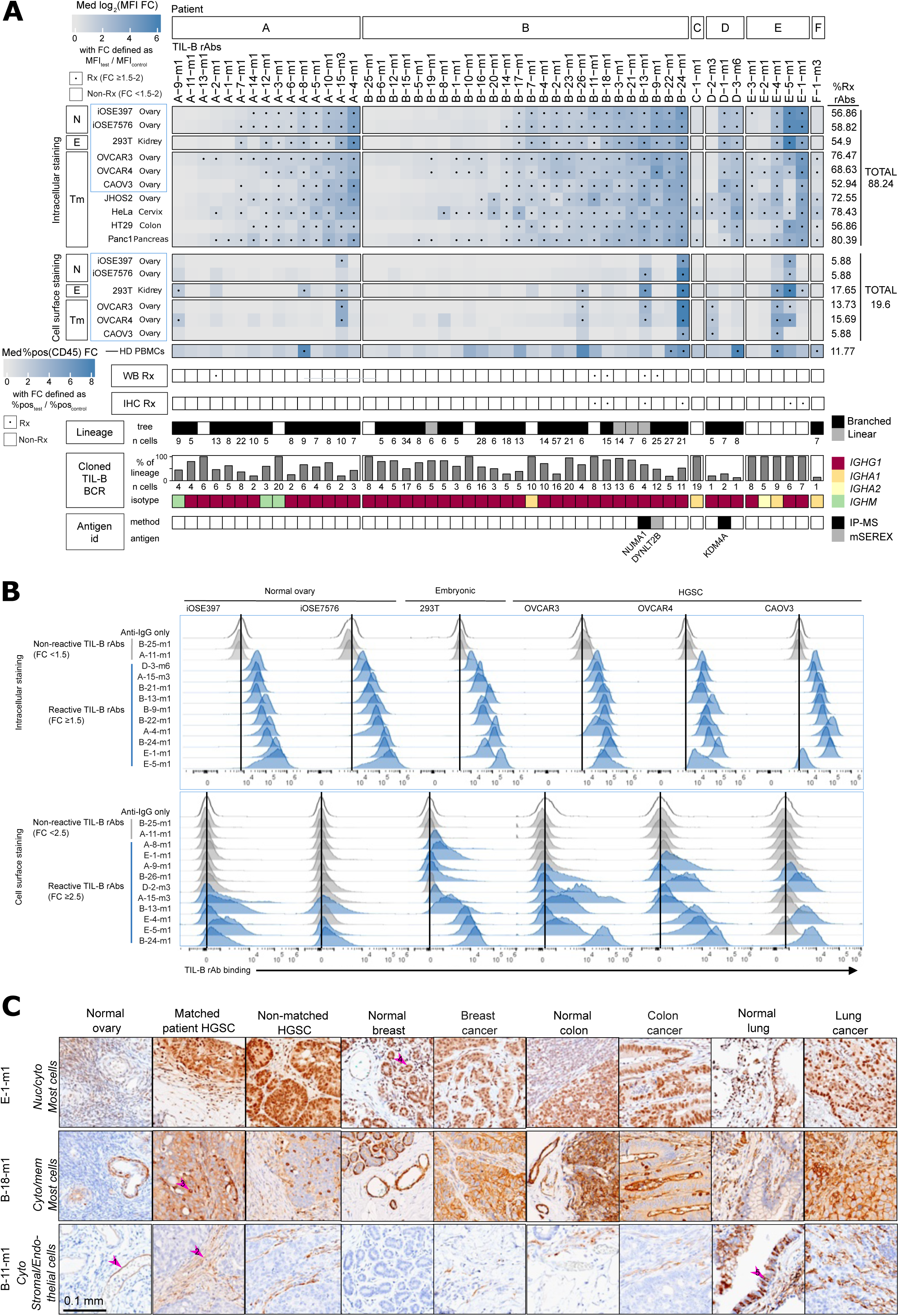
Clonally expanded TIL-Bs recognize intracellular and cell surface antigens expressed by normal and malignant cell types. **(A)** Heatmap summarizing the reactivity of all 51 unique TIL-B rAbs from cohort B relative to other clonal features. Blue boxes outline the core panel of cell lines used for both intracellular and cell surface staining. **(B)** Flow cytometry histograms showing the binding of two non-reactive TIL-B rAbs (grey) and the top 10 most reactive TIL-B rAbs (blue) by intracellular (upper) and cell surface (lower) staining methods using the core panel of cell lines. Vertical lines indicate the baseline signal from the anti-IgG only sample. **(C)** Staining patterns of three IHC-reactive TIL-B rAbs. Arrows indicate (1) endothelial cell cytoplasmic (cyto) staining, (2) stromal cell cytoplasmic staining, (3) epithelial cell membranous (mem) staining, (4) epithelial cell nuclear(nuc)/cytoplasmic staining, and (5) epithelial cell cytoplasmic staining. E: Embryonic. FC: Fold change. IP-MS: Immunoprecipitation mass spectrometry. Med: Median. MFI: Median fluorescence intensity. N: Normal. mSEREX: modified serological analysis of recombinant cDNA expression. Rx: Reactive. Tm: Tumor.

By western blot (WB), 5/37 evaluated rAbs (14%) reacted to at least one cell line (**Figure 3A**, **Supplemental Figure S6B** and **Supplemental Figure S8A**). Multiple bands were detected for 4/5 of these rAbs, suggesting they may be polyreactive. By immunohistochemistry (IHC), 6/51 (11.8%) rAbs showed binding to one or more tissue types (**Figure 3C**, **Supplementary Figure S6B** and **Supplementary Figure S8B**), including patient-matched tumors, other HGSC tumors, non-HGSC tumors (breast, colon, lung), and corresponding normal tissues. All six of these rAbs showed nuclear and/or cytoplasmic staining, and one rAb additionally showed membranous staining. Fifty percent (3/6) of rAbs recognized both epithelial and stromal cells, including what appeared to be endothelial cells in blood vessels; the remaining rAbs appeared selective for either stromal or epithelial cells.

### Identification and characterization of TIL-B target antigens

To identify TIL-B target antigens, we performed immunoprecipitation (IP) experiments with several rAbs that showed cell line recognition by flow cytometry. By silver staining, the IP outputs for two rAbs showed unique banding patterns and therefore were subjected to mass spectrometry (MS) analysis. rAb D-1-m1 uniquely immunoprecipitated a band of approximately 140 kDa (**Figure 4A**), which was identified by MS as lysine-specific demethylase 4a (KDM4a, 120.7 kDa) (**Figure 4B** and **Supplemental Table S10**); KDM4a was validated by both ELISA (**Figure 4C**) and IP-WB (**Figure 4D**, **Supplemental Figure S9A**). rAb B-13-m1 immunoprecipitated multiple bands (**Figure 4E**), similar to the pattern seen by WB (**Supplemental Figure S8A**). MS revealed enrichment of nuclear mitotic apparatus protein 1 (NuMA, 238.1 kDa) (**Figure 4F** and **Supplemental Table S10**), which matched the approximate molecular weight (∼250 kDa) of the most intense band seen by IP-silver stain (**Figure 4E**).

**Figure 4.**
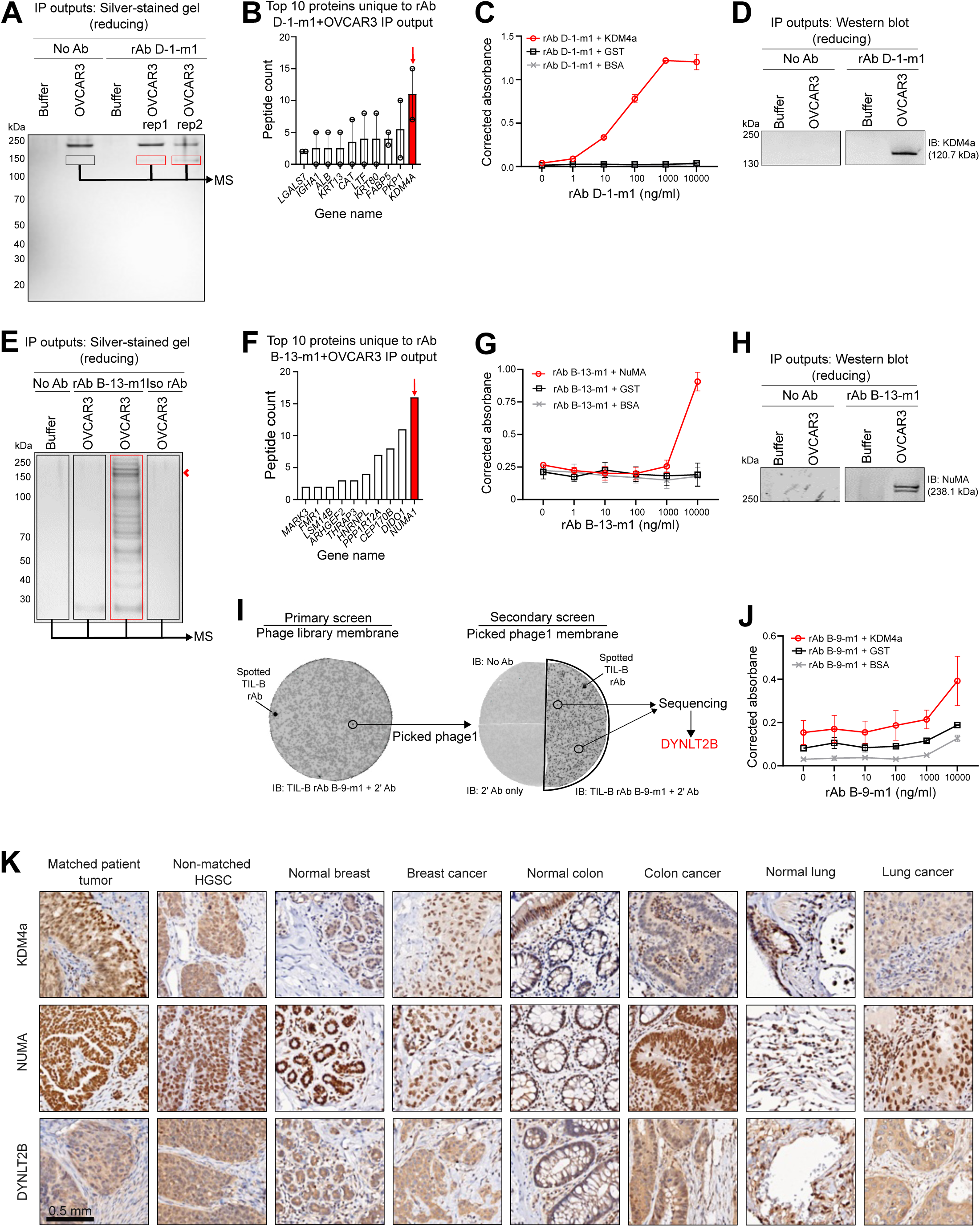
Identification and characterization of TIL-B target antigens. **(A)** Silver-stained (SS) gel of immunoprecipitation (IP) output samples showing unique bands immunoprecipitated by TIL-B rAb D-1-m1 (red boxes) relative to the control (black box). Labels indicate the IP inputs (No Ab versus TIL-B rAb D-1-m1 and buffer versus OVCAR3 lysate) related to each IP output sample analysed on the gel. Gel pieces corresponding to the red/black boxes were excised and analyzed by mass spectrometry (MS). **(B)** Top 10 proteins (ranked by peptide counts) uniquely identified by MS in both replicates of the TIL-B rAb D-1-m1 + OVCAR3 IP output samples relative to controls. **(C)** ELISA with TIL-B rAb D-1-m1 and recombinant KDM4a, GST, and BSA proteins. **(D)** IP-Western blot (WB) with IP output samples from TIL-B rAb D-1-m1 (or no Ab) + OVCAR3 lysate (or buffer) probed with a commercial anti-KDM4a antibody. **(E)** IP-SS gel of 10% of the total IP output samples with TIL-B rAb B-13-m1 (or no Ab or isotype control rAb) and OVCAR3 lysate (or buffer) input. **(F)** Top 10 proteins (ranked by peptide counts) uniquely identified by MS in the TIL-B rAb B-13-m1 + OVCAR3 IP output sample relative to controls. **(G)** ELISA with TIL-B rAb B-13-m1 and recombinant NuMA, GST, and BSA proteins. **(H)** IP-WB with TIL-B rAb B-13-m1 (or no rAb) + OVCAR3 (or buffer) IP output samples probed with a commercial anti-NuMA antibody. **(I)** Discovery of DYNLT2B as the target antigen of TIL-B rAb B-9-m1 by phage-based cDNA immunoscreening (modified SEREX). **(J)** ELISA with TIL-B rAb B-9-m1 and recombinant DYNLT2B, GST, and BSA proteins. **(K)** Examples of immunohistochemistry (IHC) staining on patient-derived tissues with commercial antibodies against all three identified TIL-B target antigens. BSA: Bovine serum albumin. GST: Glutathione S transferase.

NuMA was validated by both ELISA (**Figure 4G**) and IP-WB (**Figure 4H, Supplemental Figure S9A**). In addition to IP-MS, we identified a TIL-B target antigen using a modified version of SEREX^48–50^ which revealed a truncated version of dynein light chain TCTEX-type 2B (DYNLT2B) as a target for rAb B-9-m1 (**Figure 4I)**; by ELISA, B-9-m1 also recognized wild type DYNLT2B, albeit weakly (**Figure 4J**).

By IHC with commercial antibodies, KDM4a, NuMA, and DYNLT2B were expressed by their matched patient tumor samples, as well as various malignant and healthy tissues **(Figure 4K)**, consistent with published expression data^51–53^. Furthermore, all three antigens were expressed at the mRNA level in all healthy tissue and cancer samples in the GTex and TCGA databases, with significant over-expression in multiple cancers, including ovarian cancer in the case of KDM4a and DYNLT2B (**Supplemental Figure S10** and **Supplemental Table S11).**

### Evolution of TIL-B reactivity within clonal lineages

To evaluate how TIL-B reactivity evolves during SHM, we produced rAbs from the inferred germline BCR and two or more descendent clonotypes from a total of eight lineages (**Figure 5A** and **Supplemental Table S12**). For 3/8 lineages, the germline rAbs were non-reactive to most or all cell lines by intracellular flow cytometry, whereas their descendent clonotype rAbs showed increased reactivity (**Figure 5B)**. For the remaining 5/8 lineages, the germline rAbs showed moderate reactivity, and descendent rAbs showed either increased (2/5) or decreased (3/5) reactivity. Similar patterns were seen for five cell surface-reactive lineages. Thus, a substantial proportion (5/8) of lineages evolved toward increased reactivity against non-autologous cell lines, suggesting a breach of peripheral tolerance against broadly expressed, public antigens.

**Figure 5.**
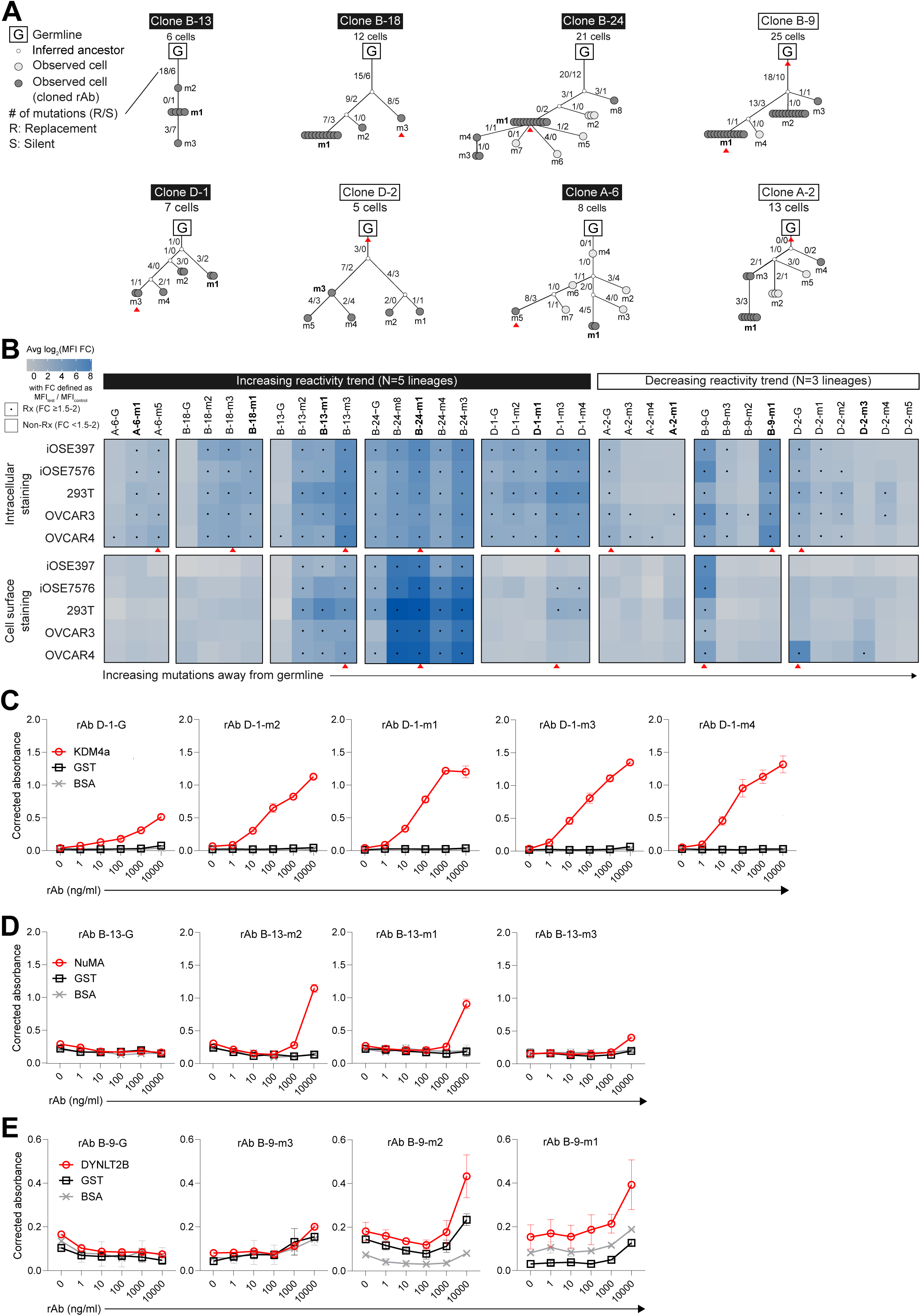
Evolution of TIL-B reactivity within clonal lineages. **(A)** Lineages of selected TIL-B clones. Mutant (m) number identifies individual clonotypes. Number of circles = number of cells. Numbers on branches indicate the total number of nucleotide mutations between relevant clonotypes. Bolded clonotypes are the dominant clonotype used in initial analyses. **(B)** Heatmap summarizing TIL-B rAb reactivity for all 35 clonally related rAbs as measured by intracellular and cell surface flow cytometry. Red arrows in panels A and B indicate the individual clonotype within each lineage that demonstrated the highest reactivity (measured by total median MFI) across all cell types evaluated. **(C-E)** ELISA data with rAbs from TIL-B clonal lineages D-1 **(C)**, B-13 **(D)**, and B-9 **(E)** and recombinant KDM4a, NuMA, DYNLT2B, GST, and BSA proteins. BSA: Bovine serum albumin. GST: Glutathione S transferase. MFI: Median fluorescence intensity.

We also assessed clonal evolution to the identified target antigens KDM4a, NuMA, and DYNLT2B. Within the KDM4a-reactive lineage (D-1), the germline rAb showed low KDM4a-specific reactivity, whereas all four descendant clonotype rAbs showed increased reactivity (**Figure 5C**). Within the NuMA-reactive lineage (B-13), the germline rAb was non-reactive to NuMA, whereas two of three descendant clonotype rAbs showed increased reactivity (**Figure 5D**). Within the DYNLT2B-reactive lineage (B-9), the germline rAb was non-reactive to DYNLT2B, while two of three descendant clonotypes rAbs showed increased, albeit low, reactivity (**Figure 5E**). Thus, TIL-B lineages generally evolved towards increased reactivity to these self-antigens.

### A large proportion of TIL-B-derived antibodies are polyreactive

The fact that many rAbs bound both intracellular and cell surface antigens, combined with the IP and WB results, suggested that some rAbs may be polyreactive. Therefore, we subjected all 51 rAbs to a polyreactivity assay that is widely used in the autoimmunity, transplantation and infectious disease fields.^54–58^ By this assay, 9/51 (17.7%) TIL-B rAbs scored as polyreactive (**Figure 6A** and **Supplemental Figure S11**), all of which had shown reactivity to cell lines and/or tissues by flow cytometry, WB or IHC. The rAbs that were used to identify KDM4a, NuMA, and DYNLT2B all scored negative. Polyreactive rAbs were identified in 4/4 patients for which a reasonable sample size was available, indicating that polyreactivity is prevalent in HGSC. Indeed, the prevalence may be as high as 25.5% if one includes rAbs that appeared polyreactive by WB (**Supplemental Figure S8A**) and/or IP-silver stain (**Supplemental Figure S9B**).

**Figure 6.**
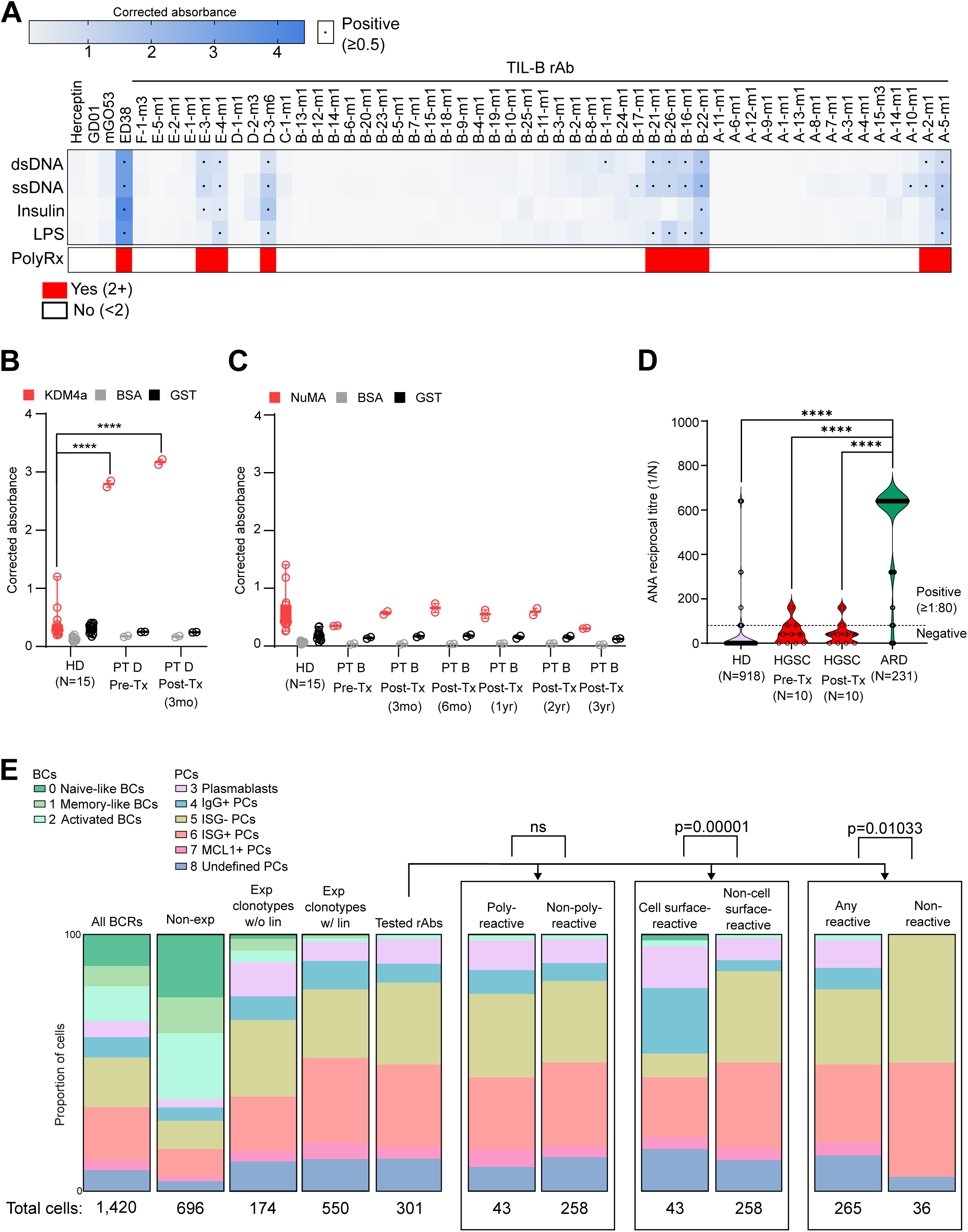
Polyreactivity, serology, and cellular phenotypes associated with TIL-B antibody responses. **(A)** Heatmap summarizing polyreactivity (polyrx) ELISA data for all 51 TIL-B rAbs. **(B)** ELISA with serum from patient (PT) D and 15 healthy donors (HD) against recombinant KDM4a, BSA, and GST. **** = p < 0.0001 as measured by two-way ANOVA. **(C)** ELISA with serum from PT B and 15 HD and recombinant NuMA, BSA, and GST. **(D)** Anti-nuclear antibody (ANA) titres from immunofluorescence staining with HGSC patient serum (N=10) in reference to previously published ANA results for HD and autoimmune rheumatic disease (ARD) patients^63^. **(E)** Bar plots showing the distribution of TIL-Bs across B cell (BC) and plasma cell (PC) phenotypic clusters within different B cell receptor (BCR) groups based on expansion status or related rAb reactivity. P values measured by Fisher’s exact test. BSA: Bovine serum albumin. dsDNA: Double-stranded DNA. Exp: Expanded. GST: Glutathione S transferase. Lin: Lineage. LPS: Lipopolysaccharide. ssDNA: single-stranded DNA. Tx: Treatment.

### Evaluating the link between TIL-B and systemic antibody responses

The observed self-reactivity of many TIL-B antibodies prompted us to investigate whether these responses were accompanied by systemic immunity. Compared to healthy donor controls, serum antibodies from patient D (the source of anti-KDM4a rAb D-1-m1) strongly recognized KDM4a at both pre- and post-treatment time points (**Figure 6B**), indicating a systemic autoantibody response to this self-antigen. In contrast, serum antibodies from patient B (the source of anti-NuMA rAb B-13-m1 and anti-DYNLT2B rAb B-9-m1) did not recognize NuMA (**Figure 6C**) or DYNLT2B (**Supplemental Figure S10A**) at any evaluated time point. Serum samples from these and eight other TIL-B-positive HGSC patients from cohort B were subjected to a clinical anti-nuclear antibody (ANA) test and, if negative for ANAs, to a rheumatoid factor (RF) test.^59–62^ Sixty percent (6/10) of HGSC patients (including patient D) scored negative for ANAs and RF (**Figure 6D** and **Supplemental Figure S10B**). The remaining 4/10 patients (including patient B) scored lowly positive for ANAs, with titres that were below the diagnostic threshold for autoimmune disease.^63^ Thus, self-reactive TIL-B responses can be accompanied by systemic autoantibodies in some patients while remaining below the threshold of clinical autoimmunity.

### Evaluating the link between TIL-B clonotype, tumor reactivity and phenotype

We next investigated how the observed rAb reactivity patterns of TIL-Bs mapped to the eight phenotypic clusters we had identified by scGEX (**Figure 3**). As previously mentioned, all but one of the 51 evaluated rAbs were derived from clonally expanded PCs, and 78.4% were members of lineages; therefore, these subgroups served as particularly relevant comparators.

Consistent with our prior analyses (**Figure 3**), expanded clonotypes without lineages were markedly skewed toward PC phenotypes, especially the ISG-PC phenotype (**Figure 6E** and **Supplemental Table S13**). Expanded clonotypes with lineages showed an even more pronounced skewing toward PC phenotypes, especially the ISG+ PC and ISG-PC phenotypes. As expected, the 51 clonotypes that were assessed as rAbs showed a similar phenotypic distribution as expanded clonotypes with lineages. Clonotypes that scored as polyreactive showed a similar phenotypic distribution as those that were non-polyreactive (**Figure 6E**).

Strikingly, clonotypes that exhibited cell surface reactivity showed a significant skewing toward the IgG+ PC and PB phenotypes, mostly at the expense of the ISG-PC phenotype. In contrast, clonotypes whose rAbs were found to be non-reactive by any of our assays almost exclusively exhibited the ISG+/- PC phenotypes, a significant difference compared to clonotypes that were reactive by any assay. We could not draw firm conclusions about the clonotypes that reacted to KDM4a, NuMA or DYNLT2B, owing to the small numbers of corresponding cells; however, their phenotypes were generally consistent with other reactive clonotypes (**Supplemental Table S13**).

### Spatial mapping of clonally expanded and tumor-reactive TIL-Bs

To assess the spatial distribution of TIL-Bs with known clonotypic, phenotypic, and tumor reactivity status, we used a modified spatial transcriptomics platform to generate linked spatial GEX (spGEX) and BCR (spBCR) sequencing data on tumors from 11 HGSC patients, including ten from cohort B (**Supplemental Table S7**). In general, BCR sequences were found in both tumor epithelium and stroma, sometimes forming “BCR hotspots” that by spGEX comprised lymphoid aggregates containing TIL-Bs, TIL-Ts and other immune cell types (**Figure 7A**). With the exception of patient D, tumors from the other ten patients showed no statistically significant differences in the spatial distribution of non-expanded clones versus expanded clones (with or without lineages) across the epithelial, boundary or stromal regions (**Supplemental Figure S13A**). For example, in patient B (who by scRNA-seq had the largest number of PCs and expanded clones; **Figure 3A,B,D**), non-expanded and expanded clones (with or without lineages) showed similarly broad distributions across the tumor bed (**Figure 7A**). Moreover, there were no examples of sizable expanded clones (using a threshold of 15 or more sequences) that were exclusively localized to tumor epithelium, stroma or boundary regions; the sole exception was patient N’s tumor, in which boundary regions were vastly over-represented (approximately 90% of total spots). Furthermore, in 9/11 patients, non-expanded and expanded clones (with or without lineages) showed similar distributions inside and outside of BCR hotspots (**Supplemental Figure S13B**). The two exceptions were patients B and D, in which expanded clones with lineages were significantly enriched in BCR hotspots (p = 0.00057 and 0.001652, respectively) (**Figure 7B**).

**Figure 7.**
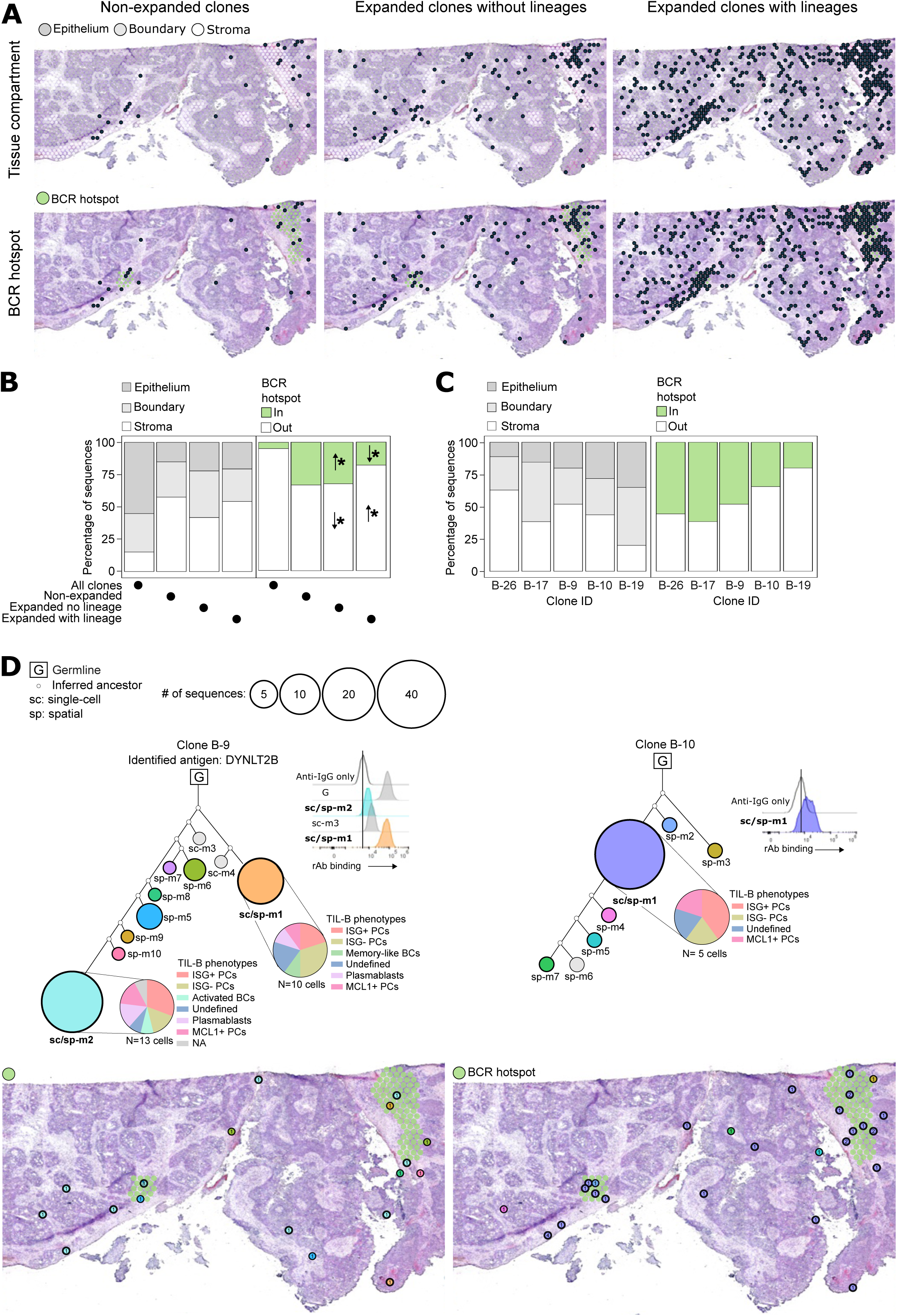
Spatial mapping of clonally expanded and tumor-reactive TIL-Bs. **(A)** Spatial mapping of BCRs by expansion status in relation to tissue compartments and BCR hotspots for patient B. **(B,C)** Bar plots showing the tissue and BCR hotspot distribution patterns of the five clones from patient B detected across both datasets (scRNA-seq and spBCR-seq) related to cloned rAbs by expanded status **(B)** and by clone **(C)**. Asterisks in (B) indicate significantly increased (up arrow) or decreased (down arrow) prevalence of the indicated clone type in that tissue compartment as measured by Chi-square test and adjusted Pearson residuals. **(D)** Integrated sc/sp lineages for clones B-9 and B-10 with related rAb reactivity to OVCAR3 cells as measured by flow cytometry and spatial mapping onto tumor tissue in relation to BCR hotspots. Clonotypes detected in both scRNA-seq and spRNA-seq datasets corresponding to evaluated rAbs are outlined with thicker lines on lineages. BCR hotspot: Gaussian BCR kernel density estimate above the 95^th^ percentile of the overall distribution in a given patient.

Uniquely for patient B, we had matched rAb reactivity and spatial data for five expanded clones with lineages. Irrespective of their rAb reactivity profiles, all five clones were broadly distributed throughout the epithelial, boundary and stromal regions as well as BCR hotspots (**Figure 7C**). For example, for lineages B-9, B-10 (**Figure 7D**) and B-26 (**Supplemental Figure S13C**), BCR clonotypes were found in epithelium, stroma, and boundary regions, as well as inside and outside of BCR hotspots, despite the two former lineages corresponding to non-polyreactive, intracellular-reactive rAbs and the latter to a polyreactive, intracellular- and cell surface-reactive rAb (**Figure 7D** and **Supplemental Figure S13B**). Thus, self-reactive PCs were able to penetrate the entire tumor bed, including the malignant epithelium, reflecting the prognostically favorable pattern characteristic of BME4 tumors.

## DISCUSSION

By evaluating the TME from the perspective of TIL-B responses, we identified four infiltration patterns. While BME2 and BME3 had considerable numbers of stromal TIL-Bs, BME4 showed a marked increase in BCs and PCs in both stroma and epithelium, a pattern that was uniquely associated with prognostic benefit. BME4 also harbored dense TIL-Ts and PD-L1+ TAMs, consistent with our findings in long-term survivors of HGSC.^27^ Most BME4 tumors were PC-rich, suggesting that PC responses drive the prognostic benefit associated with TIL-Bs.

Indeed, in sharp contrast to BCs, PCs showed strong evidence of active antigen recognition, including clonal expansion, SHM, class switching (mostly to IgG1), and intricate lineage structures. Clonally expanded PCs expressed antibodies against broadly expressed intracellular antigens (including NuMA, KDM4a and DYNLT2B) and, less frequently, cell surface antigens. About one-fifth of antibodies were also polyreactive. PCs with verified tumor reactivity and/or polyreactivity spanned all five molecular phenotypes of PCs, and cell surface-reactive PCs showed a marked skewing toward PB and IgG+ PC phenotypes. Tumor-reactive PC clones infiltrated the epithelial, boundary, and stromal regions of tumors, recapitulating the BME4 pattern. Collectively, our findings demonstrate that infiltration of tumor epithelium by highly activated, clonally expanded, self- and poly-reactive PCs is a defining feature of the most prognostically favorable anti-tumor immune responses.

As per prior studies^40,64,65^, TIL-B-positive tumors fell into BC- vs PC-rich subgroups. It remains unclear why some tumors are BC-rich and how this influences anti-tumor immunity.

Although BC-rich tumors had abundant T cells, other factors required for BC-to-PC differentiation could potentially be impaired in this subgroup. For example, HGSC has fewer and smaller TLS compared to lung cancer^66^, which may result in decreased intra-tumoral PC generation^67,68^. PCs and memory BCs can also arise through extrafollicular (EF) responses which, in theory, could also be impaired in BC-rich tumors. Finally, in pancreatic cancer, T cell-derived IL-35 was shown to prevent PC differentiation^69^; however, we found negligible co-expression of the *IL12A* and *EBI3* genes (which encode IL-35) in T cells in our dataset.

The close proximity of BCs to PCs, TIL-Ts and TAMs suggests they may play important mechanistic roles. Our scRNA-seq experiments revealed three main BC phenotypes (naïve-like, memory-like, and activated), all of which have been reported in human cancer.^34,38–41,43^ All three BC clusters, especially memory-like BCs, scored high for antigen presentation gene modules (**Supplemental Figure S4A**), consistent with prior reports of BCs serving as APCs in autoimmune disease, transplantation and cancer.^70,71^ As APCs, BCs are thought to have an advantage over dendritic cells, as they can potentially use their surface BCRs to enhance uptake of cognate antigen, thereby amplifying T cell responses to less abundant antigens. However, this concept seems less likely to operate in HGSC, as the vast majority of BCs were not clonally expanded. Instead, if appropriately stimulated, BCs could potentially serve as generic APCs to T cells, as demonstrated *in vitro*^72^.

Another well-documented function of BCs is cytokine production.^73^ Indeed, we found that BCs (in particular memory-like BCs) expressed *TNF, LTA*, and *LTB* (**Supplemental Figure S4B**), which could provide costimulatory signals to T cells and macrophages and promote the formation of TLSs^74,75^. We did not detect an “atypical memory” (Atm; also known as double-negative 2, DN2) BC cluster reported by others in cancer^38^, autoimmunity^76^ and healthy donors^77^; this may be due to under-sampling, as this phenotype represents only 3-4% of TIL-Bs^38^. Although *TGFB1* was expressed at low levels across most TIL-B clusters, we saw no evidence for a distinct cluster of Bregs^78,79^, and only negligible levels of *IL10, IL12A*, and *EBI3* were detected in any TIL-B cluster (**Supplemental Figure S4B**), which is consistent with other scRNA-seq datasets from human cancer^34,38–41,43^.

We cloned and characterized antibodies from all clonally expanded TIL-Bs in our dataset and found that the majority (88%) bound intracellular antigens, consistent with a smaller study in pancreatic cancer^80^. Antibodies against intracellular proteins could be generated by the release of cytoplasmic and nuclear antigens by dying tumor cells under immunogenic conditions, such as the coincident release of DNA and RNA.^81^ Additionally, apoptotic cells can express unusual protein isoforms and/or cleavage products, as reported for actin.^82^ Indeed, granzyme B-mediated cleavage of NuMA creates novel termini that could potentially induce autoantibody responses in patients with autoimmune disease ^83^. Thus, some self-reactive TIL-B responses could result from T and NK cell-mediated cytotoxicity against tumors.

Intriguingly, we observed significantly different phenotypic profiles between intracellular- and cell surface-reactive TIL-Bs, suggesting they experience distinct forms of antigen encounter *in vivo* and/or mediate different effector functions. While cell surface-reactive TIL-B antibodies could promote anti-tumor immunity through mechanisms such as CDC, ADCC and ADCP^31,33^, it is less clear how antibodies to intracellular antigens might enter tumor cells and mediate effector functions. Possible entry mechanisms include (a) endocytosis through the adaptor protein 2 complex^34^; (b) “hitch-hiking” on receptors for the bound antigen^84^; (c) and for IgA, transcytosis via the poly Ig receptor^33^. Once inside cells, antibodies could potentially interfere directly with the function of their cognate antigen or trigger antigen degradation through ADIN.^34,85^ Intriguingly, KDM4a promotes tumor growth in many cancers^86^, opening the possibility of direct anti-tumor effects by TIL-B-derived antibody D-1-m1. Antibodies can also recognize antigens that are normally intracellular but become aberrantly exposed on the surface of tumor cells *in vivo*, such as nucleolin in neuroblastoma^87^ and actin in breast cancer^82^; indeed, many of our rAbs showed both intracellular and cell surface reactivity. Antibodies can also form immune complexes with intracellular antigens released from dying cells; in turn, immune complexes can enhance tumor antigen presentation to CD4 and CD8 T cells^88^ and, in pathological settings such as lupus nephritis, are thought to promote end-organ damage through multiple mechanisms^89^. Finally, some TIL-B antibodies could operate by modulating other cells in the TME, such as immune cells, endothelial cells, and stromal cells, as suggested by our IHC and flow cytometric data.

By studying clonally related rAbs from lineages, we found that TIL-Bs frequently evolved towards increased reactivity to tumor cells (OVCAR3) and self-antigens, consistent with a prior report.^31^ Moreover, a substantial proportion of TIL-B antibodies (18% to 25.5% by our various assays) were polyreactive, a property that was predicted in a prior study of BCR sequence features in cancer^90^. In healthy donors, polyreactivity is exhibited by 4-6% of mature naïve BCs from peripheral blood^54,91^ and 9% of IgG+ PCs from bone marrow^58^. Self- and poly-reactivity are features associated with not only autoimmune disease but also anti-viral immunity; indeed, many broadly reactive anti-viral antibodies are also self-reactive and can exacerbate autoimmune disease through mechanisms such as CDC, ADCC, ADCP and induction of general inflammation.^92^ In theory, poly-reactivity could provide a means for TIL-B to prevent tumors from escaping immune control via antigen loss, as has been hypothesized in infectious disease.^93^

An important question is whether TIL-B responses have a systemic component which could potentially enable control of metastatic disease. In support of this idea, we detected linked TIL-B and serological responses to KDM4a in patient D. In urothelial cancer, serum antibodies against KDM4a have been associated with immune-related adverse events after treatment with ICIs^94^, suggesting that peripheral tolerance to this antigen might be relatively easily breached.

We failed to detect systemic antibody responses to DYNLT2B or NuMA; however, serum antibodies to NuMA are well described in patients with systemic erythematosus and Sjögren’s syndrome^95^, indicating that this antigen too is prone to recognition by self-reactive BCs. Finally, we detected low-titre ANAs in 4/10 patients, which is consistent with prior reports in ovarian^96^ and other cancers^97^. Thus, although more work is required, TIL-B responses may well have a systemic component. This could provide an important means to control metastatic tumor sites; however, it also brings potential risks that need to be considered when designing TIL-B-directed immunotherapies.^28^

PCs could also mediate tumor control through antibody-independent functions. Indeed, an important finding from this study is that individual PC clones exhibit wide but sparse distribution patterns, raising the question of whether they could produce sufficient levels of antibody to achieve a significant anti-tumor effect. In the fields of autoimmune disease and transplant rejection, controversy remains regarding the relative importance of antibody-versus cell-based-mechanisms, as B cell depleting therapies can reduce pathological symptoms without necessarily dampening systemic antibody levels.^98,99^ In autoimmune disease and transplant rejection, PCs can secrete cytokines and present antigen through antibody-independent mechanisms^100,101^, as discussed above for BCs. Moreover, PCs secreting IL-17^102^, IL-35 and IL-10^103^ play important roles in anti-bacterial and anti-viral immune responses. While we did not detect appreciable expression of these cytokines by PCs in our scRNA-seq data (**Supplemental Figure S4B**), these and other antibody-independent mechanisms warrant further study.

This is the first study to decipher TIL-B phenotypes, tumor reactivity, and spatial distributions at clonal resolution in human cancer. Nonetheless, a sample size of 1,741 TIL-Bs may have limited our ability to detect phenotypic clusters seen in other larger scRNA-seq studies.^34,38–41,43^ Moreover, with this number of TIL-Bs, we could not fully assess the extent of BCR sharing between BCs and PCs and clonal expansion among BCs. A further limitation is that our antigen discovery methods did not allow identification of patient-specific neoantigens; however, 90% of the rAbs we tested (representing 100% of significantly expanded TIL-B clones from six patients) showed reactivity to at least one non-autologous cell line, indicating that the vast majority of clonally expanded TIL-Bs recognize public antigens.

Our findings have implications for the design of immunotherapies that leverage TIL-B responses. A key question is whether the prognostic benefits of TIL-B are primarily due to antibody-versus cell-based effector mechanisms. Some authors have emphasized the former, providing evidence that TIL-B-derived antibodies against targets such as brain-derived neurotrophic factor and MMP14 can have anti-tumor effects.^31,33^ However, such mechanisms would require that each cancer patient develop their own neutralizing antibody response against one or more targets that are essential to their tumor’s viability and/or progression. Moreover, such antibodies would need to reach sufficiently high systemic levels to mediate anti-tumor effects across metastatic sites. While this may be conceivable in some cases, it seems an unlikely explanation for the strong prognostic effect of TIL-Bs seen across diverse patient populations and cancer types.^29^ That said, it may be possible to identify highly effective anti-tumor antibodies from cancer patients with exceptional TIL-B responses and develop recombinant versions of these as therapeutic agents. Encouraging in this regard is our finding that 20% of TIL-B rAbs recognized cell surface antigens, many of which exhibited some degree of tumor and/or epithelial specificity. As an alternative concept, the anti-tumor effects of TIL-B antibodies could be attributable to their poly-reactive properties. Given that polyreactive antibodies are prevalent in all healthy individuals^54–56,58,91,104^, it is conceivable that these could develop into TIL-B responses in a substantial proportion of patients, facilitated by the inflammatory milieu typical of most cancers^105^. Polyreactive antibodies could provide a means for TIL-Bs to combat the extensive intra-tumoral heterogeneity and evolutionary dynamics of HGSC and other advanced cancers. However, such a mechanism presumably requires localized production of such antibodies by TIL-Bs to prevent progression to systemic autoimmunity. This could be challenging to emulate using recombinant antibodies and may instead require novel, tumor-directed delivery platforms. As for the cellular functions of TIL-Bs, these could potentially be induced by vaccination strategies that induce coordinated B cell and T cell responses. Current approaches with neoantigen vaccines might be helpful in this regard, as one could potentially induce antigen spreading beyond the targeted neoantigens^29^ Other strategies being pursued involve novel immunomodulatory agents (e.g., stimulatory antibodies against CD40L, ICOS) and oncolytic viruses. However, based on our findings and others^28^, these latter “antigen agnostic” approaches should be approached with caution, given the high rate of self-reactivity exhibited by TIL-Bs. Thus, TIL-Bs hold great promise for immunotherapy provided we can learn to safely manipulate the relaxation of peripheral tolerance that underlies their native mechanisms.

## RESOURCE AVAILABILITY

### Lead contact

Requests for further information and resources should be directed to and will be fulfilled by the lead contact, Brad H. Nelson.

### Materials availability

Plasmids encoding TIL-B-derived recombinant antibody sequences are available upon request.

### Data and code availability

All data needed to evaluate the conclusions in the paper are present in the paper and/or the supplemental materials. The accession numbers can be found in the key resources table. This paper does not report any original code. For any additional information needed to reanalyze the data reported in this paper, please contact the lead contact.

## ACKNOWLEDGEMENTS

We thank the patients and their families for generously donating their time, tissues and data to make this research possible. We thank Dr. Michel Nussenzweig for kindly sharing the ED38 and mGO53 antibodies. A.C.B. was supported by a Doctoral Award from CIHR (Frederick Banting and Charles Best Canada Graduate Scholarship, FBD — 177882). C.M.L. was supported by post-doctoral fellowships from Canadian Institutes of Health Research (CIHR) (Banting postdoctoral fellowships programme, 429161) and Michael Smith Health Research BC’s Research Trainee Program (RT-2020-0630). B.H.N. is supported by the British Columbia Cancer Foundation. This work was also supported by grants from CIHR [application numbers 427647 and 487192 and the Department of Defense OC230183. Some graphics were made using biorender.com.

## AUTHOR CONTRIBUTIONS

Conceptualization: ACB, BHN, CML, GBM, CS, FN Data curation: ACB, CML, KS, GLN, SK

Formal analysis: ACB, CML, KS, GLN, SK Funding acquisition: ACB, BHN, CML

Investigation: ACB, CML, JS, SK, KS, JD, BG, LM, JW, VNP, BM, KW, ML, TO, SES, GLN

Methodology: ACB, CML, JS, SK, KS, JD, BG, LM, JW, KM, VNP, NSG, EAC, PTH, AM, GC, SES, GLN, GBM, CS, BHN

Project administration: BHN Supervision: BHN, ACB, KM Visualization: KS, ACB, CML

Writing – original draft: ACB, BHN, CML Writing – review and editing: ACB, BHN, CML

## SUPPLEMENTAL INFORMATION

**Supplemental Figure S1. Immune cell densities and survival analyses in cohort A. (A)** Epithelial and stromal immune cell densities by B cell-defined microenvironment (BME) type. The vertical dashed lines indicate the median. Y-axes represent probability densities. **(B,C)** Kaplan-Meier survival plots for patients separated by BME type for all stages (**B**) and epi-high and epi-low status (**C**). Epi: Epithelium. TAMs: Tumor-associated macrophages.

**Supplemental Figure S2. Marker genes used to define immune cell clusters identified by scRNA-seq in cohort B.**

**Supplemental Figure S3. Marker genes used to define TIL-B clusters identified by scRNA-seq in cohort B. (A)** UMAP clustering restricted to TIL-Bs using single-cell GEX (scGEX) data from HGSC samples. Each dot represents a cell. Colors represent TIL-B clusters identified by graphical clustering and marker genes. **(B,C)** Dot plots showing the marker genes used to annotate the three BC and six PC clusters presented in panel A. **(D)** Heatmap showing the top 10 genes expressed in each of the five PC clusters (p-value < 0.05, adjusted p-value < 0.05 and average log2(fold-change) > log2(1.5)). **(E)** Violin plots showing the distribution of gene module scores for 3 REACTOME pathways, associated with antibody secretion^47^ across TIL-B clusters. Each dot represents a cell. ISG+/-: Interferon-stimulated genes positive/negative.

**Supplemental Figure S4. TIL-B antigen presentation module scores and cytokine expression from scGEX data. (A)** Antigen presentation REACTOME gene module scores by TIL-B cluster. **(B)** Cytokine expression levels by TIL-B cluster.

**Supplemental Figure S5. Mutation counts for TIL-Bs from scBCR data. (A,B,C)** Mutation counts (nucleotide level, including synonymous and non-synonymous) for B cells (BCs) and plasma cells (PCs) individually by patient ranked by BC:PC ratio **(A),** for each of the nine TIL-B clusters **(B)**, and by expansion status (non-expanded [i.e., singletons], expanded no lineage, and expanded with a lineage).

**Supplemental Figure S6. Additional flow cytometry, western blot (WB), and immunohistochemistry (IHC) data relating to TIL-B rAb binding. (A)** Representative flow cytometry histograms of intracellular and cell surface staining for various TIL-B rAbs to demonstrate thresholds for positive binding (fold change [FC] of 1.5 or more for intracellular staining and 2.5 or more for cell surface staining). **(B)** Percentage of reactive TIL-B rAbs identified by each method. **(C)** Flow cytometry histograms showing the cell surface binding of TIL-B rAbs (N=7, blue) that recognized the surface of OVCAR3 cells and control antibodies (N=4, green) with cells collected with either 0.5 mM EDTA-PBS or Trypsin. Vertical lines indicate the baseline signal from anti-IgG only control sample. Intra: Intracellular.

**Supplemental Figure S7. Flow cytometry staining with TIL-B rAbs using healthy donor peripheral blood mononuclear cells (PBMC). (A)** Gating strategy to identify indicated cell populations and the rAb binding cell population. **(B)** Flow plots showing binding of 14 TIL-B rAbs from initial screening (corresponding to Figure 3A). **(C)** Histograms showing binding of two non-binding TIL-B rAbs (grey) and six binding TIL-B rAbs (blue) determined to be true binders of PBMCs based on follow up robust staining (see Methods). Histograms shown for all cell populations depicted in gating strategy in **(A)**. FC: Fold change. Macs: Macrophages. NK: Natural killer.

**Supplemental Figure S8. All western blot (WB) data and additional immunohistochemistry (IHC) data relating to TIL-B rAb binding.** Red outline indicates rAbs that were determined to be reactive by WB as visualized by detectable banding patterns. IB: immunoblot. **(B)** Representative IHC images for 3 weakly IHC-reactive TIL-B rAbs against high-grade serous ovarian cancer (HGSC) and normal ovary tissue.

**Supplemental Figure S9. Immunoprecipitation(IP)-Western blot (WB) and IP-silver stain (SS) data for TIL-B clones D-1 and B-13. (A)** IP-WB with IP outputs using TIL-B rAb D-1-Germline (G) and clonal descendant D-1-m4 or B-13-G and B-13-m1. Blots were probed with commercial anti-KDM4a or anti-NuMA antibodies. **(B)** IP-SS with IP outputs using the same TIL-B rAbs as in A.

**Supplemental Figure S10. RNA expression of TIL-B target antigens across normal and tumor tissues.** Gene expression of the three identified TIL-B target antigens in paired normal and tumor tissues was obtained from re-processed GTEx and TCGA data available from the UCSC Xena project^106^ and analyzed with DESeq2^107^. On the heatmaps, RNA expression is shown as average variance stabilized-transformed log2 counts. For each TIL-B target, asterisks indicate significant over-expression in a given tumor tissue relative to its normal counterpart (shrunken log2(Tumor/Normal) ≥ log2(1.5) and Wald P and Benjamini-Hochberg adjusted P ≤ 0.05). Tumor types are listed by TCGA abbreviations which are described in **Supplemental Table S11** along with sample size for both groups (tumor and normal).

**Supplemental Figure S11. Dose response curves for polyreactivity ELISAs.** Reactivity of all 51 main clone TIL-B rAbs to the four antigens used to assess polyreactivity by ELISA. Positive control: ED38 (black dashed line). Negative controls: mGO53 (green), Herceptin (purple) and GD01 (grey). dsDNA: double-stranded DNA. LPS: Lipopolysaccharide. ssDNA: single-stranded DNA.

**Supplemental Figure S12. Serum antibody testing high-grade serous ovarian cancer (HGSC) patient samples from cohort B. (A)** Reactivity of serum from patient B to identified TIL-B antigen DYNLT2B, GST, and BSA. **(B)** Anti-nuclear antibody (ANA) titres of serum from HGSC patients in cohort B. BSA: Bovine serum albumin. DYNLT2B: Dynein light chain type 2B. GST: Glutathione S-transferase. Mo: month. Tx: treatment. Yr: year.

**Supplemental Figure S13. Spatial transcriptomics reveals distribution patterns of TIL-Bs relative to expansion status and antibody reactivity. (A,B)** Bar plots showing the proportion of B cell receptor (BCR) sequences relating to tissue compartment **(A)** and identified BCR hotspots **(B)** per patient. BCR hotspot: a spot with a Gaussian BCR kernel density estimate above the 95^th^ percentile of the overall distribution in that patient. **(C)** Integrated single-cell (sc) and spatial (sp) lineages for clone B-26 with related rAb reactivity to OVCAR3 cells as measured by flow cytometry and spatial mapping onto tumor tissue in relation to BCR hotspots. Exp: expanded. Lin: Lineage. W: with.

**Document S1:** Excel document containing Tables S1 -S13.

## METHODS

### Study subjects

Cohort A patient characteristics are detailed in **Supplemental Table S1**.

Cohort B: Patient characteristics are detailed in **Supplemental Table S7**. From our in-house tumor tissue repository (TTR), maintained by BC Cancer Biobanking & Biospecimen (BBRS) Research Services, we selected 14 primary (untreated) HGSC cases which exhibited a range of immune cell densities by IHC.

### Multicolor IF (mcIF)

Samples from cohort A were used to generate tissue microarrays (TMAs) composed of duplicate 0.6 mm cores per case using a manual tissue microarrayer (Beecher Instruments). To evaluate TILs, we developed two seven-color immunofluorescence panels to detect CD3, CD8, CD20, CD79a, FoxP3, PD-1, PD-L1, and pan-cytokeratin (panCK, see table below).

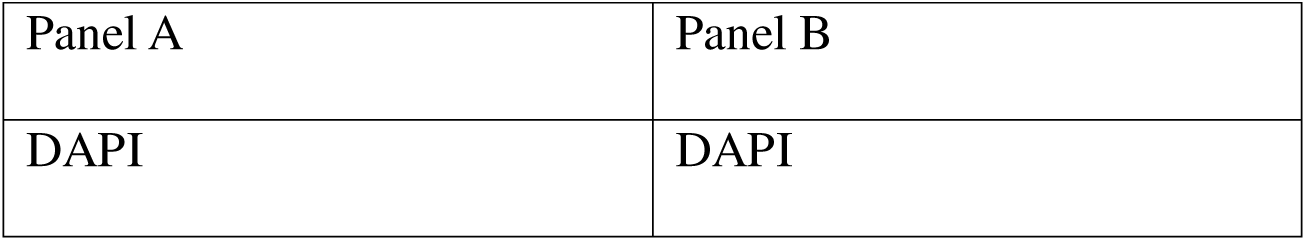
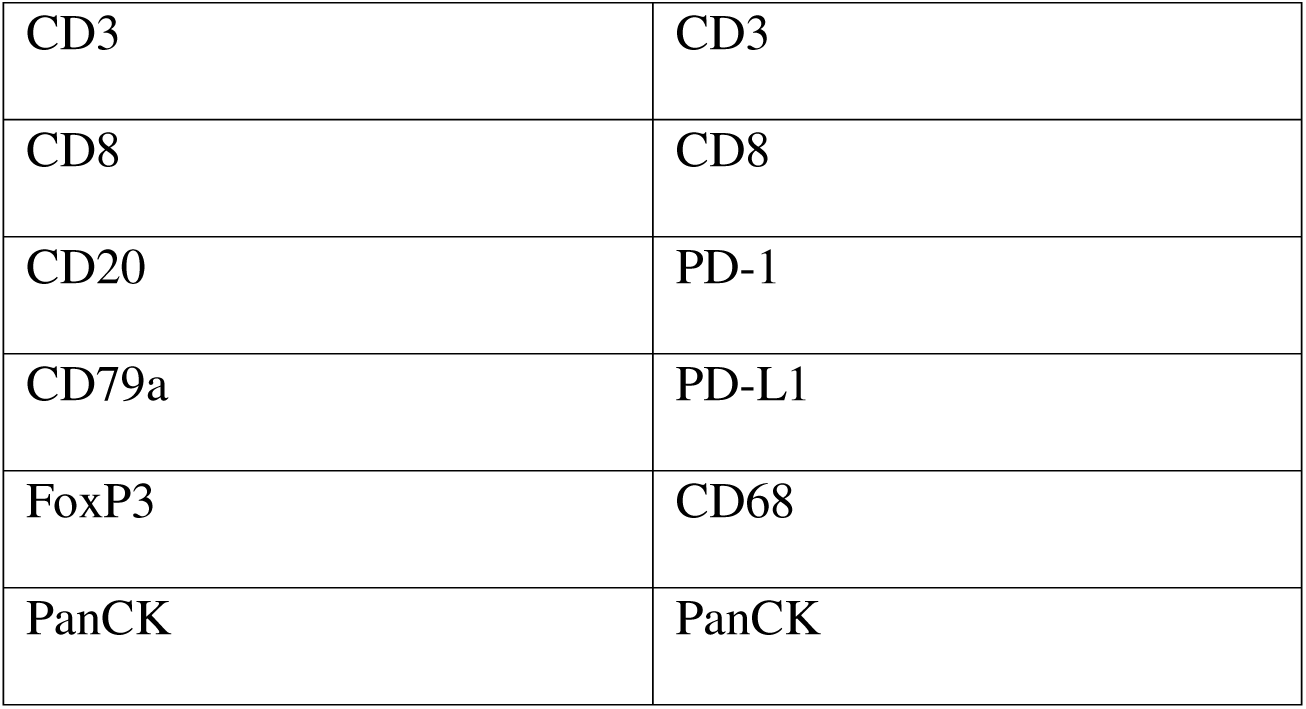

All steps were performed at RT unless otherwise noted. See tables below for antibody catalogue information, antibody dilutions, and further details.

Panel A:

| Round | Primary Antibody | Catalogue# | Clone | AR | Block | Dilution | Polymer | Fluorophore | Fluor Dilution |
| --- | --- | --- | --- | --- | --- | --- | --- | --- | --- |
| 1 | CD79a | Ab16698 | SP18 | DIVA | Terminator 12<br>no rinse | 1/500<br>DVG | M2Rb-HRP | OPAL520 | 1/500 |
| 2 | CD20 | CM004 | L26 | AR6 | Sniper 10 | 1/300<br>DVG | M2Ms-HRP | OPAL620 | 1/200 |
| 3 | CD8 | 108M-94 | C8/144B | AR6 | Sniper 5 | 1/600<br>DVG | M2Ms-HRP | OPAL690 | 1/100 |
| 4 | CD3 | CM110 | PS1 | AR6 | Sniper 5 | 1/100<br>RR | M2Ms-HRP | OPAL480 | 1/100 |
| 5 | FoxP3 | 14-4777-82 | 236A/E7 | AR6 | Terminator 12<br>no rinse | 1/100<br>DVG | M2Ms-HRP | OPAL570 | 1/200 |
| 6 | PanCK<br>+ | CM162 | AE1/AE<br>3<br>+5D3 | AR6 | Sniper 5 | 1/100<br>DVG | M2Ms-HRP | TSA-DIG | 1/100 |
| 7 | OPAL7<br>80 | FP15010<br>01KT | N/A | AR9 |  | 1/25 RR | N/A | N/A | N/A |

AR: Antigen retrieval. DVG: DaVinci green. RR: Renoir Red. Ms: Mouse. Rb: Rabbit. HRP: Horse radish peroxidase.

Panel B:

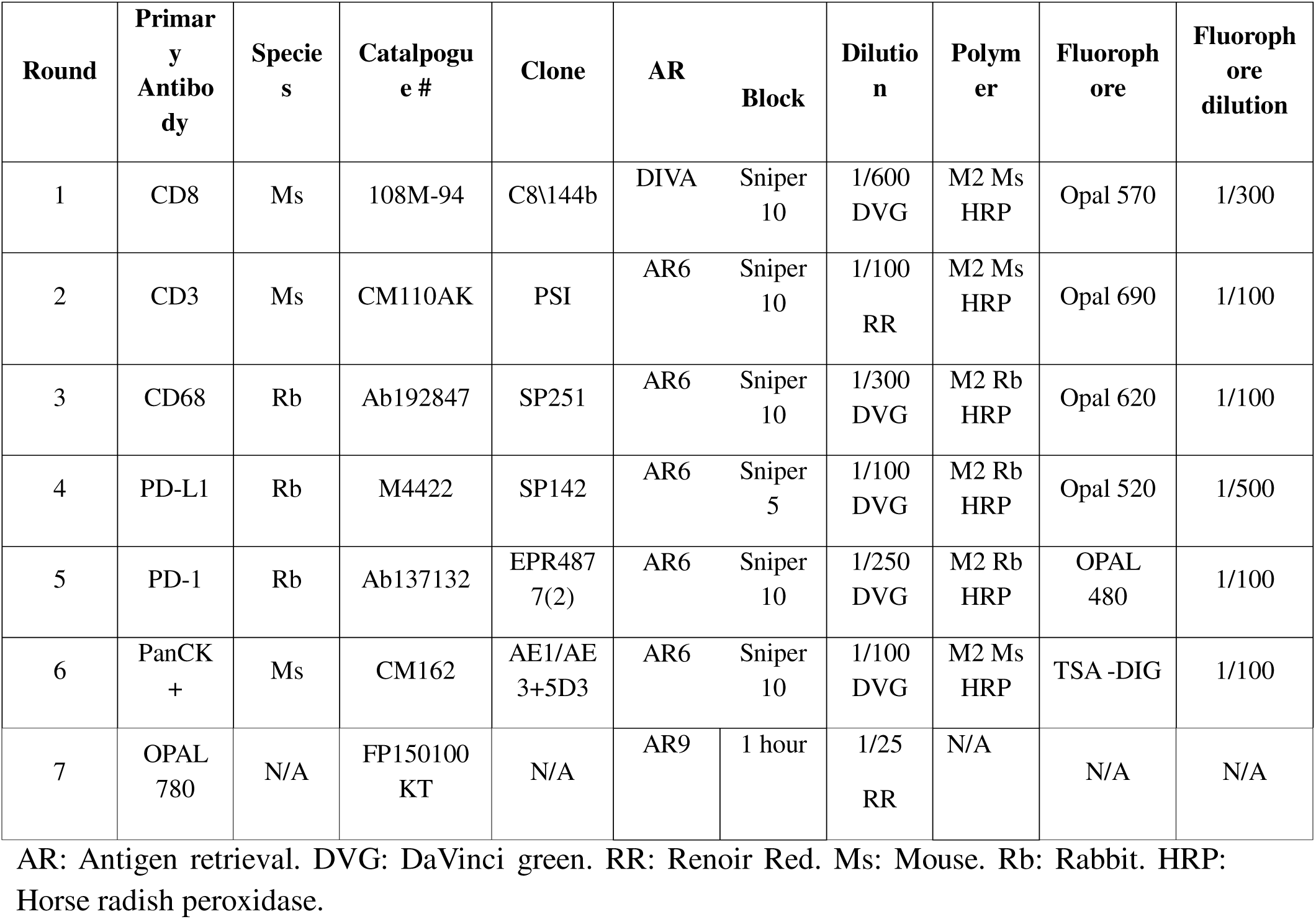

TMA slides containing formalin-fixed, paraffin-embedded (FFPE) tissues were baked at 37°C overnight, deparaffinized manually, and then subjected to an additional fixation with 10% neutral-buffered formalin (NBF) for 20 minutes. Slides were then subjected to antigen retrieval in a decloaking chamber with Diva Decloaker reagent for 15 minutes at 110°C followed by 10 minutes of cooling. Slides were then washed with dH_2_O and loaded onto the Intellipath FLX Autostainer to perform endogenous and non-specific blocking using peroxidased-1 (5 minutes) and background sniper (10 minutes), respectively.

For panel A, slides were then incubated with a primary antibody against CD79a with DaVinci Green diluent for 30 minutes followed by MACH 2 rabbit horseradish peroxidase (HRP) polymer for 10 minutes, then Opal 520 (Akoya Biosciences) for 10 minutes. To strip remaining antibody, slides were manually rinsed with dH_2_O and then antigen retrieval was performed with AR6 (Akoya) in a microwave followed by 15 minutes of cooling. For each subsequent staining round, slides were subjected to the same peroxidased-1 and background sniper treatment and then treated with the same antigen retrieval method after staining. For round 2, slides were stained with a primary antibody against CD20 with DaVinci Green diluent for 30 minutes, followed by Opal 650 (Akoya Biosciences) for 10 minutes. Slides were then re-loaded onto the Intellipath and stained with MACH 2 mouse HRP polymer for 10 minutes, followed by Opal 650 (Akoya Biosciences) for 10 minutes. For round 3, slides were stained with a primary antibody against CD8 in with DaVinci Green diluent for 30 minutes, then MACH 2 mouse HRP for 10 minutes, followed by Opal 690 (Akoya Biosciences) for 10 minutes. Between rounds 3 and 4, slides were cooled overnight. For round 4, slides were stained with a primary antibody against CD3 in with Renoir Red diluent for 30 minutes, then MACH 2 mouse HRP for 10 minutes, followed by Opal 540 (Akoya Biosciences) for 10 minutes. For round 5, slides were stained with a primary antibody against FoxP3 in with DaVinci Freen diluent for 30 minutes then MACH 2 mouse HRP for 10 minutes, followed by Opal 570 (Akoya Biosciences) for 10 minutes. For round 6, slides were stained with a primary antibody against PanCK in with DaVinci Green diluent for 30 minutes, then MACH 2 mouse HRP for 10 minutes, followed by Opal 480 (Akoya Biosciences) for 10 minutes. For round 7, slides were treated with TSA DIG probe for 10 minutes then stained with Opal 780 anti-DIG (Akoya Biosciences) for 1 hour. After round 7, slides were stained with DAPI for 5 minutes. Following staining, the slides were left to cure overnight then imaged the following day on the Vectra Polaris multispectral imaging system (Akoya Biosciences) using motif settings.

For panel B, the same staining method was used except different antibodies and polymers were used as per the table above. In detail, slides were then manually stained with a primary antibody against CD8 with DaVinci Green diluent for 30 minutes, then MACH 2 mouse HRP for 10 minutes, followed by Opal 540 (Akoya Biosciences) for 10 minutes. To strip remaining antibody, slides were manually rinsed with dH_2_O and then antigen retrieval was performed with AR6 (Akoya) in a microwave followed by 15 minutes of cooling. For each subsequent staining round, slides were first subjected to the same peroxidased-1 and background sniper treatment and then treated with the same antigen retrieval method after staining. For round 2, slides were stained with a primary antibody against CD3 with Renoir Red diluent for 30 minutes, followed by MACH2 Mouse HRP Polymer for 30 minutes then Opal 650 (Akoya Biosciences) for 10 minutes. For round 3, slides were stained with a primary antibody against CD68 in with DaVinci Green diluent for 30 minutes, then MACH 2 rabbit HRP for 10 minutes, followed by Opal 520 (Akoya Biosciences) for 10 minutes. Between rounds 3 and 4, slides were cooled overnight. For round 4, slides were stained with a primary antibody against PD-L1 in with DaVinci Green diluent for 30 minutes, then MACH 2 rabbit HRP for 10 minutes, followed by Opal 570 (Akoya Biosciences) for 10 minutes. For round 5, the background sniper treatment was extended for 10 minutes, and then slides were stained with a primary antibody against PD-1 in with Renaissance Background Reducing diluent for 30 minutes, then MACH 2 mouse HRP for 10 minutes, followed by Opal 690 (Akoya Biosciences) for 10 minutes. For round 6, slides were stained with a primary antibody against PanCK in with DaVinci Green diluent for 30 minutes, then MACH 4 universal probe HRP for 5 minutes, followed by Opal 480 (Akoya Biosciences) for 10 minutes. For round 7, slides were treated with TSA DIG probe for 10 minutes then stained with Opal 780 anti-DIG (Akoya Biosciences) for 1 hour. After round 7, slides were stained with DAPI for 5 minutes. Following staining, the slides were left to cure overnight then imaged the following day on the Vectra Polaris multispectral imaging system (Akoya Biosciences) using motif settings.

Cell types were defined by the following marker expression:

| Cell type | Marker |
| --- | --- |
| Regulatory T cell (Treg) | CD3+CD8-FoxP3+ |
| Presumptive CD4 T cell | CD3+CD8-PD1± |
| CD8 T cell | CD3+CD8+PD1± |
| Macrophage (Mac) | CD68+PDL1± |
| B cell | CD20+CD79a+ |
| Plasma cell | CD20-CD79a+ |

Qupath v3.0.2 was used to analyze mcIF images. A random trees classifier was trained on a randomly selected subset of images, using PanCK+ staining intensity to segment tissue into epithelium (PanCK+) and stroma (PanCK-). Each image was manually reviewed to ensure appropriate tissue segmentation, and images with <5% epithelium or staining artifacts were removed from downstream analysis (250 cores removed in cohort A, 77 cores removed in cohort C). The Stardist segmentation algorithm^108^ was used to segment cells. A cell classification classifier was trained based on cell features to categorize cells into cell types of interest. Cell classification was reviewed on a randomly selected set of images, and the classifier was fine-tuned in successive iterations until performance was deemed adequate. Cell densities for each cell type and total tumor core areas were summed across all available cores per patient. Cell densities were calculated as the number of cells per mm^2^ for cohort A and cells/high-powered field (HPF) for cohort B. Densities were then log-transformed (log10+1).

### B-cell-defined microenvironment (BME) clustering with cohort A

Clustering was done using the ConsensusClusterPlus package. Only TIL-B densities were used in the clustering (plasma and B cell densities in epithelium and stroma). We used an agglomerative clustering algorithm using ward.D2 linkage and euclidean distances. We set the repetitions parameter to 1000 and used all features (pFeature=1). Outputs from the ConsensusClusterPlus() function were inspected to choose the optimal number of clusters, based on the consensus matrix at each value of k and the relative change under the CDF curve plot.

Four clusters were chosen given the greatest difference in area under the CDF curve plot between k=4 and k=3.

### Cox proportional hazards model analysis with cohort A

To investigate the effect of covariates on survival, the Cox proportional hazards model was employed using the coxph function. We included age at diagnosis, stage and grade as covariates.

### Survival analysis with cohort A

To estimate differences in overall survival between the 4 BMEs, survival analysis was conducted using the “survival” package in R. Patients who did not die by follow-up were censored at their last follow-up date. We created an event indicator, where 1 indicated an event and 0 indicated censoring. Time to last follow-up/death was measured in months. We visualized the survival function using the Kaplan-Meier method as implemented in the survfit function.

Survival curve graphs were truncated at timepoints beyond which less than 10% of the original sample size was available (only for visualization). P-values are based on the log-rank test.

### Single-cell RNA sequencing

For each patient, frozen bulk tumor samples were thawed and subjected to dead cell removal (Miltenyi, 130-090-101) prior to staining with the following antibody cocktail: CD45-FITC (pan-lymphocyte marker, clone HI30, BD Biosciences, 560976), CD14-BV510 (Monocytes/macrophages, clone M5E2, Biolegend, 301842), CD3-PE (T cells, clone OKT3, Biolegend, 317308), and CD19-V450 (B cells, clone HIB19, Invitrogen eBiosciences, 48019941); ZombieNIR (Biolegend, 423106) was used as a viability marker. Briefly, samples were first washed with warm media, followed by centrifugation and decanting of the solution. Dead cell removal was then performed following the manufacturer’s instructions and live cells were stained with ZombieNIR for 15 minutes at room temperature in the dark. Cells were incubated with Fc Block (Biolegend, 422302) for 10 minutes at 4°C followed by our antibody cocktail for 30 minutes at 4°C in the dark. After each stain, samples were washed with flow buffer (PBS + 2% FBS), centrifuged, and the solution was decanted. When applicable, samples were also stained with Total-Seq-C antibodies (one per patient) following the manufacturer’s instructions (BioLegend, 394661). Samples were resuspended in flow buffer and a BD FACS Melody was used to isolate the ZombieNIR^low^CD45^+^ fraction (into 2ml LoBind Eppendorf tubes (Eppendorf, 022431021). Finally, isolated fractions were used as input for either the Chromium Single-Cell 5’ v2 (10X Genomics, CG000330) or Chromium Next GEM Single Cell V(D)J v1.1 (10X Genomics, CG000207) workflow using the corresponding reagents kits to prepare 5’ single-cell gene expression (scGEX) and BCR (scBCR) cDNA libraries, as per the manufacturer’s instructions. The resulting cDNA libraries were then sequenced on Illumina instruments (NextSeq and NovaSeq) at the sequencing depth recommended by 10X Genomics (30,000 reads/cell for scGEX and 5,000 reads/cell for scBCR).

### Single-cell RNA sequencing data analysis

Raw single-cell RNA sequencing (scRNAseq) data were first processed using cell ranger v.5.0.0 (10X Genomics). The scGEX sequencing data was further analyzed using the Seurat R package. All cell annotations were done at the CD45 annotation level.

The single-cell BCR (scBCR) sequencing data were further analyzed using the Immcantation framework v.4.1.0^109^ in order to define clonal relationships as per the definitions below.

**Clonotype:** one or more cells expressing identical BCR sequences

**Clone:** one or more clonotypes expressing related BCRs with identical V and J genes and CDRH3 lengths with an overall VDJ sequence similarity of approximately 80-90% (threshold for similarity based on Gaussian fit of the data per patient).

Three types of clones were observed: **non-expanded clones** (“singletons”), **expanded clones**, and **clones with lineages.**

**Lineage**: multiple related clonotypes represented by one or more cells per clonotype.

Within lineages, branches were defined as a path starting from the inferred germline to a unique terminal observed clonotype. SHM analysis was conducted using the SHazaM package. The overall methods for performing scRNAseq sample preparation and data analysis are summarized below.

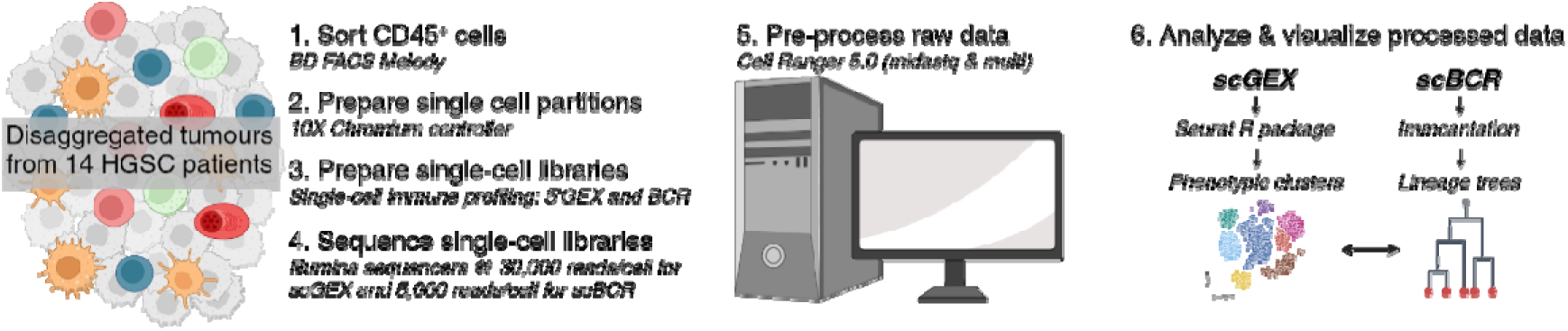

To quantify the total number of mutations for BCRs, the number of mutations per clonotype were only counted once regardless of how many cells that clonotype represented in order to prevent the inflation of mutation counts for highly expanded clonotypes/clones.

### B cell clone selection for BCR-based recombinant antibody production

TIL-B clones were selected for the production of rAbs based on 1) level of expansion and 2) consensus to the lineage. Within the scBCRseq dataset, clones representing a total of 5 cells or more were produced as rAbs. Some of the TIL-B clones that met this expansion threshold had detectable clonal lineages (i.e., were clonally related to at least one other BCR clonotype), in which case the individual clonotype that represented the greatest number of cells among the clonal lineage was selected for rAb production. In cases where multiple clonotypes were equally expanded and each represented the greatest number of cells, the clonotype that was closest to the consensus of the lineage, as measured by the Hamming distance^110^ was selected for rAb production. Expanded TIL-B clones were defined as clones with two or more cells expressing clonally related BCRs. Expanded clonotypes were defined as clonotypes with two or more cells expressing identical BCRs. Lineages, also referred to as clones or clonal groupings, were defined as groupings of two or more clonotypes expressing related BCRs.

### Producing TIL-B BCRs as recombinant antibodies

We generated human IgG1 backbone plasmid vectors based on previously published vectors from Tiller et al^57^ that were acquired from Addgene (AbVec2.0-IGHG1, AbVec1.1-IGLC2-XhoI, AbVec1.1-IGKC) and subsequently modified in-house. The IgG heavy chain backbone vector was modified to contain the full-length constant domain of the human IgG heavy chain fused to a C-terminal triple FLAG tag (3XFLAG) followed by an attenuated internal ribosome entry site (IRES) and a Zeocin resistance gene. The two isotype versions of the IgG light chain backbone vectors were modified to include an attenuated IRES downstream of the constant domain followed by a Puromycin resistance gene. The selection of an attenuated IRES and these antibiotic resistance genes was based on previously published research^111,112^ that identified these components as the optimal strategy for balance between human IgG antibody yield and purity, while simultaneously allowing us to establish antibiotic-resistant stably transfected cell lines producing TIL-B rAbs of high interest. All molecular cloning design was performed using Geneious Prime software (version 2025.0.3, https://www.geneious.com).

Customized IgG backbone vectors were molecularly cloned in-house and subsequently onboarded with Twist Bioscience. BCR variable domains from selected TIL-B clones were molecularly cloned into the appropriate pair of IgG backbone vectors by Twist Bioscience using the restriction sites indicated in the plasmids to generate paired vectors encoding full-length human IgG antibodies.

Paired vectors encoding full-length TIL-B-derived rAb sequences were co-transfected into HEK293T cells using Lipofectamine3000 (Thermo Fisher, L3000015) according to the manufacturer’s instructions. Transfection was confirmed by proxy by assessing the fluorescence of GFP and mCherry control plasmids that were co-transfected into control cells. Culture media (containing secreted rAb from cells co-transfected with paired antibody-encoding plasmids) was harvested from transfected cells at 3-and 6-days post-transfection. To generate antibiotic-resistant stably transfected rAb producing cell lines, cells were harvested with 0.25% Trypsin-PBS following the last media harvest on day 6-post transfection and split to sub-100% confluency and grown in media containing 0.25 ug/ml Puromycin and 15 ug/ml Zeocin for approximately two weeks or until the GFP and mCherry co-transfected control cells were completely killed off.

Antibodies were purified using anti-FLAG magnetic beads (Millipore Sigma, M8823). Cut-off (i.e., wide-bore) pipette tips were used for all steps involving beads to avoid damaging bead complexes. Beads were first equilibrated to remove the storage solution by washing three times with TBS-T. Harvested media for each sample was incubated with 250 ul (125 ul bead volume) of anti-FLAG magnetic beads overnight at 4°C with end-over-end rotation. Following this incubation, magnetic beads were washed three times with TBS-T and purified antibody was eluted from the beads by competitive elution with excess 3XFLAG peptide (Thermo Fisher, A36805) for 30 minutes with end-over-end rotation. Purified antibodies were quantified by Bradford assay and analyzed for purity on a silver-stained non-reducing SDS-PAGE gel.

Antibody purity was assessed by quantifying a lack of protein in the sample purified from media harvested from GFP and mCherry co-transfected control cells by Bradford assay as well as by observing a lack of protein bands in the corresponding lane on the silver-stained reducing SDS-PAGE gel (since no rAb should be present in that sample, suggesting that any protein in the other samples is indeed pure rAb). Proper assembly of rAbs was confirmed by observing a dominant band at ∼150 kDa in the corresponding lanes on the silver-stained non-reducing SDS-PAGE gel comparable to a polyclonal human IgG positive control run in a separate lane.

### Cell culture

A list of all cell lines used and their culture conditions are summarized in the table below.

All cells were cultured in 5% CO_2_ at 37°C.

| Cell line | Tissue type | Source | Culture media |
| --- | --- | --- | --- |
| HEK293T | Human embryonic kidney cells | American Type Culture Collection (ATCC) | Dulbecco's modified Eagle's medium (DMEM, Gibco, 11965092) supplemented with 10% fetal bovine serum (FBS) |
| iOSE397 | Normal ovarian secretory epithelial cells | UBC, Vancouver, Canada | 1:1 mix of Media 199 (Millipore sigma, M4530-500ML) and Media 105 (Millipore sigma, 117-500) supplemented with 5% FBS |
| iOSE7576 | Normal ovarian secretory epithelial cells | UBC, Vancouver, Canada | 1:1 mix of Media 199 (Millipore sigma, M4530-500ML) and Media 105 (Millipore sigma, 117-500) supplemented with 5% FBS |
| OVCAR3 | HGSC | David Bowtell laboratory | Roswell Park Memorial Institute (RPMI) medium 1640 (ATCC formulation, Gibco, A1049101) supplemented with 20% FBS and 0.1% bovine insulin |
| OVCAR4 | HGSC | David Bowtell laboratory | RPMI medium 1640 (Gibco, 22400089) supplemented with 10% FBS |
| CAOV3 | HGSC | David Bowtell<br>laboratory | Dulbecco's modified Eagle's medium (DMEM) supplemented with 10% FBS |
| JHOS2 | HGSC | David Bowtell<br>laboratory | 500 ml DMEM/Ham F12 (Gibco, 1320033) supplemented with 10% FBS and 0.1 mM non-essential amino acids |
| HeLa | Cervical cancer | ATCC | DMEM (Gibco, 11965092) supplemented with 10% fetal bovine serum (FBS) |
| HT29 | Colorectal cancer | ATCC | MyCoy's 5A medium (Gibco, 36600021) supplemented with 10% FBS |
| Panc1 | Pancreatic cancer | ATCC | DMEM (Gibco, 11965092) supplemented with 10% fetal bovine serum (FBS) |

### Flow cytometry

To evaluate TIL-B-derived rAb antigen reactivity to cell lines by flow cytometry, we stained a panel of three normal and seven malignant cell lines (none of which were related to the cases being studied) with individual TIL-B-derived rAbs and an anti-IgG antibody (Alexa Fluor 647 conjugated polyclonal goat anti-human IgG + IgM (H+L), Jackson Immunoresearch, 109-605-044). Intracellular antigen reactivity was assessed by collecting cells by Trypsin-mediated dissociation followed by fixing and permeabilizing cells (Biolegend True-Nuclear Transcription Factor Buffer Set, 424401), TIL-B-derived rAb staining (10 ug/ml), and finally anti-IgG staining (1.5 ug/ml). Cell surface antigen reactivity was assessed by collecting cells with non-enzymatic dissociation (EDTA-PBS) followed by TIL-B-derived rAb staining and subsequent anti-IgG staining. All incubations were 30 minutes long and followed by two washing steps. All staining and washing was performed with PBS + 2% FBS (flow buffer). After the final washing step, cells were resuspended in flow buffer and analysed using a Cytek Aurora cytometer. Reactive rAbs were identified by assessing the median fold change of the median fluorescence intensity from the anti-IgG antibody for each rAb stained sample to the anti-IgG only sample. For intracellular TIL-B-derived rAb staining, the threshold for positivity was set as a fold change of 1.5 or more relative to the anti-IgG only stained control, which corresponded to visually distinguishable shifts in flow histograms (**Supplemental Figure S6A**). For cell surface TIL-B-derived rAb staining, the threshold for positivity was set as a fold change of 2.5 or more. To test TIL-B-derived rAb binding to healthy donor peripheral blood mononuclear cells (PBMCs), viably frozen PBMCs were thawed and then stained as follows (with wash steps between every stain): PBMCs were first stained with a viability dye (1:5000 ZombieNIR, Biolegend) then cells were incubated with the manufacturer’s recommended amount of Fc block (5 ul, Fx TruStain, Biolegend) and Monoblock (5 ul, Biolegend) followed by staining with individual TIL-B rAbs and then with a mix of antibodies for CD45, CD3, CD14, CD56, CD19, and an anti-FLAG detection antibody to detect TIL-B rAb binding. Reactive rAbs were identified by assessing the median fold change of the percent positive cells based on the anti-FLAG antibody fluorophore in the cells/singlets/viability-/CD45+ gate relative to the anti-FLAG only stained sample. All 51 rAbs were tested in preliminary experiments with and without Fc block to rule out non-antigen-specific binding via Fc receptors. Putatively positively binding rAbs were further tested tested in additional experiments with more stringent Fc and mono blocking amounts (7.5 and 10 ul, respectively); only rAbs that showed a fold change of 2.6 or more from these experiments were considered truly positive based on visual interpretation of the flow scatter plots..

### Western blotting

Lysis buffer (20 mM Tris, 150 mM NaCl, 1 mM EDTA, 0.1% TritonX-100, 0.05% w/v deoxycholate, 2 mM Na_3_VO_4_, 10 mM β-glycerophosphate) was prepared fresh (deoxycholate, Na_3_VO_4_, and β-Glycerophosphate from powder) and supplemented with 1 cOmplete mini EDTA-free protease inhibitor tablet (Millipore Sigma, 11836170001) per 50 ml. Fresh cell lysates were then prepared from a panel of cell lines (HEK293T, OVCAR3, OVCAR4, CAOV3) by first collecting the cells grown in 15 cm dishes by scraping, pelleting and washing the cells with cold PBS, then resuspending the cells in lysis buffer (∼1 ml per dish). Samples were incubated at 4°C for 30 minutes with end-over-end rotation to allow cell lysis. Samples were then centrifuged at 10,000 rpm for 20 minutes at 4°C to pellet the DNA and the supernatant (protein lysate) was transferred to a fresh tube and quantified by Bradford assay or using the Qubit 4 Fluorometer (Thermo Fisher).

Up to 40 ug of each lysate was loaded into individual lanes of a Bis-Tris 8% SurePAGE gel (Genscript, M00661) and run under reducing conditions to separate the proteins. Proteins were then transferred to PVDF membranes using either the Trans-Blot SD Semi-Dry Transfer Cell (Bio-Rad) or the Power Blotter (Thermo Fisher). After transfer, membranes were blocked with orbital shaking for one hour with 10 ml per membrane Intercept (TBS) blocking buffer (LI-COR, 927-80001) containing 0.1% Tween-20. Following blocking, membranes were incubated with individual TIL-B rAb solutions: 10 ug/ml rAb in Intercept (TBS) blocking buffer (LI-COR, 927-80001) containing 0.1% Tween-20. Sufficient solution was used to cover the entire membrane. Membranes were incubated with rAb solutions overnight in the dark at 4°C with rocking or orbital shaking. The next day, rAb solutions were recovered and membranes were washed three times with TBS containing 0.1% Tween-20 (TBS-T). After washing, membranes were incubated with secondary antibody solutions: IRDye680 or IRDye800 goat anti-human IgG (LI-COR, 926-68078 and 926-32232) diluted 1:15,000 in TBS-T. Sufficient solution was used to cover the entire membrane. Membranes were incubated with antibody solutions for one hour at room temperature in the dark with orbital shaking. Membranes were then washed three times with TBS-T and finally one time with TBS before being imaged using the Odyssey Classic imager (LI-COR). After initial TIL-B rAb probing, membranes were probed with a rabbit anti-β-actin antibody (Cell Signalling Technologies, 8457S, clone D6A89) diluted 1:1000 in Intercept (TBS) blocking buffer (LI-COR, 927-80001) containing 0.1% Tween-20. Membranes were incubated with anti-β-actin antibody solution for one hour overnight in the dark at 4°C with rocking or orbital shaking. The next day, rAb solutions were recovered and membranes were washed three times with TBS containing 0.1% Tween-20 (TBS-T). After washing, membranes were incubated with secondary antibody solutions: IRDye680 or IRDye800 goat anti-rabbit IgG (LI-COR, 926-68071 and 926-32211) diluted 1:15,000 in TBS-T. Sufficient solution was used to cover the entire membrane. Membranes were incubated with antibody solutions for one hour at room temperature in the dark with orbital shaking. Membranes were then washed three times with TBS-T and finally one time with TBS before being imaged using the Odyssey Classic imager (LI-COR).

### Immunohistochemistry with TIL-B-derived rAbs

TIL-B rAb binding to human tissues was assessed by IHC. TMAs were generated in-house from neutral buffered formalin (NBF)-fixed histogel-embedded cell pellets or formalin-fixed paraffin-embedded (FFPE) tissues. Duplicate cores of 0.6 or 1 mm diameter were punched from tissues using a manual tissue microarrayer (Beecher Instruments). A commercial TMA containing normal ovarian tissue as well as ovarian cancer tissue was also purchased from AMSBIO (OV244).

A staining protocol was established using positive and negative control rAbs and then applied to all rAbs. TIL-B rAb binding was detected using an anti-FLAG detection antibody to prevent false positive staining that could result from direct detection of endogenous IgG antibodies. One TIL-B rAb (A-11-m1) that consistently showed no reactivity to any cell lines by flow cytometry or WB was used as a negative control for IHC. Two TIL-B rAbs that showed either high or low reactivity to multiple cell lines by flow cytometry were used as positive controls for IHC. Parameters of the staining protocol were adjusted to optimize positive signal for the two positive control TIL-B rAbs against the cell lines they were expected to bind based on their reactivity as measured by flow cytometry while maintaining negative signal with the negative control TIL-B rAb. Once optimized, the staining protocol was further tested using an internal TMA consisting of a small set of HGSC samples and control tissues (tonsil, liver, placenta), which confirmed positive staining of HGSC samples and showed no staining of control tissues.

Using the established TIL-B rAb IHC protocol, we first tested each individual TIL-B rAb for IHC reactivity using a cell pellet TMA. All rAbs that showed IHC reactivity on the cell pellet TMA were used for staining tissues on further in-house and commercial TMAs.

All reagents used were from Biocare and all steps were performed at room temperature unless otherwise noted. TMA slides were incubated at 37°C overnight, deparaffinized manually, and subjected to antigen retrieval in a decloaking chamber with Diva Decloaker reagent for 15 minutes at 110°C followed by 10 minutes of cooling. Slides were washed with dH_2_O and loaded onto the Intellipath FLX Autostainer to perform endogenous and non-specific blocking using peroxidased-1 (5 minutes) and background sniper (10 minutes), respectively. Slides were manually stained with individual TIL-B-derived rAbs at 10 ug/ml for 30 minutes. Slides were secondarily stained with a mouse anti-FLAG antibody (Millipore Sigma, F180) diluted 1:2000 for 30 minutes. Slides were incubated with Mach 2 mouse-HRP polymer for 30 minutes on the intelliPATH^TM^, followed by diaminobenzidene (DAB) chromogen for 5 minutes. Slides were counterstained with CAT hematoxylin (1:20 dilution) on the intelliPATH^TM^. Finally, slides were removed from the Intellipath, rinsed with dH_2_O, air dried, and cover-slipped with Ecomount.

All slides were imaged using the Vectra imaging system (Akoya Biosciences) to obtain a 20X image for each core. inForm automated image analysis software (Akoya Biosciences) was used to generate component TIFF images compatible with downstream analysis. TIFF images were analysed qualitatively by eye to determine if staining was present. Positive staining was identified as signal from the DAB chromogen above the level seen with samples stained with the negative control TIL-B rAb, which was selected as a TIL-B rAb that consistently showed no staining by any previously used methods (flow cytometry, WB). If staining was present, all images for the corresponding TIL-B rAb were further qualitatively analysed to determine the strength of the staining, which tissue types were recognized, and which cell types were recognized (based on morphology).

### Immunoprecipitation mass spectrometry

All incubations were done at room temperature unless otherwise specified. TIL-B rAbs were prepared for IP by binding to magnetic protein G dynabeads (Thermo Fisher, 10004D).

Cut-off (i.e., wide-bore) pipette tips were used in all steps involving beads to avoid damaging bead complexes. Protein G dynabeads were aliquoted, placed on a magnet to allow beads to pellet, and the storage solution was removed. Beads were incubated with individual TIL-B rAbs at a ratio of 50 ul of beads (25 ul bead volume) to 5 ug of rAb in a volume of 100 ul PBS. TIL-B rAbs were incubated with beads for 30 minutes while rotating end-over-end. Following this incubation, samples were placed on a magnet to allow beads to pellet, and the supernatant was removed. Bead-bound rAbs were then crosslinked using bis(sulfosuccinimidyl)suberate (BS3) (Fisher Scientific, PI21580) according to manufacturer’s instructions. Briefly, 5 ug rAb was crosslinked with 250 ul of 5 mM BS3 for 30 minutes while rotating end-over-end, followed by quenching with 12.5 ul 1M Tris pH 7.5 for 15 minutes while rotating end-over-end. Crosslinked bead-bound rAbs were then washed three times with PBS + 0.02% Tween-20 (PBS-T) and resuspended in PBS to be stored at 4°C overnight prior to IP the following day.

The following day, IP-lysis buffer (20 mM Tris, 150 mM NaCl, 1 mM EDTA, 0.1% TritonX-100, 0.05% w/v deoxycholate, 2 mM Na_3_VO_4_, 10 mM β-glycerophosphate) was prepared fresh (deoxycholate, Na_3_VO_4_, and β-Glycerophosphate from powder) and supplemented with 1 cOmplete mini EDTA-free protease inhibitor tablet (Millipore Sigma, 11836170001) per 50 ml. Fresh cell lysates were prepared by first collecting cells grown in 15 cm dishes by scraping, pelleting and washing the cells with cold PBS, then resuspending the cells in IP-lysis buffer (∼1 ml per dish). Samples were incubated at 4°C for 30 minutes with end-over-end rotation to allow cell lysis. Samples were centrifuged at 10,000 rpm for 20 minutes and 4°C to pellet the DNA and the supernatant (protein lysate) was transferred to a fresh tube and quantified by Bradford assay or using the Qubit 4 Fluorometer (Thermo Fisher). Lysates were pre-cleared with 100 ul protein G dynabeads (50 ul bead volume) per 1 mg of lysate by incubating for 30 minutes with end-over-end rotation. Following pre-clearing, lysates were divided evenly across samples and added to previously prepared crosslinked bead-bound rAbs. 800-1400 ug of lysate was used with 1-2.5 ug of rAb for each IP sample in a total volume of 500-1400 ul. IP samples were incubated overnight at 4°C with end-over-end rotation.

The following day, IP samples were placed on a magnet to allow beads to pellet, and the supernatant was removed. The beads were washed three times with IP-lysis buffer. After this step, different methods were used for antigen discovery with TIL-B rAbs D-1-m1 and B-13-m1.

#### rAb D-1-m1

Since IP-SS analysis revealed a band of interest uniquely in the rAb D-1-m1 + lysate IP output lanes compared to controls, these samples were prepared for IP-MS with the intention of excising the band of interest for MS protein identification. IP samples were eluted with 50 mM glycine pH 2.8 for 2 minutes with end-over-end rotation and then neutralized with 2 ul of 1 M Tris pH 7.5. The entirety of the IP output samples was resolved by SDS-PAGE under reducing conditions and silver-stained (Thermo Fisher, 24600) to visualize protein bands. Gel pieces that corresponded to the band of interest between approximately 100 and 150 kDa were excised from all lanes including controls, destained according to the manufacturer’s instructions, dehydrated, and snap frozen at -80°C.

Gel band pieces were washed with 500 µL of HPLC water for 15 min with 1200 rpm shaking at room temperature. The water was removed and the gel pieces were destained with 100 µL of 30/70 HPLC methanol/HPLC water for 15 min at room temperature with 1200 rpm shaking. The supernatant was removed and the sample was destained with 100 µL 50/50 HPLC methanol/HPLC water for 15 min at room temperature with 1200 rpm shaking. Supernatant was removed and the gel slices were dehydrated in 100 µL acetonitrile (ACN) for 15 min at room temperature with 1200 rpm shaking. Excess ACN was removed and the gel slices were dried in a speedvac . 50 µL 10 mM dithiolthreitol in 50 mM ammonium bicarbonate was added and gel slices were incubated in the dark for 15 min at 50 °C. Samples were cooled to room temperature prior to adding 0.5 µL of 1 M iodoacetamide (IAA) and incubating at room temperature in the dark for 15 minutes. Excess IAA solution was removed and samples were washed with 100 µL HPLC water for 15 min at room temperature with 1200 rpm shaking. Wash solution was removed and the wash step repeated. The second wash solution was removed and gel bands were dehydrated in 100 µL ACN for 15 min at room temperature with 1200 rpm shaking. Excess ACN was removed and samples were speedvac’d to dryness. 10 µL of 0.1 µg/µL trypsin/lysC in 50 mM ammonium bicarbonate pH 7.8 was added to each sample and incubated overnight at 37 °C. Supernatant was removed to a fresh tube and digest was extracted from gel bands with 50 µL of 30% ACN in HPLC water for 15 min at room temperature with 1200 rpm shaking. Supernatants were combined and peptides were extracted from the gel bands with 50 µL 80% ACN in HPLC water for 15 min at room temperature with 1200 rpm shaking. Peptide containing supernatants were combined,dried in a speedvac and stored at -80C until use.

#### rAb B-13-m1

Since pilot IP-SS analysis had revealed a complex banding pattern uniquely in the rAb B-13-m1 + lysate IP output lanes compared to controls, samples were prepared for IP-MS with more stringent washing to try and isolate the proteins with the strongest (and hypothesized most meaningful or important) interactions with the rAb to reduce complexity of the MS output. After initial washing with IP-lysis buffer, samples were additionally washed with increasing concentrations of NaCl (0.25, 0.5, and 1 M). After the final wash with 1 M NaCl, beads (and any remaining rAb-bound proteins) were resuspended in Laemelli elution buffer (0.1 M Dithiothreitol [DTT], 2% SDS, 200 mM Tris-Cl pH 7.5) and incubated at 95°C for 5 minutes.

Following this incubation, samples were placed on the magnet and the eluted supernatant was harvested, snap frozen at -80°C. 0.5 μL of 1M dithiothreitol (DTT) in HPLC water was added to the eluted protein and samples were incubated for 30 min at 60 °C with shaking at 1000 rpm.

Samples were cooled to room temperature and 1.5 μL of 1M iodoacetamide (IAA) in HPLC water was added and the reaction was incubated for 30 min at room temperature in the dark. An additional 1 μL of DTT was added to quench the reaction. Single Pot Solid-Phase Enhanced Sample Preparation (SP3) was performed as previously published^113^. Specifically, SP3 beads (GE Healthcare, CAT#45152105050250, 65152105050250) were homogenized by vortex mixing and 10 μL of each SP3 bead per sample was transferred to a single fresh tube. The tube was placed on a magnet for 30s and the supernatant removed. Beads were washed by pipette mixing in 200 μL of HPLC water and the tubes again placed on a magnet and supernatant discarded. The beads were reconstituted in 10 μL HPLC water per sample. 12 μL of SP3 beads were transferred to a fresh snap-lock Eppendorf tube and the sample added to the beads. An equal volume of ethanol was added to result in a 50% ethanol solution. Samples were incubated at 24 °C with 1000rpm of shaking for 5 minutes to induce binding. Tubes were placed on a magnetic rack and the supernatant removed and discarded. SP3 beads were rinsed three times with 200 μL 80% ethanol, the supernatant discarded each time. 10 μL of a 0.1 ug/μL solution of Trypsin/LysC (Promega, CAT#V5073) in 0.2M HEPES pH 8 was added to the tube, then 90 μL of HPLC water was added and samples were pipette mixed and incubated for 18 hr at 37 °C with 1000 rpm shaking. Tubes were returned to the magnet after digestion and the supernatant recovered.

50% of the supernatant was desalted by stage tipping for MS analysis^114^. Sample was acidified to 1% TFA using 10% TFA in HPLC water and diluted to 100 μL with HPLC water.

P200 tips were packed with 3 punches of C18 Empore Discs (Sigma CAT#66883-U) using a Sonation StageTip centrifuge (Sonation GMBH, QT#20240101, ∼1500xG). Stage Tips were washed with 100 μL of 0.1% trifluoroacetic acid (TFA) in acetonitrile then conditioned with two aliquots of 100 μL 0.1% TFA in water. Sample was loaded onto the StageTip then rinsed with 200 μL 0.1% formic acid (FA) in water. Peptide was eluted in 100 μL 80% acetonitrile, 0.1% FA in water, dried to completion in a speed vac, then reconstituted in 20 μL 0.1% FA in HPLC water.

Analysis of peptides was carried out on an Orbitrap Fusion Tribrid MS platform (Thermo Scientific) by data dependent acquisition. Samples were introduced using an EASY-nLC 1200 system (Thermo Scientific). Columns used for trapping and separations were packed in-house.

Trapping columns were packed in 100 μm internal diameter capillaries to a length of 2.5 mm with C18 beads (Reprosil-Pur, Dr. Maisch, 3 μm particle size). Trapping was carried out for a total volume of 10 μL at a flow of 25 µL/min. After trapping, gradient elution of peptides was performed on a C18 (Reprosil-Pur, Dr. Maisch, 3 μm particle size) column packed in-house to a length of 25 cm in 100 μm internal diameter capillaries with a laser-pulled and fritted electrospray tip. Elution was performed at a flow rate of 450 nl/min. Mobile phase B (80% HPLC acetonitrile and 0.1% FA) was increased from 6-15% in mobile phase A (water and 0.1% FA) over 1 min, to 35% B over 16 min, to 53% B over 5 min, then a 1 min increase to 100% and the column was washed for 7 min. The instrument was operated in positive ion mode with an electrospray voltage of 2.4 kV and ion transfer tube temperature of 325 °C. MS1 profile data were collected in the orbitrap with a resolution of 120k, a mass range of normal, quadrupole isolation over 400-1200 *m/z*. The RF lens was 60% and the AGC target standard in automatic maximum injection time mode with one microscan. MIPS was set to Peptide and the charge state limited to 2-4. Ions were excluded for 60 seconds after 1 observation with a mass tolerance of 10ppm. HCD was performed at a fixed collision energy of 33% in the quadrupole with an isolation window of 1.4 and no isolation offset. Centroid detection was performed in the ion trap over the Normal mass range, auto scan range and maximum injection time, standard AGC target, with 1 microscan. The cycle time was set to 3 seconds.

### Proteomics data analysis

MS/MS were searched against the UniProt reference proteome (20351 sequences, 2021/07/16) using Sequest HT algorithm through the Proteome Discoverer suite (v2.4, Thermo Scientific). Precursor and fragment ion tolerance were set to 20 ppm and 0.6 Da, respectively. Minimum peptide length was set to 6 amino acids (AA) with a maximum of 2 missed cleavages allowed. Full Trypsin specificity was required. Dynamic modifications included Oxidation (+15.995 Da, M), Acetylation (+42.011 Da, N-Term), and static modification included Carbamidomethyl (+57.021 Da, C). Peptide-to-spectrum matches (PSMs) were calculated using Percolator by searching the results against a decoy sequence set; only PSMs with FDR< 1% were retained in the analysis. Raw mass spectrometry proteomics data have been deposited to the ProteomeXchange Consortium via the PRIDE^115^ partner repository with the dataset identifier PXD063509.

Detected proteins were ranked by their spectral and/or peptide counts in samples of interest (TIL-B rAb plus lysate) and candidate target proteins were identified as any protein that was uniquely detected in all replicates of the TIL-B rAb + lysate IP outputs and not detected in any of the control IP outputs, including IPs performed with no rAb, isotype control rAbs, and TIL-B rAbs + IP-lysis buffer (no lysate).

### Phage-based immunoscreening assay (modified SEREX)

A modified version of SEREX (serological analysis of recombinant tumor cDNA expression libraries)^48–50^ was used for all relevant TIL-B rAbs to attempt to identify their target antigens. A cDNA library was made to represent proteins expressed by the cell line that was most frequently recognized by TIL-B rAbs by flow cytometry (HEK293T). First, total RNA was isolated from cultured HEK293T cells using an RNA isolation kit following manufacturer’s instructions (Qiagen, 74104). Next, polyA+ mRNA was isolated using a kit following manufacturer’s instructions (NEB, E7490S). polyA+ mRNA was converted to cDNA by PCR using oligo-dT primers and used as the starting material to generate the cDNA library using the SMART cDNA library preparation kit (Takara Bio, 634901) according to the manufacturer’s instructions. The resulting cDNA library was packaged using MaxPlax lambda packaging extracts and expressed in *E. coli* XL1-blue. The HEK293T phage cDNA library demonstrated a recombination efficiency of 95% with 2.41x10^6^ unique plaque forming units (pfu)/ml, which was amplified to a final titre of 2.864x10^9^ pfu/ml.

Individual or pooled TIL-B rAbs were screened for reactivity to the HEK293T cDNA phage library using a custom immunoscreening protocol similar to a WB protocol. First, *E. coli* XL1-blue were grown in liquid culture (NZCYM broth supplemented with MgSO4 and maltose) with orbital shaking overnight and the following morning were infected with 4x10^3^ pfu/plate HEK293T cDNA phage library. Infected bacteria were overlaid in soft top broth-agar onto broth-agar plates containing tetracycline and left to grow for 6-7 hours at 37°C. After this time point, circular disc nitrocellulose membranes were overlaid onto each plate containing infected agar.

Plates were incubated for another 12-16 hours at 37°C for a total incubation time of 18-22 hours. Following incubation, the formation of protein-containing bacteriophage plaques was confirmed visibly prior to lifting of nitrocellulose membranes and placing membranes into new plates containing TBS-T. All membranes were washed three times with TBS-T with orbital shaking at room temperature. 17 membranes were then incubated with 3 ml each of individual or pooled TIL-B rAbs at a concentration of 1 ug/ml in Intercept (TBS) blocking buffer (LI-COR, 927-80001) containing 0.1% Tween-20. Membranes were incubated with rAb solutions overnight in the dark at 4°C with rocking or orbital shaking. The next day, rAb solutions were recovered and membranes were washed three times with TBS-T. After washing, membranes were incubated with secondary antibody solutions: IRDye680 or IRDye800 goat anti-human IgG (LI-COR, 926-68078 and 926-32232) diluted 1:15,000 in TBS-T. Sufficient solution was used to cover the entire membrane. Membranes were incubated with antibody solutions for one hour at room temperature in the dark with orbital shaking. Membranes were then washed three times with TBS-T and finally one time with TBS before being imaged using the Odyssey Classic imager (LI-COR).

When reactive plaques were observed, the corresponding phage was isolated from the originating agar plate using a clean pipette tip and incubated in lambda dilution buffer overnight at 4°C with orbital shaking. The following day, the isolated phage was used to infect new bacteria to repeat the protocol as described with the same TIL-B rAb(s). Since the phage were relatively amplified/purified through this isolation process, the resulting membrane (thought to contain mainly a single phage cDNA clone in great abundance) was cut into three pieces to incorporate controls: one piece was probed with TBS-T only, one piece was probed with secondary antibody only (anti-IgG only), and the third piece was probed with TIL-B rAb plus secondary antibody. If a signal was uniquely observed on the piece of the membrane probed with the TIL-B rAb, multiple isolated phage plaques were picked from the originated broth-agar plate and prepared for cDNA excision and sequencing.

cDNA was excised from phage according to the manufacturer’s instructions (Takara Bio, 634901). In brief, isolated phage were used to infect *E. coli* BM25.8 which express the cre recombinase required to excise the cDNA insert in the *E. coli* XL1-blue strain resulting in a plasmid containing the cDNA insert of interest that could be propagated and isolated from resulting BM25.8 cultures. Resulting BM25.8 cultures were grown and cDNA plasmids were isolated using a DNA miniprep kit according to manufacturer’s instructions (Promega, A1223). Purified plasmids were sent for Sanger sequencing using the Nanuq platform from Genome Quebec. Resulting sequences were used for NCBI blast searches to determine the identity of isolated cDNA clones.

### RNA and protein expression analysis

We downloaded gene-level expression data from The Cancer Genome Atlas (TCGA, https://www.cancer.gov/tcga) and the Genotype-Tissue Expression (GTEx)^116^, both generated using UCSC’s Toil RNA-seq workflow^106^. This dataset encompasses approximately 20,000 samples covering 26 tumor/normal pairs (**Supplemental Table S11**).

For each tumor/normal pair, we identified differentially expressed genes using the DESeq2 R package setting normal samples as the reference. Genes were considered differentially expressed if their log2(tumor/normal) ≥ log_2_(1.5), their p-value (Wald test) ≤ 0.05 and their adjusted p-value (Benjamini-Hochberg) ≤ 0.05. Finally, for our three genes of interest (*KDM4A*, *NUMA1*, and *DYNLT2B*), we generated summary heatmaps showing their average log_2_ count across tumor/normal pairs obtained using a variance stabilizing transformation, and their log_2_ fold-change (tumor/normal).

### TIL-B Antigen and polyreactivity ELISAs

Antigens were sourced commercially as follows: KDM4a (Signal Chem, K424-31G-50), NuMA (LSBio, LS-G24789-50), bovine serum albumin (BSA) (Thermo Fisher, 23200), glutathione-S-transferase (GST) (Signal Chem, G52-30U). DYNLT2B was not available as a commercial product and thus was custom synthesized by Sino Biological. For polyreactivity ELISAs, antigens were sourced as follows: dsDNA from calf thymus (Sigma, D8515), LPS from E. coli (Sigma, L2637), human insulin (Sigma, I9278). ssDNA was prepared by heating dsDNA at 95°C for 30 minutes.

ELISAs testing all 51 unique TIL-B rAbs to the “Polyreactivity panel” (dsDNA, ssDNA, LPS, insulin) were performed following previously published protocols from the Nussenzweig laboratory^54–58^. With this assay, antibodies are deemed polyreactive if they respond to at least two of four structurally distinct antigens: double-stranded DNA (dsDNA), single-stranded DNA (ssDNA), insulin, and lipopolysaccharide (LPS). Antigens were diluted to 10 ug/ml (dsDNA, ssDNA, LPS) or 5 ug/ml (all protein antigens) in PBS and used to coat the bottom of wells in a 96-well high-binding EIA/RIA polystyrene plate (Costar, 9018) with 50 ul per well overnight at room temperature. The next day, plates were washed three times with 200 ul per well filtered water and blocked for 1-2 hours with 200 ul per well ELISA buffer (PBS with 1 mM EDTA pH 8 and 0.05% Tween-20). Following blocking, plates were washed three times with 200 ul per well filtered water and incubated with 50 ul per well primary antibodies diluted in PBS for 2 hours at room temperature. TIL-B rAbs were tested at 1, 0.25, 0.0625, and 0.015625 ug/ml (4-fold serial dilutions). Following this incubation, wells were washed three times with 200 ul per well filtered water and incubated with 50 ul per well secondary antibody (goat anti-human IgG/IgM HRP-conjugated, Jackson Immunoresearch, 109-035-044) diluted 1:5,000 in ELISA buffer for 1 hour and room temperature. Following this incubation, wells were washed three times with 200 ul per well filtered water and incubated with 100 ul TMB substrate (1-step Ultra TMB substrate, Thermo Fisher, 34028) for 5-10 minutes at room temperature, followed by the addition of 50 ul of stop solution (Invitrogen, CNB0011). After stopping the reaction, the absorbance of each well was read within 10 minutes using the Varioskan LUX plate reader.

For all ELISAs, absorbance readings were done at 450 nm and 570 nm. The absorbance value at each wavelength was corrected by subtracting the average reading from triplicate blank wells at the relevant wavelength. The final corrected absorbance value was calculated by subtracting the blank-corrected 570 nm absorbance from the blank-corrected 450 nm absorbance. A rAb was considered positive for a given antigen if it yielded a corrected absorbance value of 0.5 or greater, as the negative control rAb consistently yielded values below this threshold. mGO53, GD01, and clinical Herceptin rAbs were used as negative controls as per previously published studies^31,54–58^.

### Serum ELISAs

All buffers, TMB substrate, and stop solution were used from a commercially available kit (Invitrogen, CNB0011). Antigens were diluted in carbonate coating buffer (Invitrogen, CB01100) and used to coat the bottom of wells in a 96-well Nunc Maxisorp high-binding polystyrene plate (Thermo Fisher, 442404) at 1 ug/ml in 100 ul per well overnight at 4°C. The following day, plates were washed four times with 300 ul per well 1X wash buffer (Invitrogen, WB01) and blocked for 1 hour with 300 ul per well 1X assay buffer (Invitrogen, DS98200) at 37°C. Following blocking, plates were washed four times with 300 ul per well 1X was buffer and incubated with 100 ul per well serum samples diluted 1:2000 in 1X assay buffer for 1.5 hours at 37°C. Following this incubation, plates were washed four times with 300 ul per well 1X wash buffer and incubated for 1.5 hours at 37°C with 100 ul per well secondary antibody (goat anti-human IgG/IgM HRP-conjugated, Jackson Immunoresearch, 109-035-044) diluted 1:15,000 in 1X assay buffer. Following this incubation, wells were washed four times with 300 ul per well 1X wash buffer and incubated with 200 ul TMB substrate (1-step Ultra TMB substrate, Thermo Fisher, 34028) for 5-10 minutes at 37°C, followed by the addition of 50 ul of stop solution (Invitrogen, CNB0011). After stopping the reaction, the absorbance of each well was read within 10 minutes using the Varioskan LUX plate reader.

15 age- and sex-matched healthy donor serum samples were tested as a control. All absorbance readings were done at 450 nm and 570 nm. The absorbance value at each wavelength were corrected by subtracting the average reading from triplicate blank wells at each of the wavelengths. The final corrected absorbance was calculated by subtracting the blank-corrected 570 nm absorbance from the blank-corrected 450 nm absorbance. A patient serum sample was considered positive for a given antigen if the corrected absorbance value was greater than three times the standard deviation of the healthy donor serum samples.

### Anti-nuclear antibody testing

All patient serum samples were assessed using the highest-tier clinical tests according to the hierarchy of testing for the diagnosis of rheumatic autoimmune disease outlined in below: the ANA^59^ and rheumatoid factor (RF) tests^62^. RF corresponds to anti-IgG antibodies of all isotypes (though mainly IgM) known to drive rheumatoid autoimmune diseases^117^.

All serum samples were tested at both the earliest and latest available time point per patient by ANA and, if negative by ANA, by RF. Serum samples were sent to Vancouver Island Health Authority (VIHA) and tested according to their clinical protocols. In brief, serum samples were tested by IF for binding to Hep-2 cells at serial dilutions starting at 1:40. Since all samples came from patients of over 40 years of age and autoantibody titres are known to rise with age(Meier et al., 2019; Satoh et al., 2012), the threshold for a positive ANA score was set as positive staining at a dilution of 1:80; any positive staining observed at a dilution of 1:40 was scored as “weakly positive”. Results were reported according to the clinical diagnostic outline, including the highest titre at which a positive score was seen for each sample (or no titre if negative) and the staining pattern.

### Visium spatial transcriptomics Preparation of Standard Visium Libraries

Tissues were cryosectioned at 8 µm thickness and mounted onto Visium spatial gene expression slides (V1, 10X Genomics). Corresponding gene expression (GEX) and immune receptor (VDJ) libraries were generated following the manufacturer’s standard protocols (CG000239) using the Visium Spatial Gene Expression Slide & Reagent Kit (16 rxn, 1000184).

### Preparation of custom BCR panel

An Xgen lockdown probe kit (IDT, 1080577) with custom oligo pools (IDT) was used to enrich the Visium cDNA pools for BCR and TCR sequences. A probe panel of 125 probes corresponding to Immunoglobulin Light chain and 19 probes corresponding to Immunoglobulin Heavy chains were designed with the IMGT database for use with the kit. Each probe was optimized to ensure high specificity and coverage across the target regions. 500 ng of cDNA was used per library as input. A single pre-hybridization PCR of four cycles was conducted to amplify target regions before hybridization. Unlike standard protocols, universal blockers were omitted during the hybridization step.

### Sequencing

Separate BCR and TCR enriched libraries were quality checked on an Agilent Bioanalyzer, multiplexed and sequenced on a PacBio Sequel II SMRTcell (QB3 Genomics, UC Berkeley Sequencing Centre).

### Bioinformatic Processing of Long Read Sequences

Raw reads were processed using the PacBio IsoSeq pipeline (v3.70). The pipeline was used to identify primer sequences, remove polyA tails, correct barcodes, and remove PCR duplicates. This yielded a set of ∼1.2M deduplicated, high-quality reads. IgBLAST was used to align the deduplicated sequences to germline gene references for immunoglobulins, enabling the identification of V, D, and J gene usage and the reconstruction of CDR3 regions.

To assess BCR sequence distribution across tissue compartments (epithelium, stroma, boundary), only BCR heavy chain sequences were considered. The proportion of BCR heavy chain sequences per compartment were reported as the number of sequences per compartment, whereas the total tissue distribution was reported as the number of spots per compartment. The former was not reported as the number of spots to avoid violating the assumptions of the statistical Chi-square test used to assess significant differences in the frequency of BCR heavy chain sequences across different compartments. For patients with Chi square p-values ≤ 0.05, adjust Pearson residuals were calculated and specific clonal expansion groups were considered significant if they yielded an adjusted Pearson residual above the calculated critical value for each test.

### Mapping of kernel density estimate (KDE) values

The spatial locations of all BCR sequences were treated as a point pattern within the two-dimensional coordinate system of the Visium slide. Each occurrence of a BCR sequence was represented as a single spatial point, with all points collectively forming the input dataset for kernel density mapping. A Gaussian kernel was applied to the point pattern using the *spatstat* package in R to compute density values across the spatial domain. Similar to tissue compartments, the proportion of BCR heavy chain sequences inside and outside BCR hotspots were reported as the number of sequences in either region.

## REFERENCES

1. Adams SF, Levine DA, Cadungog MG, et al. Intraepithelial T Cells and Tumor Proliferation: Impact on the Benefit from Surgical Cytoreduction in Advanced Serous Ovarian Cancer. Cancer. 2009;115(13):2891. doi:10.1002/CNCR.24317

2. Hao J, Yu H, Zhang T, An R, Xue Y. Prognostic impact of tumor-infiltrating lymphocytes in high grade serous ovarian cancer: a systematic review and meta-analysis. Ther Adv Med Oncol. 2020;12. doi:10.1177/1758835920967241

3. Hwang WT, Adams SF, Tahirovic E, Hagemann IS, Coukos G. Prognostic Significance of Tumor-infiltrating T-cells in Ovarian Cancer: a Meta-analysis. Gynecol Oncol. 2012;124(2):192. doi:10.1016/J.YGYNO.2011.09.039

4. McAlpine JN, Porter H, Köbel M, et al. BRCA1 and BRCA2 mutations correlate with TP53 abnormalities and presence of immune cell infiltrates in ovarian high-grade serous carcinoma. Modern Pathology. 2012;25(5):740–750. doi:10.1038/MODPATHOL.2011.211/ATTACHMENT/4A3DCB84-C3C6-4105-A00A-27DE629F94FA/MMC1.DOC

5. Milne K, Köbel M, Kalloger SE, et al. Systematic analysis of immune infiltrates in high-grade serous ovarian cancer reveals CD20, FoxP3 and TIA-1 as positive prognostic factors. PLoS One. 2009;4(7). doi:10.1371/journal.pone.0006412

6. Nersesian S, Arseneau RJ, Mejia JP, et al. Improved overall survival in patients with high-grade serous ovarian cancer is associated with CD16a+ immunologic neighborhoods containing NK cells, T cells and macrophages. Front Immunol. 2023;14:1307873. doi:10.3389/FIMMU.2023.1307873/BIBTEX

7. Boudreau JE. Immune-molecular interactions in high-grade serous ovarian cancer distinguish long-term survivors. J Clin Invest. 2024;134(24). doi:10.1172/JCI184790

8. Bowtell DD, Böhm S, Ahmed AA, et al. Rethinking ovarian cancer II: Reducing mortality from high-grade serous ovarian cancer. Nat Rev Cancer. 2015;15(11):668–679. doi:10.1038/nrc4019

9. Pedersen M, Westergaard MCW, Milne K, et al. Adoptive cell therapy with tumor-infiltrating lymphocytes in patients with metastatic ovarian cancer: a pilot study. Oncoimmunology. 2018;7(12). doi:10.1080/2162402X.2018.1502905

10. Bobisse S, Bianchi V, Tanyi JL, et al. A phase 1 trial of adoptive transfer of vaccine-primed autologous circulating T cells in ovarian cancer. Nature Cancer 2023 4:10. 2023;4(10):1410-1417. doi:10.1038/s43018-023-00623-x

11. Gaillard SL, Secord AA, Monk B. The role of immune checkpoint inhibition in the treatment of ovarian cancer. Gynecol Oncol Res Pract. 2016;3(1):1–14. doi:10.1186/s40661-016-0033-6

12. Zamarin D, Burger RA, Sill MW, et al. Randomized Phase II Trial of Nivolumab Versus Nivolumab and Ipilimumab for Recurrent or Persistent Ovarian Cancer: An NRG Oncology Study. Journal of Clinical Oncology. 2020;38(16):1814. doi:10.1200/JCO.19.02059

13. Martin SD, Brown SD, Wick DA, et al. Low Mutation Burden in Ovarian Cancer May Limit the Utility of Neoantigen-Targeted Vaccines. PLoS One. 2016;11(5). doi:10.1371/JOURNAL.PONE.0155189

14. Sha D, Jin Z, Budczies J, Kluck K, Stenzinger A, Sinicrope FA. Tumor Mutational Burden (TMB) as a Predictive Biomarker in Solid Tumors. Cancer Discov. 2020;10(12):1808. doi:10.1158/2159-8290.CD-20-0522

15. Bell D, Berchuck A, Birrer M, et al. Integrated genomic analyses of ovarian carcinoma. Nature 2011 474:7353. 2011;474(7353):609-615. doi:10.1038/nature10166

16. Ciriello G, Miller ML, Aksoy BA, Senbabaoglu Y, Schultz N, Sander C. Emerging landscape of oncogenic signatures across human cancers. Nat Genet. 2013;45(10):1127. doi:10.1038/NG.2762

17. Priestley P, Baber J, Lolkema MP, et al. Pan-cancer whole-genome analyses of metastatic solid tumours. Nature. 2019;575(7781):210-216. doi:10.1038/S41586-019-1689-Y

18. Zhang AW, McPherson A, Milne K, et al. Interfaces of Malignant and Immunologic Clonal Dynamics in Ovarian Cancer. Cell. 2018;173(7):1755–1769.e22. doi:10.1016/j.cell.2018.03.073

19. Burdett NL, Willis MO, Alsop K, et al. Multiomic analysis of homologous recombination-deficient end-stage high-grade serous ovarian cancer. Nature Genetics 2023 55:3. 2023;55(3):437-450. doi:10.1038/s41588-023-01320-2

20. Jiménez-Sánchez A, Memon D, Pourpe S, et al. Heterogeneous Tumor-Immune Microenvironments among Differentially Growing Metastases in an Ovarian Cancer Patient. Cell. 2017;170(5):927–938.e20. doi:10.1016/J.CELL.2017.07.025

21. Kroeger DR, Milne K, Nelson BH. Tumor-infiltrating plasma cells are associated with tertiary lymphoid structures, cytolytic T-cell responses, and superior prognosis in ovarian cancer. Clinical Cancer Research. 2016;22(12):3005–3015. doi:10.1158/1078-0432.CCR-15-2762

22. Nielsen JS, Sahota RA, Milne K, et al. CD20+ Tumor-Infiltrating Lymphocytes Have an Atypical CD27− Memory Phenotype and Together with CD8+ T Cells Promote Favorable Prognosis in Ovarian Cancer. Clinical Cancer Research. 2012;18(12):3281–3292. doi:10.1158/1078-0432.CCR-12-0234

23. Wouters MCA, Nelson BH. Prognostic significance of tumor-infiltrating B cells and plasma cells in human cancer. Clinical Cancer Research. 2018;24(24):6125–6135. doi:10.1158/1078-0432.CCR-18-1481

24. Truxova I, Kasikova L, Hensler M, et al. Mature dendritic cells correlate with favorable immune infiltrate and improved prognosis in ovarian carcinoma patients. J Immunother Cancer. 2018;6(1). doi:10.1186/S40425-018-0446-3

25. Iglesia MD, Vincent BG, Parker JS, et al. Prognostic B-Cell Signatures using mRNA-Seq in Patients with Subtype-Specific Breast and Ovarian Cancer. Clin Cancer Res. 2014;20(14):3818. doi:10.1158/1078-0432.CCR-13-3368

26. Garsed DW, Pandey A, Fereday S, et al. The genomic and immune landscape of long-term survivors of high-grade serous ovarian cancer. Nature Genetics 2022 54:12. 2022;54(12):1853-1864. doi:10.1038/s41588-022-01230-9

27. Nelson BH, Hamilton PT, Phung MT, et al. Immunological and molecular features of the tumor microenvironment of long-term survivors of ovarian cancer. J Clin Invest. Published online October 29, 2024. doi:10.1172/JCI179501

28. Laumont CM, Nelson BH. B cells in the tumor microenvironment: Multi-faceted organizers, regulators, and effectors of anti-tumor immunity. Cancer Cell. 2023;41(3):466–489. doi:10.1016/J.CCELL.2023.02.017

29. Laumont CM, Banville AC, Gilardi M, Hollern DP, Nelson BH. Tumour-infiltrating B cells: immunological mechanisms, clinical impact and therapeutic opportunities. Nature Reviews Cancer 2022. Published online April 7, 2022:1-17. doi:10.1038/s41568-022-00466-1

30. Helmink BA, Reddy SM, Gao J, et al. B cells and tertiary lymphoid structures promote immunotherapy response. Nature. 2020;577(7791):549-555. doi:10.1038/s41586-019-1922-8

31. Mazor RD, Nathan N, Gilboa A, et al. Tumor-reactive antibodies evolve from non-binding and autoreactive precursors. Cell. Published online March 15, 2022. doi:10.1016/j.cell.2022.02.012

32. Biswas S, Mandal G, Anadon CM, et al. Targeting intracellular oncoproteins with dimeric IgA promotes expulsion from the cytoplasm and immune-mediated control of epithelial cancers. Immunity. 2023;56(11):2570–2583.e6. doi:10.1016/J.IMMUNI.2023.09.013

33. Biswas S, Mandal G, Payne KK, et al. IgA transcytosis and antigen recognition govern ovarian cancer immunity. Nature. 2021;591(7850):464-470. doi:10.1038/s41586-020-03144-0

34. Chen J, Tan Y, Sun F, et al. Single-cell transcriptome and antigen-immunoglobin analysis reveals the diversity of B cells in non-small cell lung cancer. Genome Biol. 2020;21(1):152. doi:10.1186/s13059-020-02064-6

35. Bruno TC, Ebner PJ, Moore BL, et al. Antigen-presenting intratumoral B cells affect CD4+ TIL phenotypes in non–small cell lung cancer patients. Cancer Immunol Res. 2017;5(10):898–907. doi:10.1158/2326-6066.CIR-17-0075

36. Garaud S, Buisseret L, Solinas C, et al. Tumor-infiltrating B cells signal functional humoral immune responses in breast cancer. Reference information: JCI Insight. 2019;4(18). doi:10.1172/jci.insight.129641

37. Shi JY, Gao Q, Wang ZC, et al. Margin-infiltrating CD20(+) B cells display an atypical memory phenotype and correlate with favorable prognosis in hepatocellular carcinoma. Clin Cancer Res. 2013;19(21):5994–6005. doi:10.1158/1078-0432.CCR-12-3497

38. Ma J, Wu Y, Ma L, et al. A blueprint for tumor-infiltrating B cells across human cancers. Science (1979). 2024;384(6695). doi:10.1126/SCIENCE.ADJ4857/SUPPL_FILE/SCIENCE.ADJ4857_MDAR_REPROD UCIBILITY_CHECKLIST.PDF

39. Hao D, Han G, Sinjab A, et al. The Single-Cell Immunogenomic Landscape of B and Plasma Cells in Early-Stage Lung Adenocarcinoma. Cancer Discov. 2022;12(11):2626–2645. doi:10.1158/2159-8290.CD-21-1658

40. Fitzsimons E, Qian D, Enica A, et al. A pan-cancer single-cell RNA-seq atlas of intratumoral B cells. Cancer Cell. 2024;42(10):1784–1797.e4. doi:10.1016/J.CCELL.2024.09.011

41. Xia J, Xie Z, Niu G, et al. Single-cell landscape and clinical outcomes of infiltrating B cells in colorectal cancer. Immunology. 2023;168(1):135–151. doi:10.1111/IMM.13568

42. Sivakumar S, Jainarayanan A, Arbe-Barnes E, et al. Distinct immune cell infiltration patterns in pancreatic ductal adenocarcinoma (PDAC) exhibit divergent immune cell selection and immunosuppressive mechanisms. Nature Communications 2025 16:1. 2025;16(1):1-20. doi:10.1038/s41467-024-55424-2

43. Qian J, Olbrecht S, Boeckx B, et al. A pan-cancer blueprint of the heterogeneous tumor microenvironment revealed by single-cell profiling. Cell Res. 2020;30(9):745–762. doi:10.1038/s41422-020-0355-0

44. Duan M, Nguyen DC, Joyner CJ, et al. Understanding heterogeneity of human bone marrow plasma cell maturation and survival pathways by single-cell analyses. Cell Rep. 2023;42(7):112682. doi:10.1016/J.CELREP.2023.112682

45. Ma J, Wu Y, Ma L, et al. A blueprint for tumor-infiltrating B cells across human cancers. Science. 2024;384(6695):eadj4857. doi:10.1126/SCIENCE.ADJ4857/SUPPL_FILE/SCIENCE.ADJ4857_MDAR_REPRODUCIBILITY_CHECKLIST.PDF

46. Peperzak V, Xiao Y, Veraar EAM, Borst J. CD27 sustains survival of CTLs in virus-infected nonlymphoid tissue in mice by inducing autocrine IL-2 production. Journal of Clinical Investigation. 2010;120(1):168–178. doi:10.1172/JCI40178

47. Cheng RYH, de Rutte J, Ito CEK, et al. SEC-seq: association of molecular signatures with antibody secretion in thousands of single human plasma cells. Nature Communications 2023 14:1. 2023;14(1):1-15. doi:10.1038/s41467-023-39367-8

48. Nesslinger NJ, Ng A, Tsang KY, et al. A viral vaccine encoding PSA induces antigen spreading to a common set of self proteins in prostate cancer patients. Clin Cancer Res. 2010;16(15):4046. doi:10.1158/1078-0432.CCR-10-0948

49. Sahin U, Türeci Ö, Schmitt H, et al. Human neoplasms elicit multiple specific immune responses in the autologous host. Proc Natl Acad Sci U S A. 1995;92(25):11810–11813. doi:10.1073/PNAS.92.25.11810

50. Stone B, Schummer M, Paley PJ, et al. Serologic analysis of ovarian tumor antigens reveals a bias toward antigens encoded on 17q. Int J Cancer. 2003;104(1):73–84. doi:10.1002/IJC.10900

51. Taimen P, Viljamaa M, Kallajoki M. Preferential Expression of NuMA in the Nuclei of Proliferating Cells. Exp Cell Res. 2000;256(1):140–149. doi:10.1006/EXCR.2000.4799

52. Chen Y, Yang S, Yu T, et al. KDM4A promotes malignant progression of breast cancer by down-regulating BMP9 inducing consequent enhancement of glutamine metabolism. Cancer Cell Int. 2024;24(1):322. doi:10.1186/S12935-024-03504-0

53. Zhao J, Li B, Ren Y, et al. Histone demethylase KDM4A plays an oncogenic role in nasopharyngeal carcinoma by promoting cell migration and invasion. Exp Mol Med. 2021;53(8):1207. doi:10.1038/S12276-021-00657-0

54. Wardemann H, Yurasov S, Schaefer A, Young JW, Meffre E, Nussenzweig MC. Predominant autoantibody production by early human B cell precursors. Science (1979). 2003;301(5638):1374-1377. doi:10.1126/SCIENCE.1086907/SUPPL_FILE/WARDEMANN.SOM.PDF

55. Meffre E, Schaefer A, Wardemann H, Wilson P, Davis E, Nussenzweig MC. Surrogate Light Chain Expressing Human Peripheral B Cells Produce Self-reactive Antibodies. J Exp Med. 2004;199(1):145–150. doi:10.1084/jem.20031550

56. Tiller T, Tsuiji M, Yurasov S, Velinzon K, Nussenzweig MC, Wardemann H. Autoreactivity in Human IgG+ Memory B Cells. Immunity. 2007;26(2):205–213. doi:10.1016/j.immuni.2007.01.009

57. Tiller T, Meffre E, Yurasov S, Tsuiji M, Nussenzweig MC, Wardemann H. Efficient generation of monoclonal antibodies from single human B cells by single cell RT-PCR and expression vector cloning. J Immunol Methods. 2007;329(1-2):112–124. doi:10.1016/j.jim.2007.09.017

58. Scheid JF, Mouquet H, Kofer J, Yurasov S, Nussenzweig MC, Wardemann H. Differential regulation of self-reactivity discriminates between IgG + human circulating memory B cells and bone marrow plasma cells. Proc Natl Acad Sci U S A. 2011;108(44):18044–18048. doi:10.1073/pnas.1113395108

59. Kavanaugh A, Tomar R, Reveille J, Solomon DH, Homburger HA. Guidelines for Clinical Use of the Antinuclear Antibody Test and Tests for Specific Autoantibodies to Nuclear Antigens. Arch Pathol Lab Med. 2000;124(1):71–81. doi:10.5858/2000-124-0071-GFCUOT

60. Tebo AE. Recent approaches to optimize laboratory assessment of antinuclear antibodies. Clinical and Vaccine Immunology. 2017;24(12). doi:10.1128/CVI.00270-17,

61. Liu F, Wang XQ, Zou JW, Li M, Pan CC, Si YQ. Association between serum antinuclear antibody and rheumatoid arthritis. Front Immunol. 2024;15. doi:10.3389/FIMMU.2024.1358114/PDF

62. Ingegnoli F, Castelli R, Gualtierotti R. Rheumatoid factors: Clinical applications. Dis Markers. 2013;35(6):727–734. doi:10.1155/2013/726598,

63. Mariz HA, Sato EI, Barbosa SH, Rodrigues SH, Dellavance A, Andrade LEC. Pattern on the antinuclear antibody-HEp-2 test is a critical parameter for discriminating antinuclear antibody-positive healthy individuals and patients with autoimmune rheumatic diseases. Arthritis Rheum. 2011;63(1):191–200. doi:10.1002/ART.30084

64. Deng Y, Tan Y, Zhou D, et al. Single-Cell RNA-Sequencing Atlas Reveals the Tumor Microenvironment of Metastatic High-Grade Serous Ovarian Carcinoma. Front Immunol. 2022;13:1. doi:10.3389/FIMMU.2022.923194/FULL

65. Olalekan S, Xie B, Back R, Eckart H, Basu A. Characterizing the tumor microenvironment of metastatic ovarian cancer by single-cell transcriptomics. Cell Rep. 2021;35(8). doi:10.1016/J.CELREP.2021.109165

66. Kasikova L, Rakova J, Hensler M, et al. Tertiary lymphoid structures and B cells determine clinically relevant T cell phenotypes in ovarian cancer. Nature Communications 2024 15:1. 2024;15(1):1-19. doi:10.1038/s41467-024-46873-w

67. Germain C, Gnjatic S, Tamzalit F, et al. Presence of B cells in tertiary lymphoid structures is associated with a protective immunity in patients with lung cancer. Am J Respir Crit Care Med. 2014;189(7):832–844. doi:10.1164/rccm.201309-1611OC

68. Meylan M, Petitprez F, Becht E, et al. Tertiary lymphoid structures generate and propagate anti-tumor antibody-producing plasma cells in renal cell cancer. Immunity. 2022;55(3):527–541.e5. doi:10.1016/j.immuni.2022.02.001

69. Mirlekar B, Wang Y, Li S, et al. Balance between immunoregulatory B cells and plasma cells drives pancreatic tumor immunity. Cell Rep Med. 2022;3(9). doi:10.1016/J.XCRM.2022.100744

70. Ghosh D, Jiang W, Mukhopadhyay D, Mellins ED. New insights into B cells as antigen presenting cells. Curr Opin Immunol. 2021;70:129–137. doi:10.1016/J.COI.2021.06.003

71. Rastogi I, Jeon D, Moseman JE, Muralidhar A, Potluri HK, McNeel DG. Role of B cells as antigen presenting cells. Front Immunol. 2022;13:954936. doi:10.3389/FIMMU.2022.954936

72. Possamaï D, Pagé G, Panès R, Gagnon É, Lapointe R. CD40L-Stimulated B Lymphocytes Are Polarized toward APC Functions after Exposure to IL-4 and IL-21. The Journal of Immunology. 2021;207(1):77–89. doi:10.4049/JIMMUNOL.2001173,

73. Shen P, Fillatreau S. Antibody-independent functions of B cells: a focus on cytokines. Nat Rev Immunol. 2015;15(7):441–451. doi:10.1038/NRI3857

74. Furtado GC, Marinkovic T, Martin AP, et al. Lymphotoxin beta receptor signaling is required for inflammatory lymphangiogenesis in the thyroid. Proc Natl Acad Sci U S A. 2007;104(12):5026–5031. doi:10.1073/PNAS.0606697104

75. Luther SA, Lopez T, Bai W, Hanahan D, Cyster JG. BLC expression in pancreatic islets causes B cell recruitment and lymphotoxin-dependent lymphoid neogenesis. Immunity. 2000;12(5):471–481. doi:10.1016/S1074-7613(00)80199-5

76. Wing E, Sutherland C, Miles K, et al. Double-negative-2 B cells are the major synovial plasma cell precursor in rheumatoid arthritis. Front Immunol. 2023;14. doi:10.3389/FIMMU.2023.1241474/FULL

77. Wang Y, Li R, Tong R, et al. Integrating single-cell RNA and T cell/B cell receptor sequencing with mass cytometry reveals dynamic trajectories of human peripheral immune cells from birth to old age. Nat Immunol. 2025;26(2):308–322. doi:10.1038/S41590-024-02059-6

78. Rosser EC, Mauri C. Regulatory B cells: origin, phenotype, and function. Immunity. 2015;42(4):607–612. doi:10.1016/J.IMMUNI.2015.04.005

79. Catalán D, Mansilla MA, Ferrier A, et al. Immunosuppressive Mechanisms of Regulatory B Cells. Front Immunol. 2021;12. doi:10.3389/FIMMU.2021.611795

80. Yao M, Preall J, Yeh J, et al. Plasma cells in human pancreatic ductal adenocarcinoma secrete antibodies to self-antigens. JCI Insight. Published online September 26, 2023. doi:10.1172/JCI.INSIGHT.172449

81. Montes CL, Acosta-Rodríguez E V, Merino MC, Bermejo DA, Gruppi A. Polyclonal B cell activation in infections: infectious agents’ devilry or defense mechanism of the host? J Leukoc Biol. 2007;82(5):1027–1032. doi:10.1189/JLB.0407214

82. Hansen MH, Nielsen H, Ditzel HJ. The tumor-infiltrating B cell response in medullary breast cancer is oligoclonal and directed against the autoantigen actin exposed on the surface of apoptotic cancer cells. Proceedings of the National Academy of Sciences. 2001;98(22):12659–12664. doi:10.1073/pnas.171460798

83. Andrade F, Roy S, Nicholson D, Thornberry N, Rosen A, Casciola-Rosen L. Granzyme B directly and efficiently cleaves several downstream caspase substrates: implications for CTL-induced apoptosis. Immunity. 1998;8(4):451–460. doi:10.1016/S1074-7613(00)80550-6

84. Hansen JE, Tse CM, Chan G, Heinze ER, Nishimura RN, Weisbart RH. Intranuclear protein transduction through a nucleoside salvage pathway. Journal of Biological Chemistry. 2007;282(29):20790–20793. doi:10.1074/jbc.C700090200

85. Guo K, Li J, Tang JP, et al. Targeting intracellular oncoproteins with antibody therapy or vaccination. Sci Transl Med. 2011;3(99). doi:10.1126/SCITRANSLMED.3002296

86. Wang L, Chang J, Varghese D, et al. A small molecule modulates Jumonji histone demethylase activity and selectively inhibits cancer growth. Nat Commun. 2013;4. doi:10.1038/NCOMMS3035

87. Brignole C, Bensa V, Fonseca NA, et al. Cell surface Nucleolin represents a novel cellular target for neuroblastoma therapy. Journal of Experimental and Clinical Cancer Research. 2021;40(1):1–13. doi:10.1186/S13046-021-01993-9/FIGURES/5

88. Rafiq K, Bergtold A, Clynes R. Immune complex–mediated antigen presentation induces tumor immunity. Journal of Clinical Investigation. 2002;110(1):71–79. doi:10.1172/jci15640

89. Lou H, Ling GS, Cao X. Autoantibodies in systemic lupus erythematosus: From immunopathology to therapeutic target. J Autoimmun. 2022;132. doi:10.1016/J.JAUT.2022.102861,

90. Crescioli S, Correa I, Ng J, et al. B cell profiles, antibody repertoire and reactivity reveal dysregulated responses with autoimmune features in melanoma. Nat Commun. 2023;14(1). doi:10.1038/S41467-023-39042-Y

91. Tsuiji M, Yurasov S, Velinzon K, Thomas S, Nussenzweig MC, Wardemann H. A checkpoint for autoreactivity in human IgM+ memory B cell development. J Exp Med. 2006;203(2):393. doi:10.1084/JEM.20052033

92. Labombarde JG, Pillai MR, Wehenkel M, et al. Induction of broadly reactive influenza antibodies increases susceptibility to autoimmunity. Cell Rep. 2022;38(10):110482. doi:10.1016/J.CELREP.2022.110482

93. Reyes-Ruiz A, Dimitrov JD. How can polyreactive antibodies conquer rapidly evolving viruses? Trends Immunol. 2021;42(8):654–657. doi:10.1016/J.IT.2021.06.008

94. Ravi P, Freeman D, Thomas J, et al. Comprehensive multiplexed autoantibody profiling of patients with advanced urothelial cancer. J Immunother Cancer. 2024;12(2). doi:10.1136/JITC-2023-008215

95. Arcani R, Bertin D, Bardin N, et al. Anti-NuMA antibodies: clinical associations and significance in patients with primary Sjögren’s syndrome or systemic lupus erythematosus. Rheumatology (Oxford). 2021;60(9):4074–4084. doi:10.1093/RHEUMATOLOGY/KEAA881

96. Heegaard NHH, West-Nørager M, Tanassi JT, et al. Circulating antinuclear antibodies in patients with pelvic masses are associated with malignancy and decreased survival. PLoS One. 2012;7(2). doi:10.1371/JOURNAL.PONE.0030997

97. Vlagea A, Falagan S, Gutiérrez-Gutiérrez G, et al. Antinuclear antibodies and cancer: A literature review. Crit Rev Oncol Hematol. 2018;127:42–49. doi:10.1016/J.CRITREVONC.2018.05.002

98. Vallerskog T, Gunnarsson I, Widhe M, et al. Treatment with rituximab affects both the cellular and the humoral arm of the immune system in patients with SLE. Clin Immunol. 2007;122(1):62–74. doi:10.1016/J.CLIM.2006.08.016

99. Lee DSW, Rojas OL, Gommerman JL. B cell depletion therapies in autoimmune disease: advances and mechanistic insights. Nat Rev Drug Discov. 2021;20(3):179–199. doi:10.1038/S41573-020-00092-2

100. Høydahl LS, Richter L, Frick R, et al. Plasma Cells Are the Most Abundant Gluten Peptide MHC-expressing Cells in Inflamed Intestinal Tissues From Patients With Celiac Disease. Gastroenterology. 2019;156(5):1428–1439.e10. doi:10.1053/J.GASTRO.2018.12.013

101. Wang AA, Luessi F, Neziraj T, et al. B cell depletion with anti-CD20 promotes neuroprotection in a BAFF-dependent manner in mice and humans. Sci Transl Med. 2024;16(737). doi:10.1126/SCITRANSLMED.ADI0295

102. Bermejo DA, Jackson SW, Gorosito-Serran M, et al. Trypanosoma cruzi trans-sialidase initiates an ROR-γt–AHR-independent program leading to IL-17 production by activated B cells. Nat Immunol. 2013;14(5):514. doi:10.1038/NI.2569

103. Shen P, Roch T, Lampropoulou V, et al. IL-35-producing B cells are critical regulators of immunity during autoimmune and infectious diseases. Nature. 2014;507(7492):366. doi:10.1038/NATURE12979

104. Mietzner B, Tsuiji M, Scheid J, et al. Autoreactive IgG memory antibodies in patients with systemic lupus erythematosus arise from nonreactive and polyreactive precursors. Proceedings of the National Academy of Sciences. 2008;105(28):9727–9732. doi:10.1073/PNAS.0803644105

105. Grivennikov SI, Greten FR, Karin M. Immunity, Inflammation, and Cancer. Cell. 2010;140(6):883. doi:10.1016/J.CELL.2010.01.025

106. Vivian J, Rao AA, Nothaft FA, et al. Toil enables reproducible, open source, big biomedical data analyses. Nature Biotechnology 2017 35:4. 2017;35(4):314-316. doi:10.1038/nbt.3772

107. Love MI, Huber W, Anders S. Moderated estimation of fold change and dispersion for RNA-seq data with DESeq2. Genome Biol. 2014;15(12):1–21. doi:10.1186/S13059-014-0550-8/FIGURES/9

108. Schmidt U, Weigert M, Broaddus C, Myers G. Cell Detection with Star-convex Polygons. Lecture Notes in Computer Science (including subseries Lecture Notes in Artificial Intelligence and Lecture Notes in Bioinformatics). 2018;11071 LNCS:265–273. doi:10.1007/978-3-030-00934-2_30

109. Gupta NT, Vander Heiden JA, Uduman M, Gadala-Maria D, Yaari G, Kleinstein SH. Change-O: a toolkit for analyzing large-scale B cell immunoglobulin repertoire sequencing data. Bioinformatics. 2015;31(20):3356–3358. doi:10.1093/BIOINFORMATICS/BTV359

110. Ralph DK, Matsen FA. Using B cell receptor lineage structures to predict affinity. PLoS Comput Biol. 2020;16(11). doi:10.1371/JOURNAL.PCBI.1008391

111. Spidel JL, Vaessen B, Chan YY, Grasso L, Kline JB. Rapid high-throughput cloning and stable expression of antibodies in HEK293 cells. J Immunol Methods. 2016;439:50–58. doi:10.1016/j.jim.2016.09.007

112. Ho SC, Bardor M, Feng H, et al. IRES-mediated Tricistronic vectors for enhancing generation of high monoclonal antibody expressing CHO cell lines. J Biotechnol. 2012;157:130–139. doi:10.1016/j.jbiotec.2011.09.023

113. Asleh K, Lluch A, Goytain A, et al. Triple-Negative PAM50 Non-Basal Breast Cancer Subtype Predicts Benefit from Extended Adjuvant Capecitabine. Clinical Cancer Research. 2023;29(2):389–400. doi:10.1158/1078-0432.CCR-22-2191,

114. Rappsilber J, Mann M, Ishihama Y. Protocol for micro-purification, enrichment, pre-fractionation and storage of peptides for proteomics using StageTips. Nat Protoc. 2007;2(8):1896–1906. doi:10.1038/NPROT.2007.261,

115. Perez-Riverol Y, Bandla C, Kundu DJ, et al. The PRIDE database at 20 years: 2025 update. Nucleic Acids Res. 2025;53(D1):D543–D553. doi:10.1093/NAR/GKAE1011,

116. Lonsdale J, Thomas J, Salvatore M, et al. The Genotype-Tissue Expression (GTEx) project. Nat Genet. 2013;45(6):580–585. doi:10.1038/ng.2653

117. Tiwari V, Jandu JS, Bergman MJ. Rheumatoid Factor. Measuring Immunity: Basic Science and Clinical Practice. Published online July 24, 2023:187–192. doi:10.1016/B978-012455900-4/50276-2

