## Supplementary figures and images for "Prognostically favorable immune responses to ovarian cancer are distinguished by self-reactive intra-epithelial plasma cells"

### Supplemental Figure S1

Supplemental Figure S1

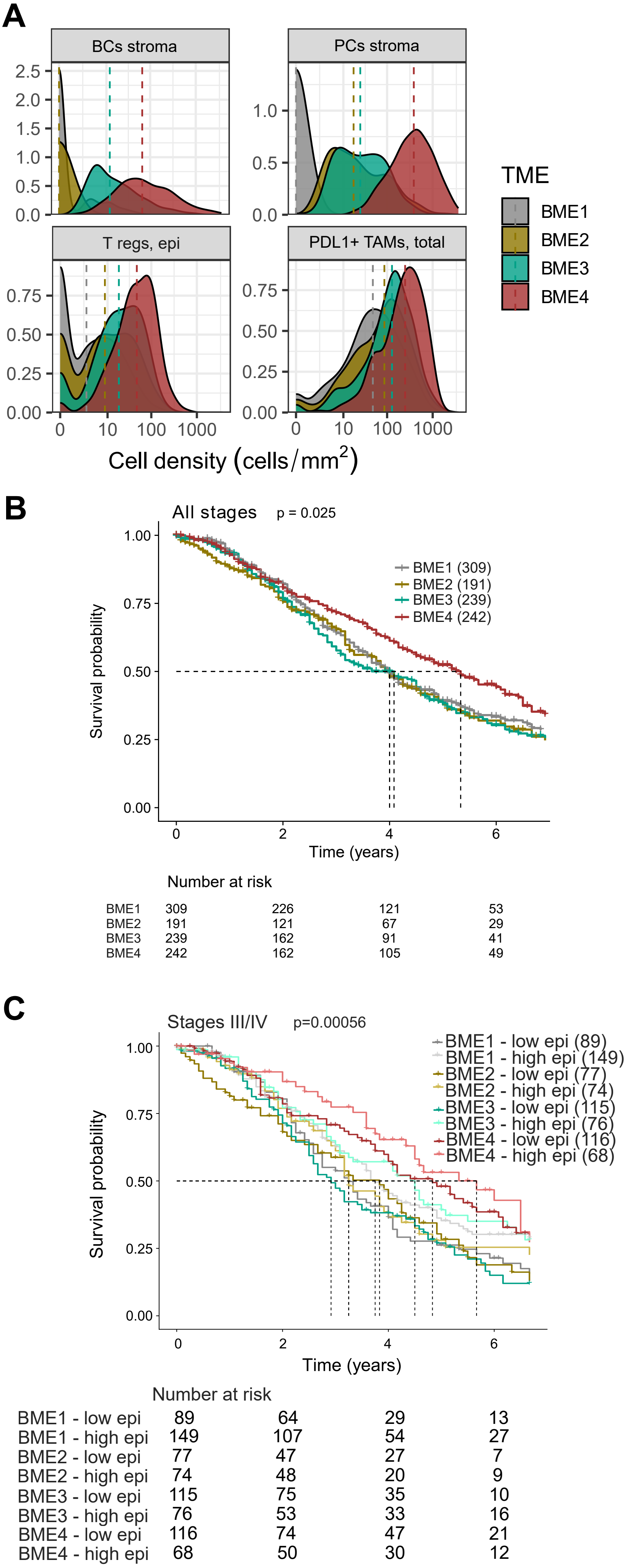

### Supplemental Figure S2

Supplemental Figure S2

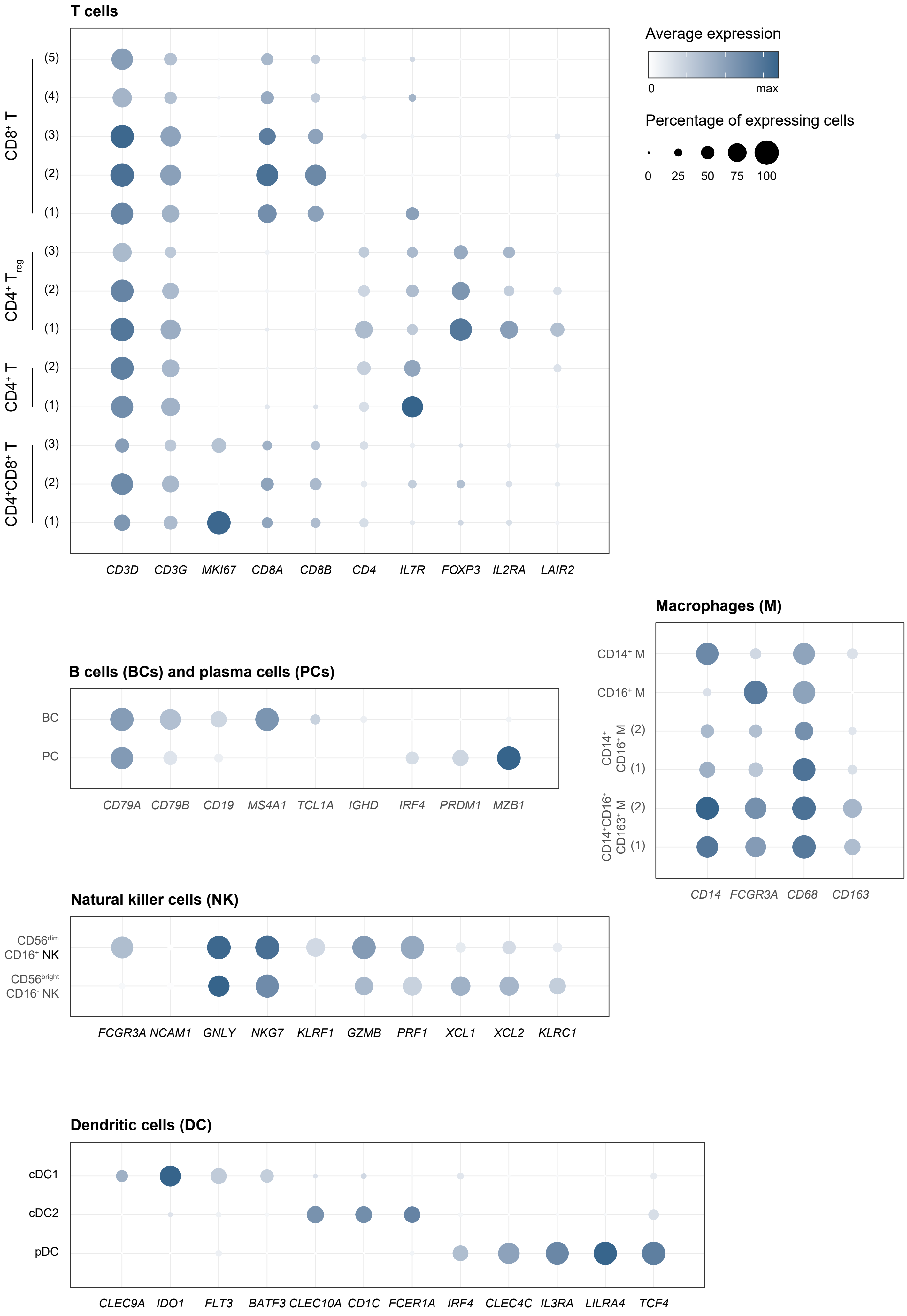

### Supplemental Figure S3

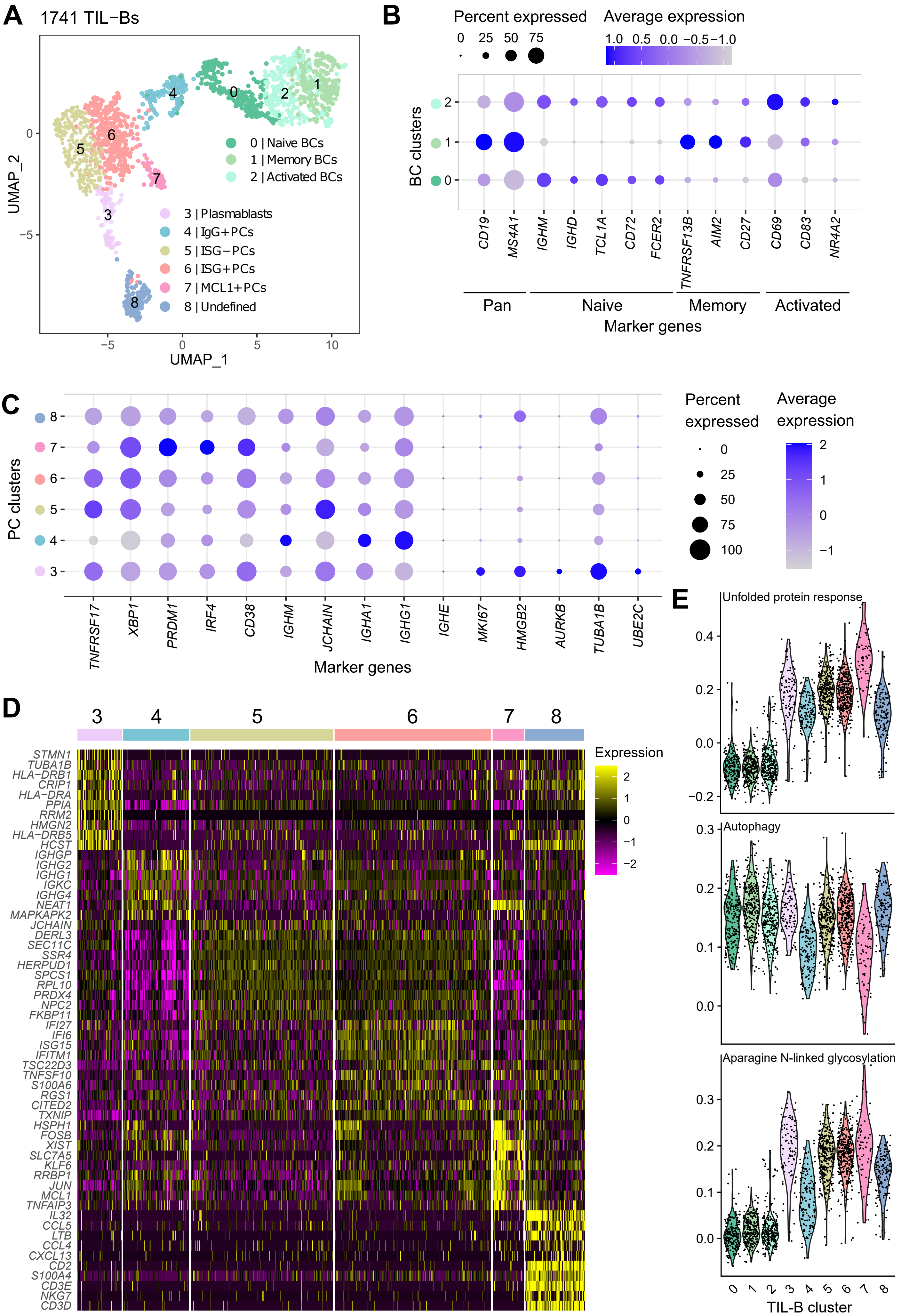

### Supplemental Figure S4

Supplemental Figure S4

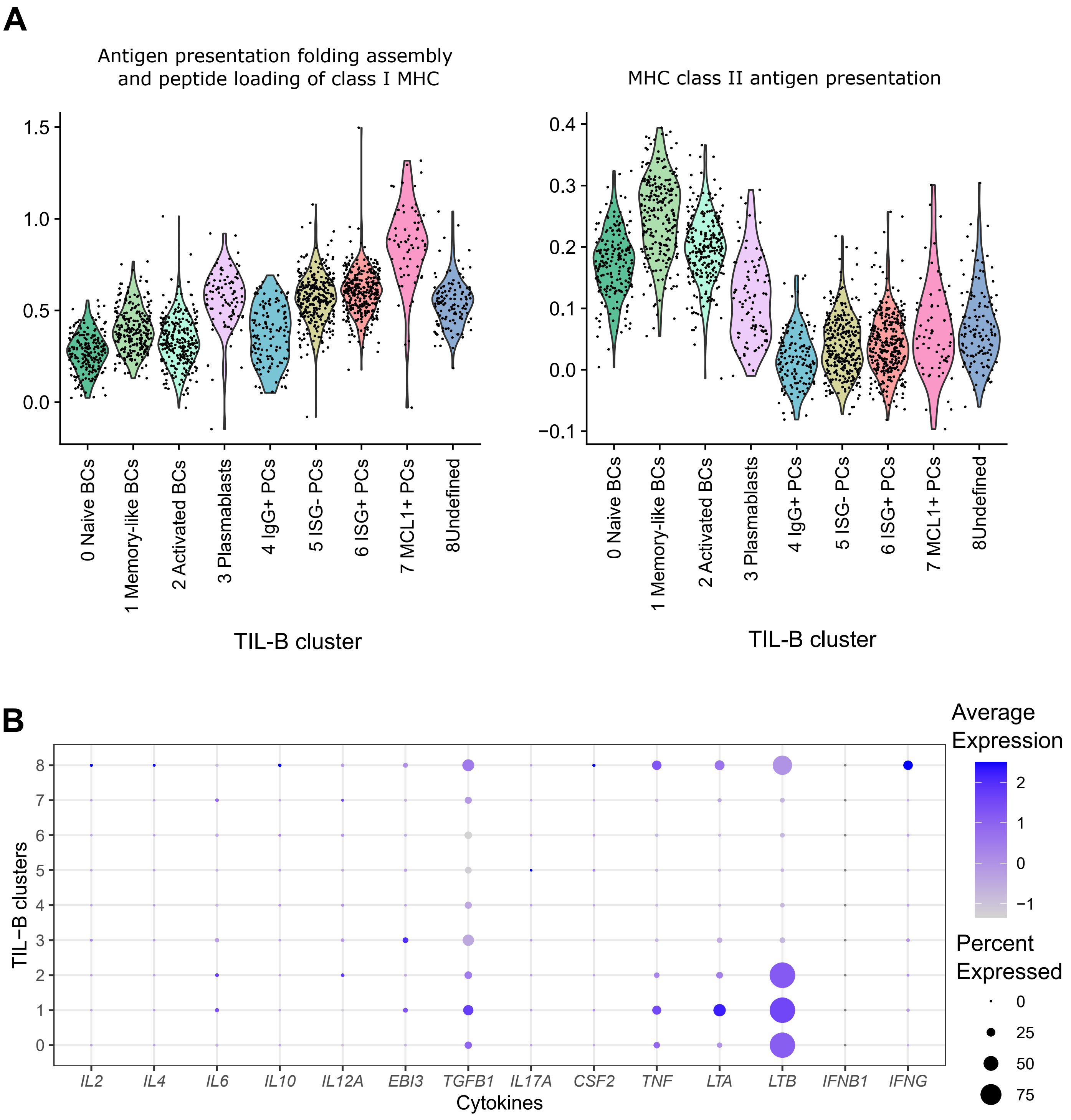

### Supplemental Figure S5

Supplemental Figure S5

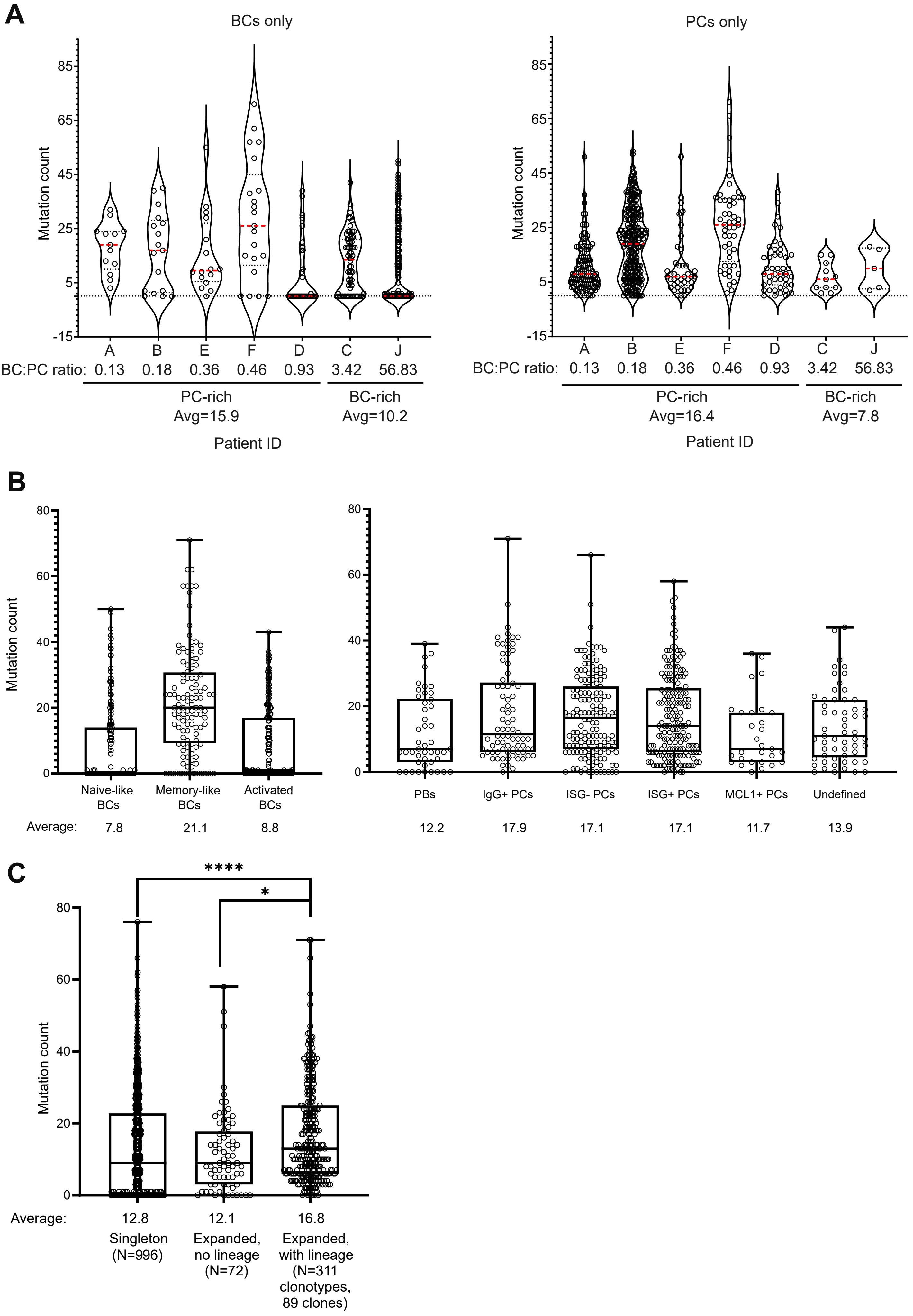

### Supplemental Figure S6

Supplemental Figure S6

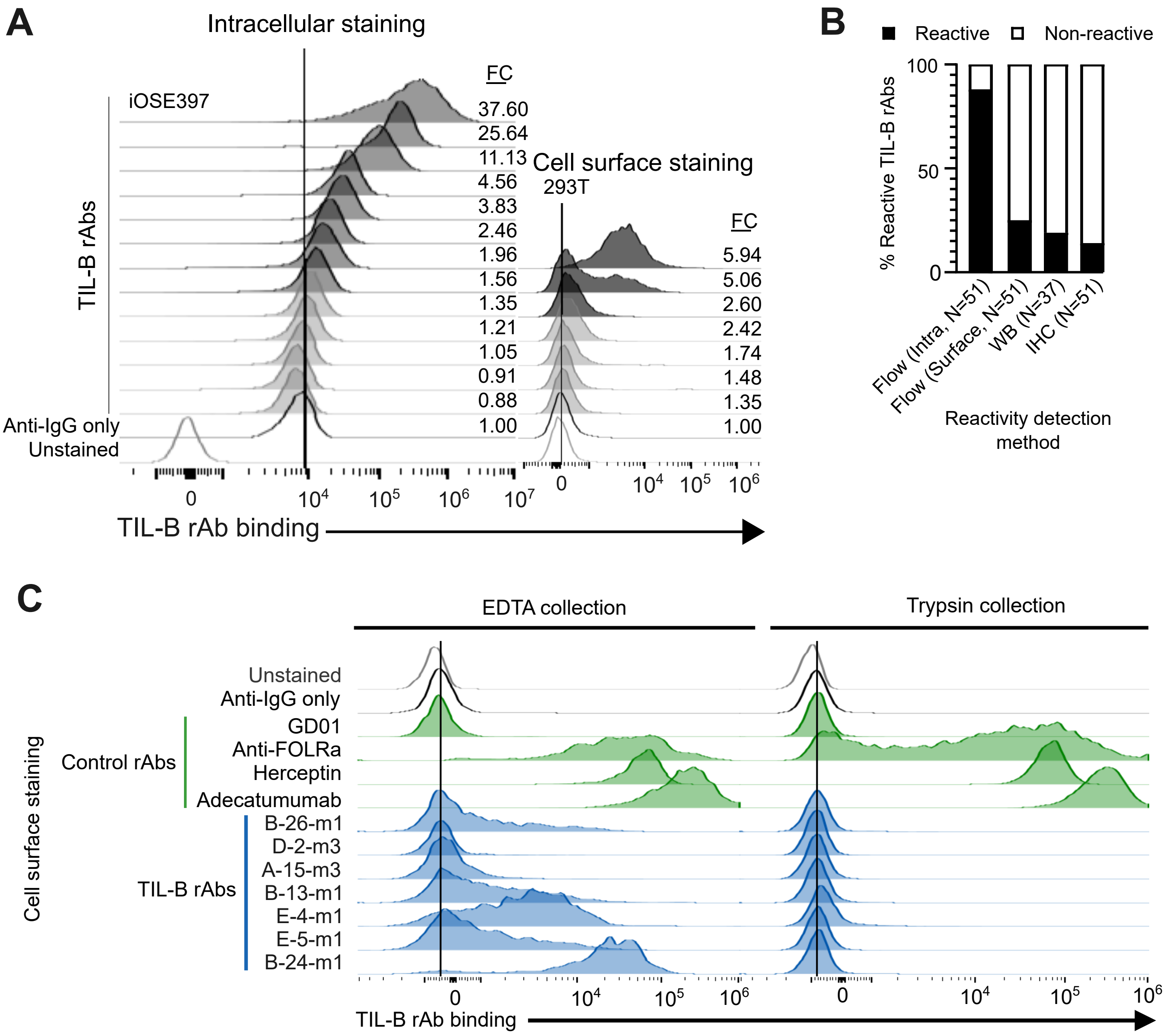

### Supplemental Figure S7

#### Supplemental Figure S7

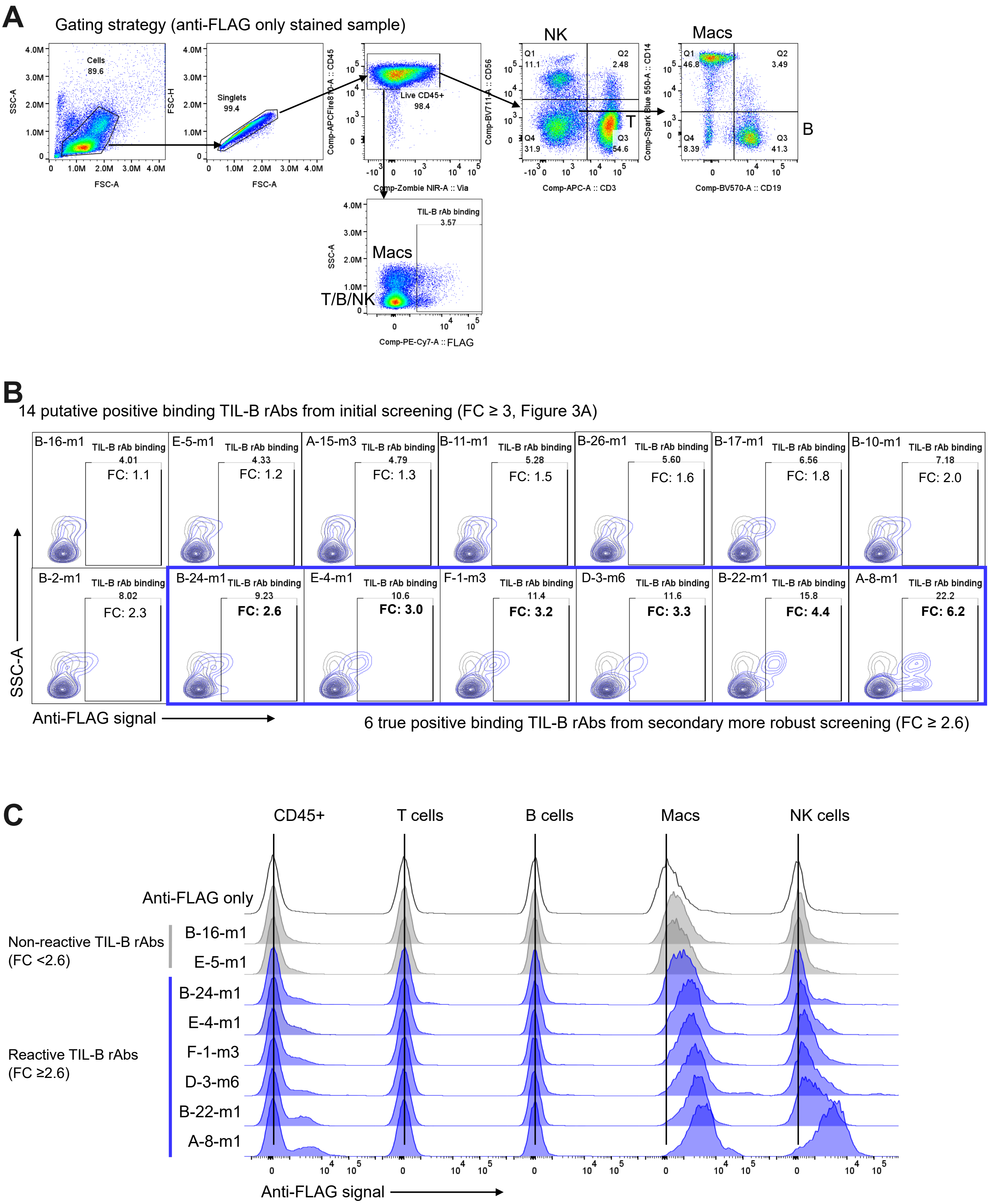

### Supplemental Figure S8

Supplemental Figure S8

A

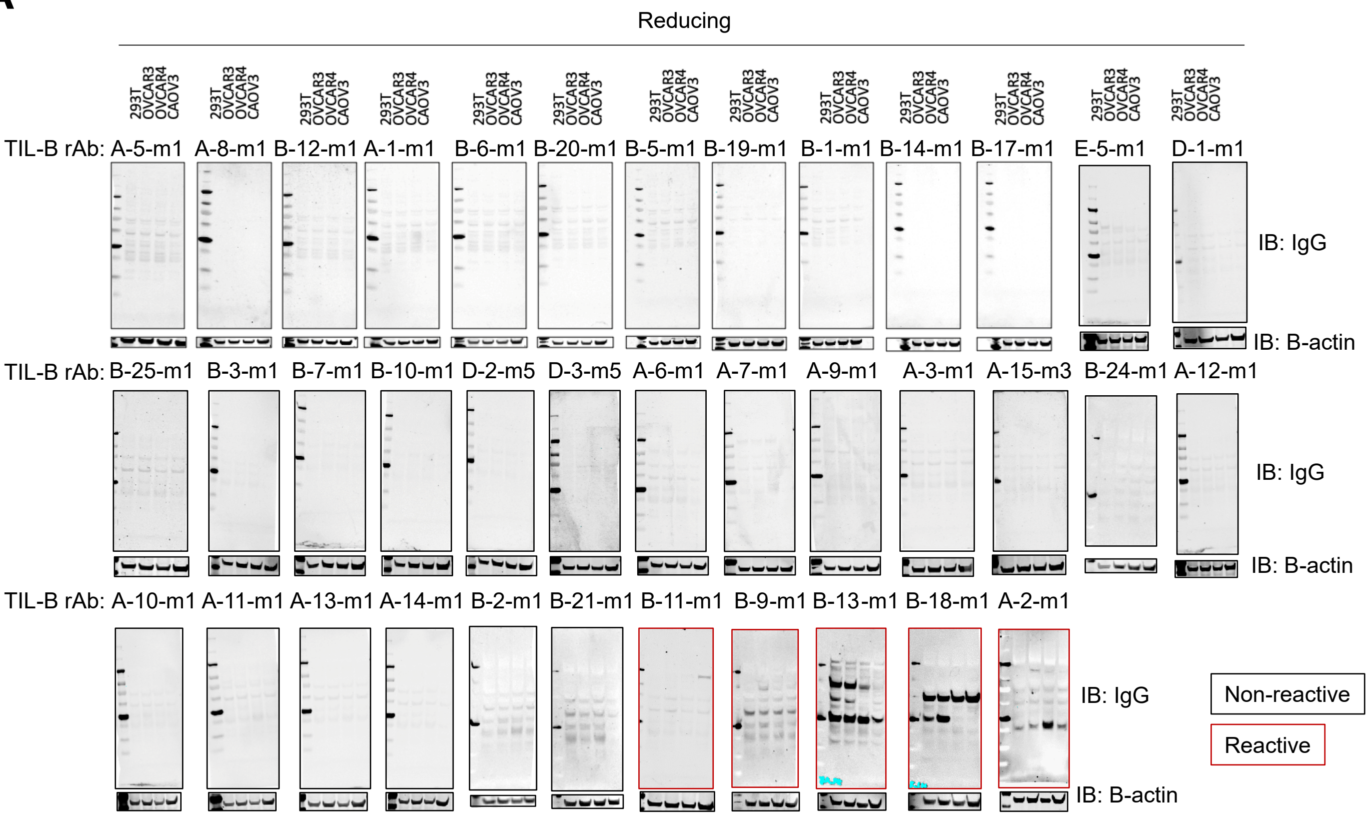


B

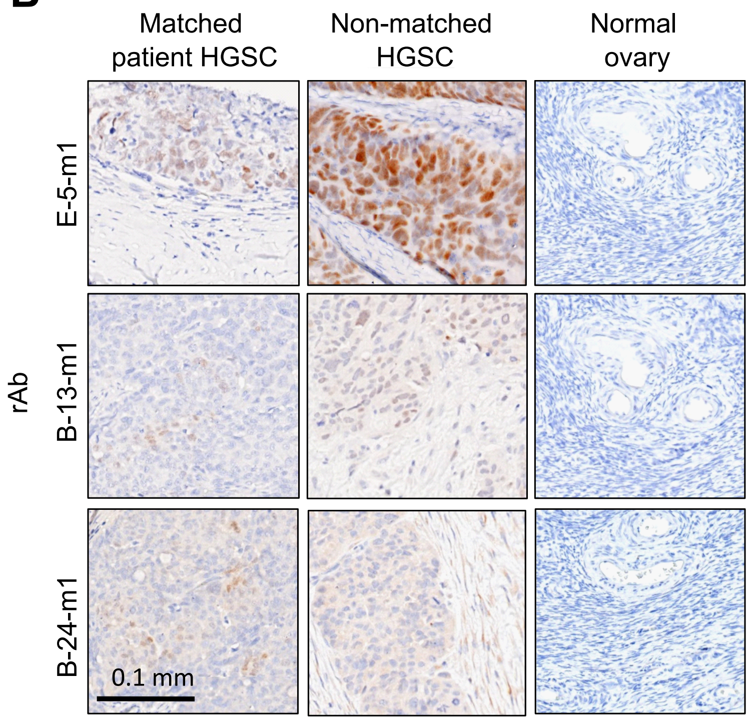

### Supplemental Figure S9

#### Supplemental Figure S9

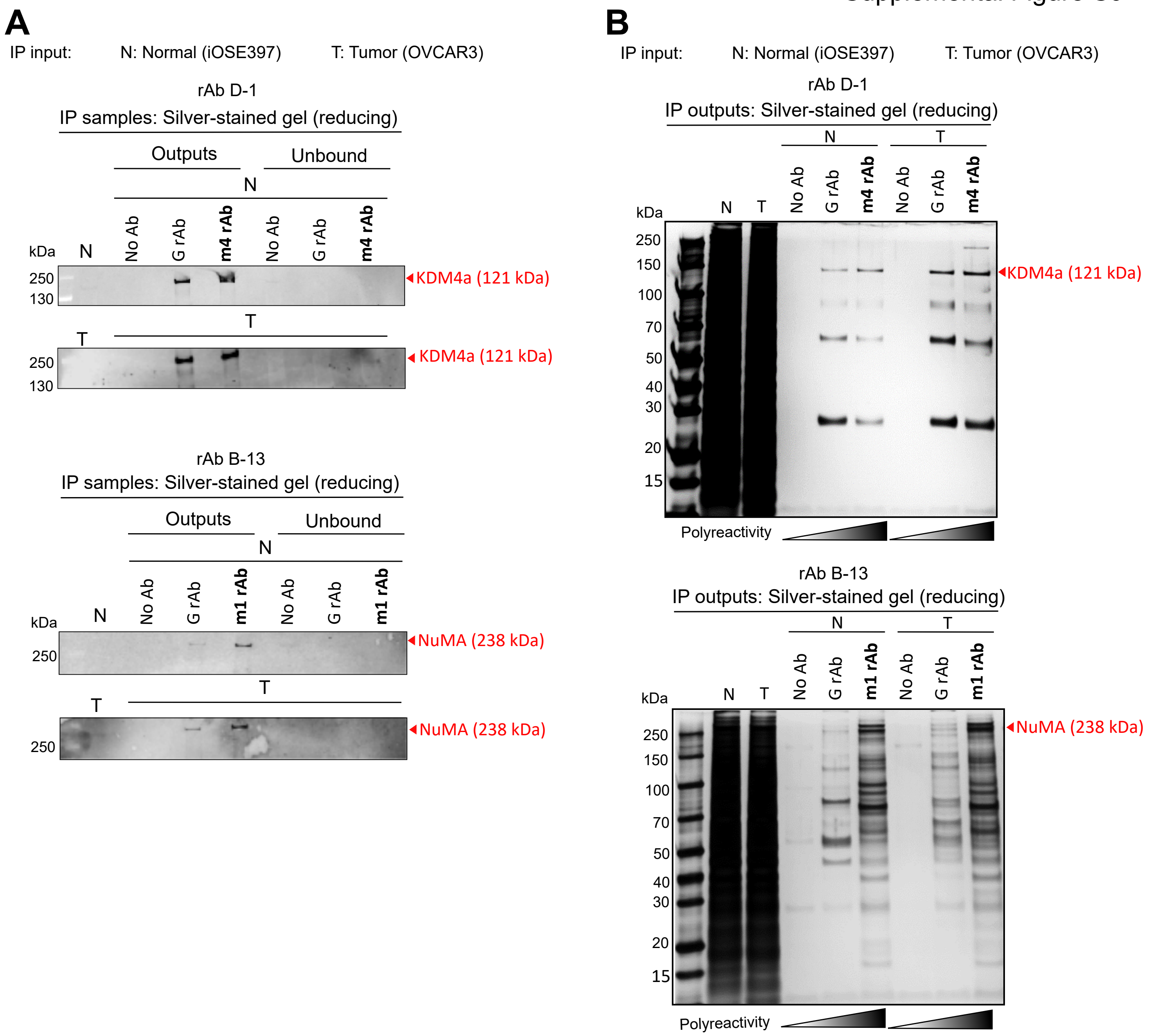

### Supplemental Figure S10

Supplemental Figure S10

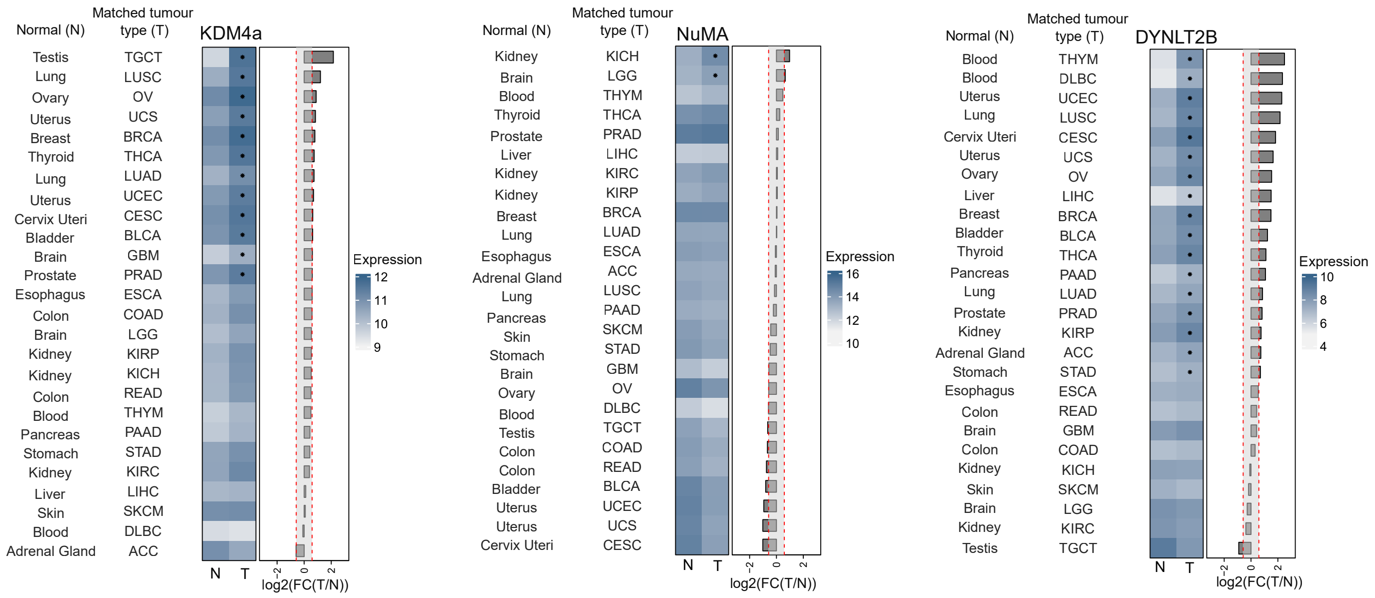

### Supplemental Figure S11

Supplemental Figure S11

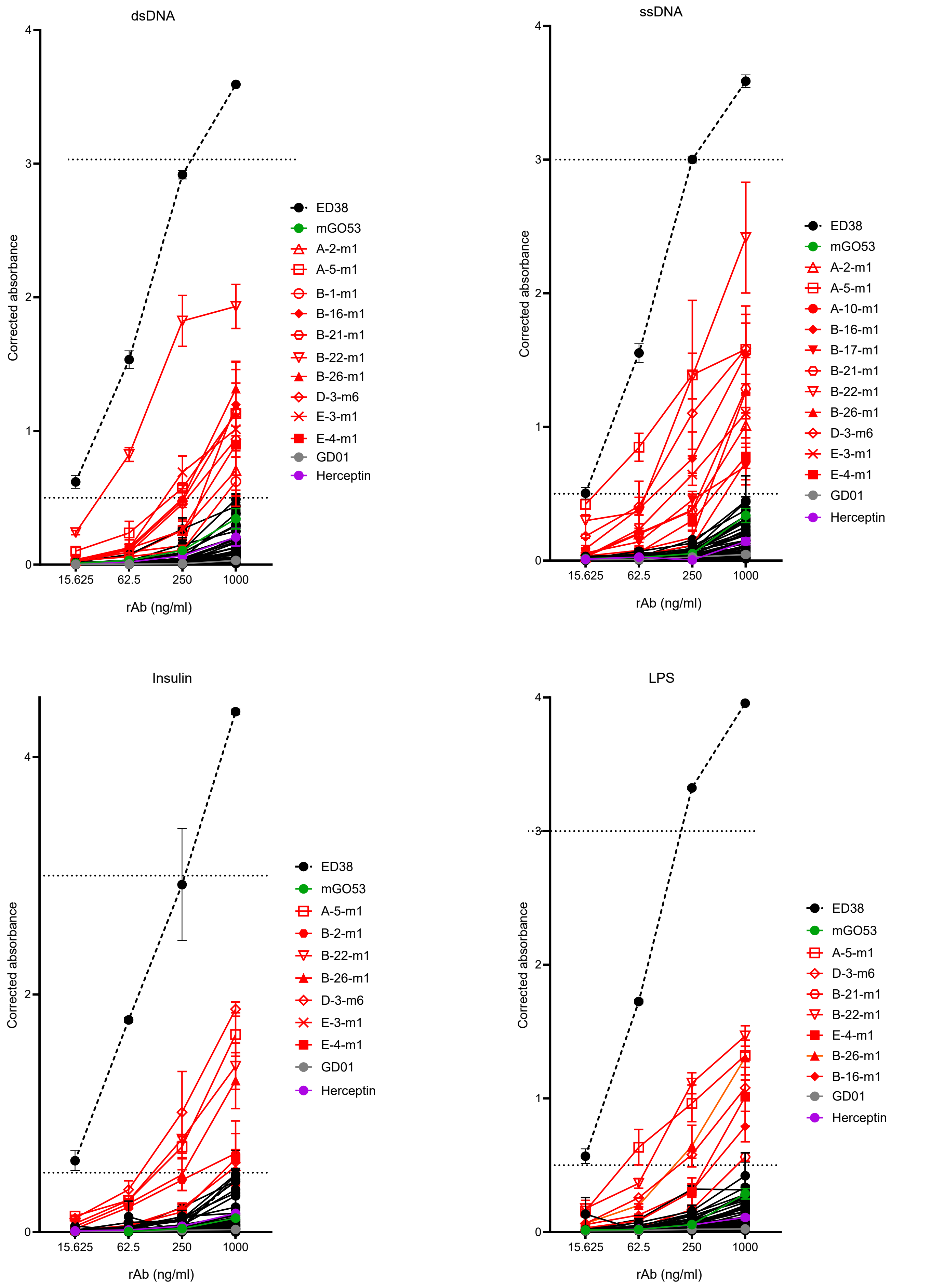

### Supplemental Figure S12

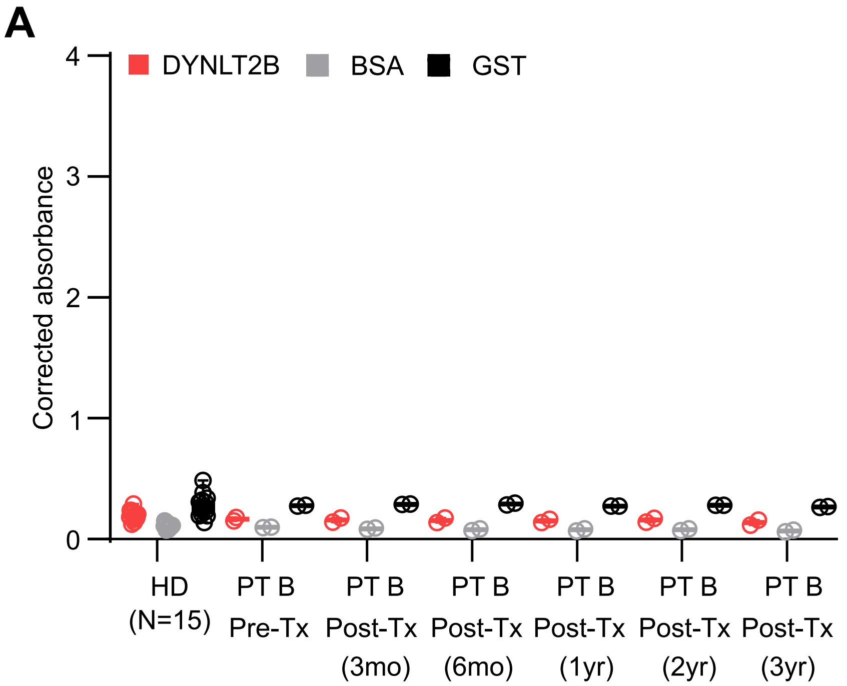

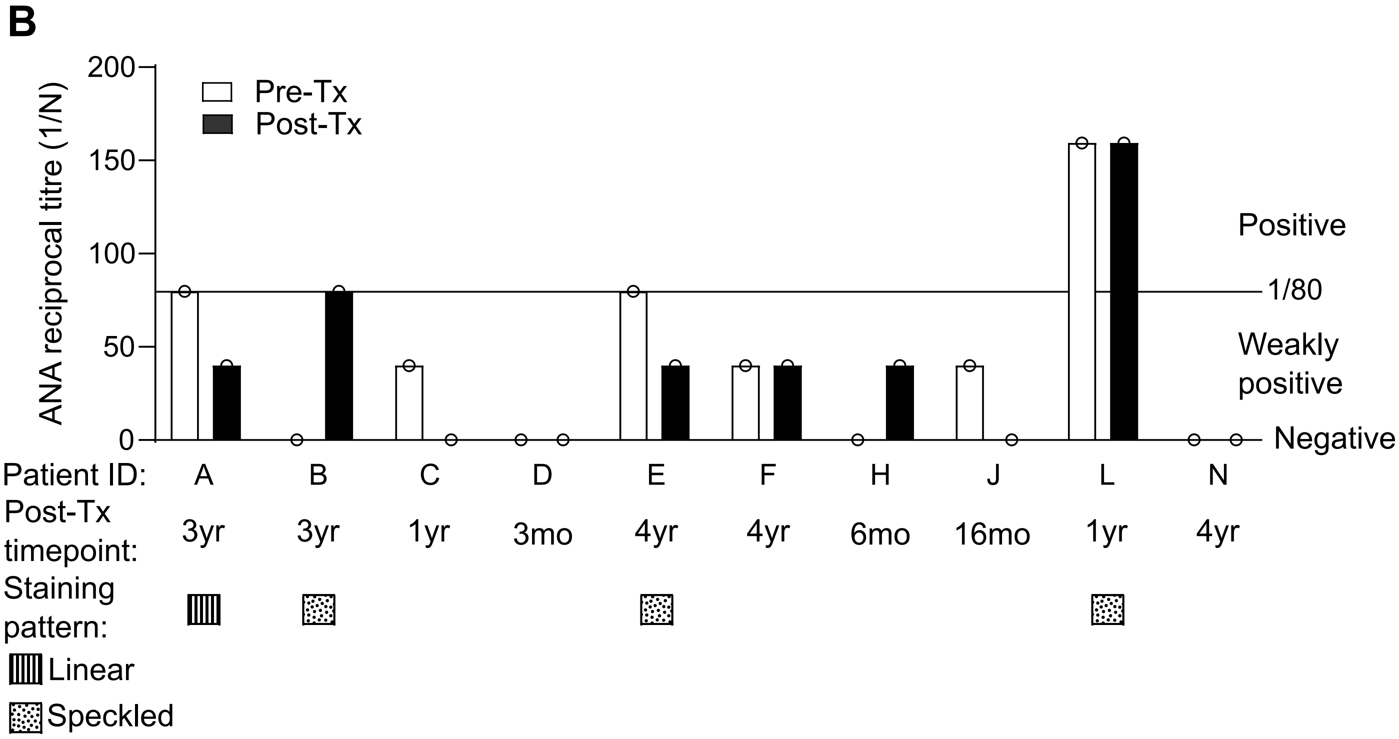

### Supplemental Figure S13

Supplemental Figure S13

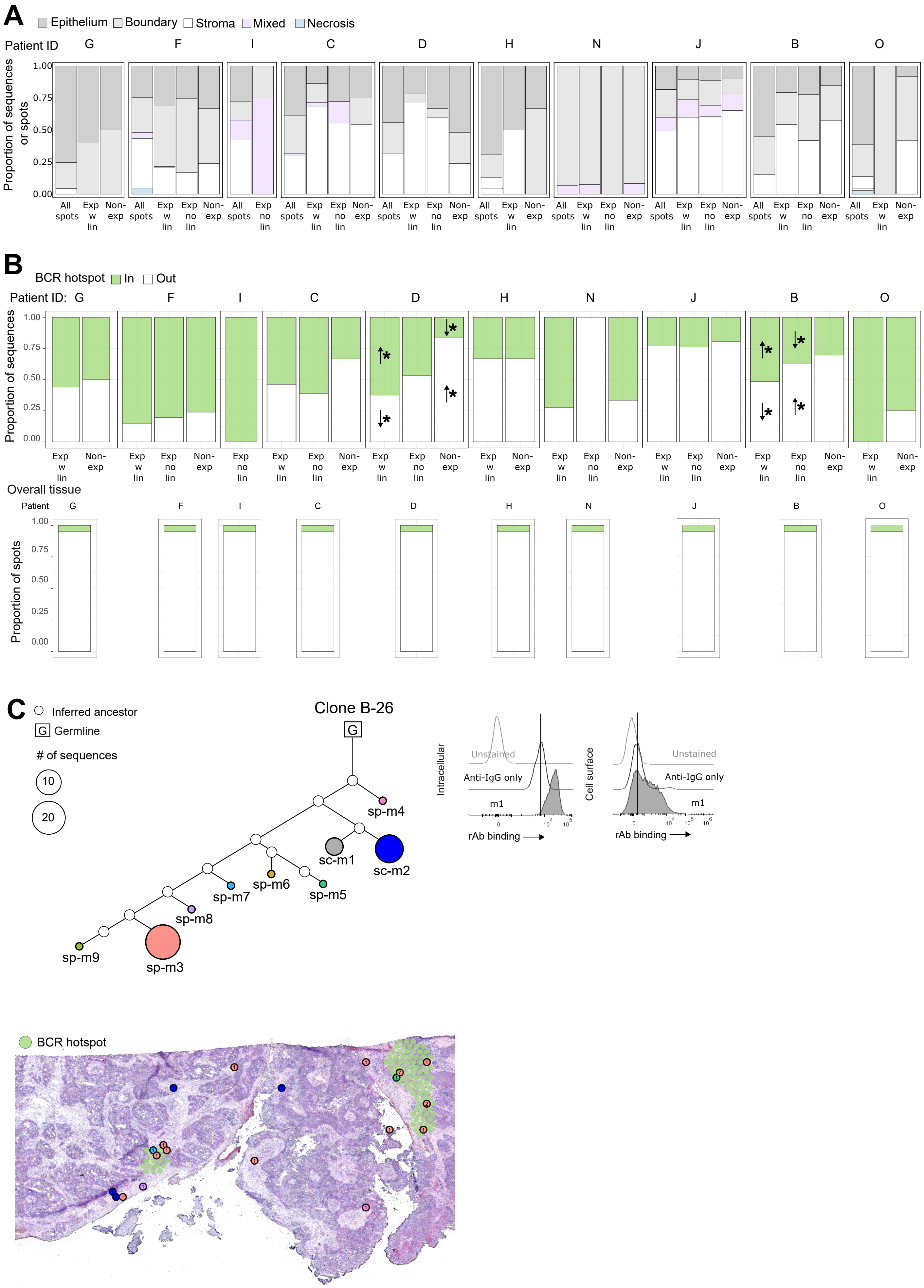
